# Visualizing PIEZO1 Localization and Activity in hiPSC-Derived Single Cells and Organoids with HaloTag Technology

**DOI:** 10.1101/2023.12.22.573117

**Authors:** Gabriella A. Bertaccini, Ignasi Casanellas, Elizabeth L. Evans, Jamison L. Nourse, George D. Dickinson, Gaoxiang Liu, Sayan Seal, Alan T. Ly, Jesse R. Holt, Tharaka D. Wijerathne, Shijun Yan, Elliot E. Hui, Jerome J. Lacroix, Mitradas M. Panicker, Srigokul Upadhyayula, Ian Parker, Medha M. Pathak

## Abstract

PIEZO1 is critical to numerous physiological processes, transducing diverse mechanical stimuli into electrical and chemical signals. Recent studies underscore the importance of visualizing endogenous PIEZO1 activity and localization to understand its functional roles. To enable physiologically and clinically relevant studies on human PIEZO1, we genetically engineered human induced pluripotent stem cells (hiPSCs) to express a HaloTag fused to endogenous PIEZO1. Combined with advanced imaging, our chemogenetic platform allows precise visualization of PIEZO1 localization dynamics in various cell types. Furthermore, the PIEZO1-HaloTag hiPSC technology facilitates the non-invasive monitoring of channel activity across diverse cell types using Ca^2+^-sensitive HaloTag ligands, achieving temporal resolution approaching that of patch clamp electrophysiology. Finally, we used lightsheet imaging of hiPSC-derived neural organoids to achieve molecular scale imaging of PIEZO1 in three-dimensional tissue organoids. Our advances offer a novel platform for studying PIEZO1 mechanotransduction in human cells and tissues, with potential for elucidating disease mechanisms and targeted therapeutic development.

## Introduction

PIEZO channels are pivotal in transducing mechanical stimuli into electrical and chemical signals, and play a significant role in a wide range of physiological functions^1,2^. PIEZO1, in particular, is expressed across various excitable and non-excitable tissues, shaping key biological processes such as vascular development^3,4^, exercise physiology^5^, blood pressure regulation^6–8^, red blood cell volume regulation^9,10^, neural stem cell differentiation^11,12^, and wound healing^13,14^. The channel has been linked to several human diseases, including hereditary xerocytosis^15–18^, lymphatic dysplasia^19,20^, iron overload^21^, and malaria^22^, and ongoing studies on this recently-identified protein are likely to uncover more disease associations.

So far, the study of PIEZO1 channel function has primarily utilized patch clamp electrophysiology^1,23–25^ and, more recently, measurements of Ca^2+^ influx through the channel^11,26–30^. These methodologies have provided valuable insights into the biophysical properties of PIEZO1. However, modulation of PIEZO1 function is complex, and the nature of PIEZO1 activity, as well as its downstream outcomes, are highly context-dependent^31^. Thus, to fully understand how PIEZO1 orchestrates diverse physiological roles and how its malfunction leads to diseases, it is crucial to study endogenous PIEZO1 in its native cellular environment. An emerging theme in PIEZO1’s physiological roles is its increased localization and activity at specific cellular structures, including focal adhesions^30,32–36; 30,32–36^, nuclei^30,32–36;30,32–36^, and cell-cell junctions^30,32–36^, suggesting a subcellular spatial organization of PIEZO1-mediated Ca^2+^ signals. Notably, the dynamic spatial positioning of PIEZO1 in migrating keratinocytes is a key determinant of their wound-healing capabilities^13^, highlighting the importance of PIEZO1 spatiotemporal dynamics in governing physiological processes. Taken together, these findings highlight the need for new methodologies to precisely and non-invasively monitor the spatiotemporal organization and activity of endogenous PIEZO1.

Current methods for visualizing PIEZO1 localization predominantly utilize the fusion of fluorescent proteins such as GFP^37,38^ or tdTomato^3,30,39^, but these methods are hampered by dimness, photobleaching, and an inability to measure channel activity. For measuring PIEZO1 activity, patch clamp electrophysiology is the standard assay^1,23,40^, but it disrupts cellular mechanics and offers limited insights into the channel’s spatial localization. In contrast, PIEZO1 activity measurements using cytosolic Ca^2+^-sensitive indicators in native cells, though spatially informative, lack specificity and can be confounded by Ca^2+^ flux through other channels. Additionally, non-human model organisms commonly used to explore the physiological roles of PIEZO1 may not fully recapitulate the channel’s behavior in human physiology^41,42^.

Here we present a novel platform to overcome these challenges and to provide a versatile human-specific system to complement animal studies, thus advancing physiologically- and clinically-relevant research on human PIEZO1. Utilizing CRISPR engineering, we introduced a self-labeling HaloTag domain fused to endogenous PIEZO1 in human induced pluripotent stem cells (hiPSCs), which can be differentiated into a variety of specialized cells and tissue organoids. Combined with the use of bright and photostable Janelia Fluor (JF)-based HaloTag ligands (HTLs)^43–45^, super-resolution imaging approaches, and automated image analysis pipelines, our platform allows the study of endogenous human PIEZO1 through imaging assays in various hiPSC-derived cell types and in vitro tissue organoids. This advance not only enables quantitative imaging of PIEZO1 channel localization and activity across diverse physiological scenarios but also enables human disease modeling of PIEZO1. Our work opens avenues towards mechanistic studies of endogenous human PIEZO1 in a variety of cell types, disease conditions as well as large-format drug screening for the development of targeted therapeutics.

## Results

### Development and validation of PIEZO1-HaloTag hiPSC lines

To create a novel, multifaceted tool for visualizing endogenous PIEZO1, we utilized CRISPR engineering to tag the endogenous channel with HaloTag, a modified bacterial haloalkane dehydrogenase that covalently binds to exogenously-provided chloroalkane HaloTag ligands^46^. We performed the genetic edits at both copies of PIEZO1 in hiPSCs, which are capable of self-renewal as well as differentiation into a variety of specialized cell types and tissue organoids (Fig. 1A). We attached the HaloTag to the C-terminus of PIEZO1 (Fig. 1B), as this location has previously been used for endogenous PIEZO1 tagging without affecting channel function^3^. The PIEZO1-HaloTag hiPSCs, as well as their differentiated progeny, are expected to express the HaloTag protein fused to endogenous PIEZO1, enabling covalent labeling with cognate HaloTag ligands in a variety of cell types.

**Figure 1.**
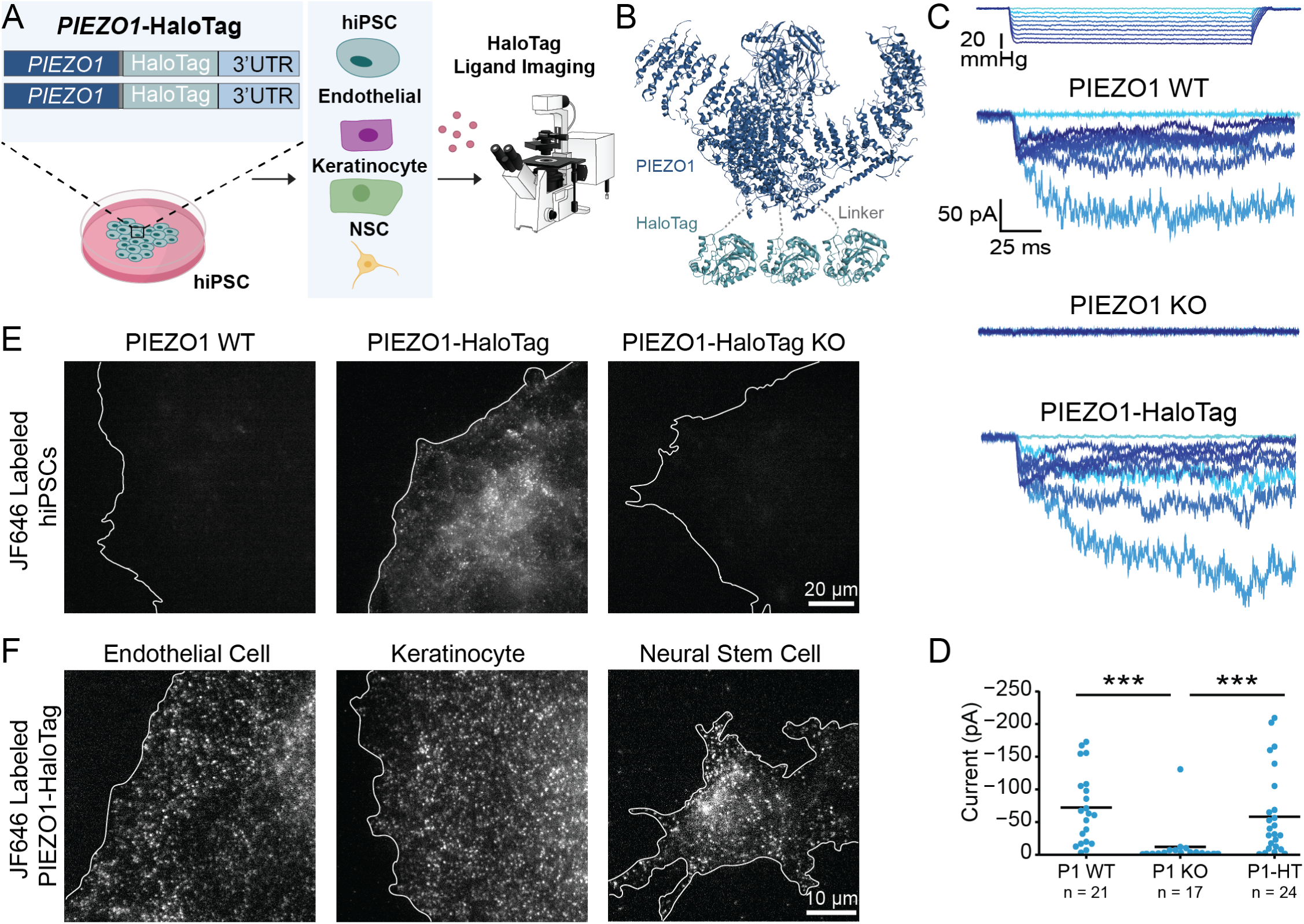
Generation and validation of the PIEZO1-HaloTag hiPSC line. **A.** Flowchart illustrating PIEZO1-HaloTag CRISPR knock-in in WTC-11 hiPSCs; multiple human cell types differentiated from the PIEZO1-HaloTag hiPSC line; and subsequent HaloTag-ligand probe labeling and imaging. **B.** Structural schematic of the trimeric PIEZO1 channel (Dark blue, PDB: 5Z10) with HaloTag (Cyan, PDB: 5UY1) attached to the cytosolic C-terminus. Dashed grey lines highlight the linker sequence (G-S-G-A-G-A) between PIEZO1 and HaloTag. **C.** Representative traces of cell-attached patch clamp measurements with mechanical stimulation imparted through negative suction pulses for endothelial cells derived from WTC-11, PIEZO1 KO, and PIEZO1-HaloTag hiPSCs. Blue color gradient indicates strength of negative pressure steps associated with suction pulses (light blue: lowest pressure, darkest blue: highest pressure). **D.** Maximal suction-evoked current amplitudes recorded in each condition from 5 independent experiments. All values are expressed as mean ± SEM (WTC-11 mean: -71 ± 12.1 pA, n = 21; PIEZO1 KO mean: -10.6 ± 7.5 pA, n = 17; PIEZO1-HaloTag mean: -60.8 ± 13.4 pA, n = 24) (*** p-value < 0.005). Cohen’s d effect sizes are -1.31 for PIEZO1 KO and as compared to WT. WTC-11 and PIEZO1-HaloTag did not show a statistically significant difference (p-value = 0.58) **E.** TIRF images, representative of 3 independent experiments, of a wild-type WTC-11 hiPSC without the Halo Tag introduced (left); PIEZO1-HaloTag hiPSC labeled with JF646 HTL (middle); and a PIEZO1-HaloTag Knockout hiPSC (right) showing lack of punctate labeling with the HaloTag ligand. **F.** TIRF images, representative of 3 independent experiments, of differentiated PIEZO1-HaloTag endothelial cell, keratinocyte, and neural stem cell labeled with JF646 HTL.See also Supplemental Figs. 1, 2, and 3 and Supplemental Videos 1, 2, and 3.

We first confirmed the effective generation of the PIEZO1-HaloTag fusion protein by western blot (Supplemental Fig. 1). Using an anti-PIEZO1 antibody, we observed a band in whole-cell lysate from the parent WTC-11 hiPSCs at the expected size of approximately 289 kDa for PIEZO1 and from PIEZO1-HaloTag hiPSCs at approximately 319 kDa. This increase in mass is consistent with fusion of HaloTag (33 kDa) to PIEZO1. As an orthogonal assay, we performed western blotting used an anti-HaloTag antibody and observed a signal in the PIEZO1-HaloTag hiPSC line, also at approximately 319 kDa, matching the mass observed with the anti-PIEZO1 antibody. This 319 kDa protein band was absent in the parent WTC-11 line. To further confirm HaloTag incorporation to PIEZO1, we knocked out PIEZO1 in the engineered PIEZO1-HaloTag hiPSCs. The PIEZO1-HaloTag Knockout hiPSCs did not exhibit a band using either the anti-PIEZO1 or the anti-HaloTag antibody. Taken together, we conclude that the heavier band corresponding to 319 kDa represents the endogenous PIEZO1 channel fused to the HaloTag domain and that the genetic modification results in the tagging of all PIEZO1 protein produced by the cell.

To determine whether the fusion of the HaloTag to PIEZO1 affected the channel’s function, we evaluated PIEZO1 ionic currents by cell-attached patch clamp electrophysiology using hiPSC-derived endothelial cells, which show high expression of PIEZO1^3,4^. Cell-attached patch clamp measurements, with mechanical stimulation of the membrane patch through negative pressure pulses, revealed mechanically-evoked currents with slow inactivation and deactivation as previously demonstrated in primary endothelial cells^47^. These currents were absent in endothelial cells differentiated from PIEZO1-Knockout WTC-11 hiPSCs, confirming that they represent ionic current through PIEZO1 (Fig. 1C). Notably, maximal currents from endothelial cells differentiated from the PIEZO1-HaloTag hiPSCs were similar in magnitude and indistinguishable to those from endothelial cells generated from the parent hiPSC line indicating that the HaloTag fusion did not abrogate channel activation (Fig. 1D).

To test labeling specificity, we incubated hiPSCs with the Janelia Fluor 646 HaloTag ligand^45^ (JF646 HTL, see Methods) and imaged the cells with Total Internal Reflection Fluorescence (TIRF) microscopy (Fig. 1E). The PIEZO1-HaloTag hiPSCs displayed a punctate signal, as previously observed for endogenous, tagged PIEZO1-td-Tomato channels^30^. In contrast, PIEZO1-HaloTag Knockout hiPSCs and wild-type WTC-11 hiPSCs both showed little or no punctate staining. To confirm the specific labeling of PIEZO1-HaloTag channels in the differentiated progeny of PIEZO1-HaloTag hiPSCs, we differentiated PIEZO1-HaloTag hiPSCs into three cell types which are known to express PIEZO1: endothelial cells (ECs) ^3,4,39^, keratinocytes^13^, and neural stem cells (NSCs) ^30^ (Fig. 1F, Supplemental Fig. 2). Upon labeling with the JF646 HTL, each differentiated cell type exhibited a punctate signal (Fig. 1F, Supplemental Videos 1, 2, and 3), whereas cells differentiated from PIEZO1-HaloTag Knockout hiPSCs lacked these punctate signals (Supplemental Fig. 3). To determine whether the attachment of the HaloTag ligand affected channel activity, we performed whole-cell patch clamp on PIEZO1-HaloTag ECs and NSCs with mechanical stimulus imparted with poking using a blunt glass probe (poking assay). We compared unlabeled and HaloTag ligand-labeled samples from each cell type, and PIEZO1-HaloTag KO cells were again used as controls. The labeled cells showed no changes in inactivation or deactivation kinetics, and no change in inactivation or deactivation maximal current amplitude, compared to unlabeled PIEZO1-HaloTag cells, indicating that the labeled PIEZO1-HaloTag channel functions similarly to the wild-type channel (Supplemental Fig. 4).

Overall, we validated the PIEZO1-HaloTag hiPSC line as a method to attach HaloTag ligands specifically to human PIEZO1 in multiple cell types while preserving PIEZO1 function.

### PIEZO1-HaloTag imaging with high signal-to-noise and reduced photobleaching

PIEZO1 has been shown to be mobile in the plasma membrane^30,32,38,39^ and we confirmed this in hiP-SC-derived neural stem cells, keratinocytes, and endothelial cells differentiated from PIEZO1-HaloTag hiPSCs and labeled with JF646 HTL (Supplemental Videos 1, 2, and 3). Previous studies on the mobility of endogenous PIEZO1 have utilized cells harvested from a PIEZO1-tdTomato knock-in mouse^30,39^ where the tdTomato fluorescence is limited by rapid photobleaching and low signal-to-noise ratio. We compared the spatial and temporal resolution of data obtained from PIEZO1-tdTomato endothelial cells harvested from PIEZO1-tdTomato mice and from PIEZO1-HaloTag hiPSC-derived endothelial cells (Fig. 2A).

**Figure 2.**
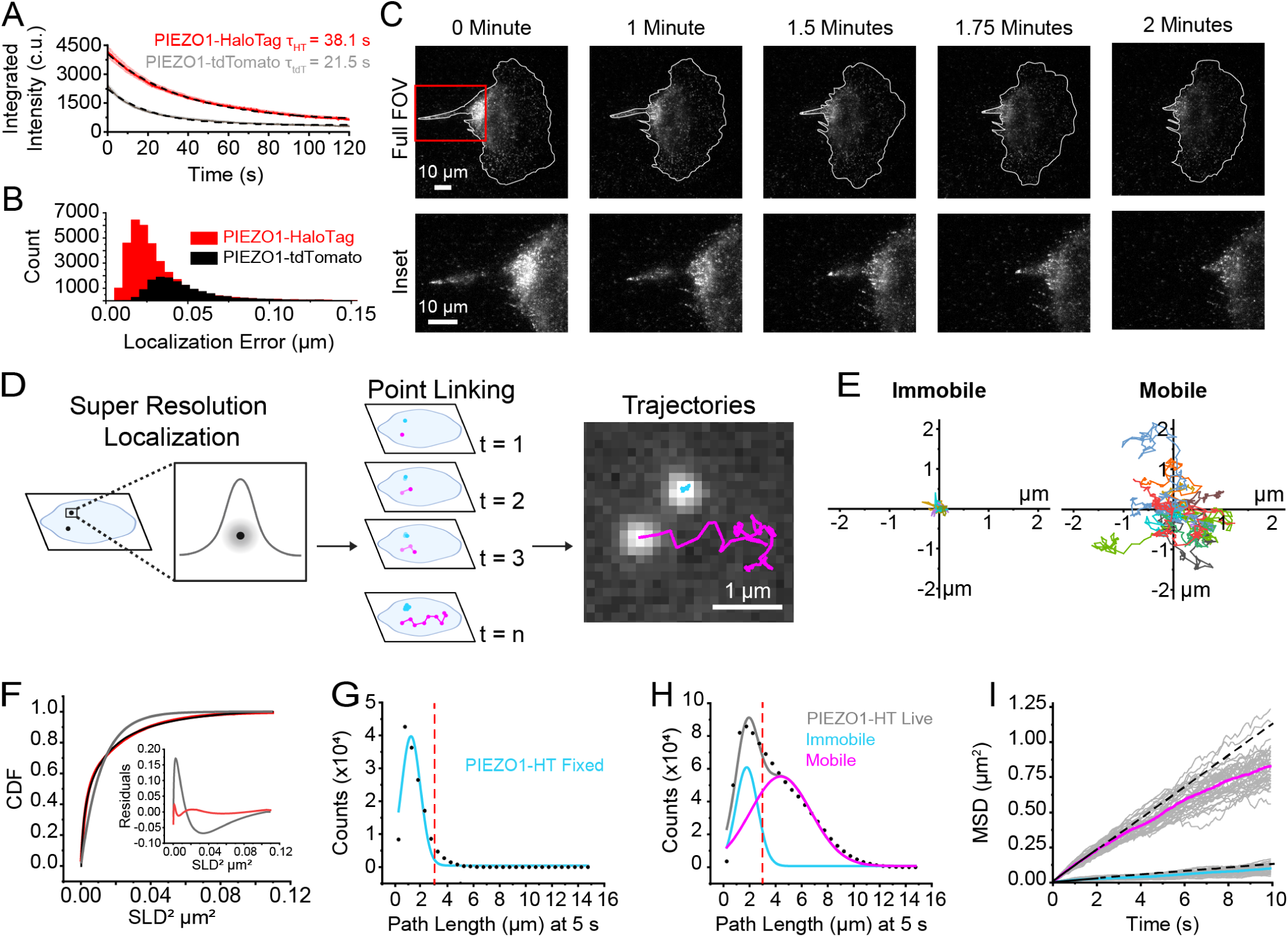
PIEZO1-HaloTag localization and tracking reveal populations of PIEZO1 with different motilities. **A.** PIEZO1-HaloTag puncta are brighter and bleach slower than PIEZO1-tdTomato puncta. TIRF image series of PIEZO1-HaloTag and PIEZO1-tdTomato endothelial cells were acquired for 2 minutes with identical acquisition settings (see Methods). Red trace indicates the average integrated intensity of PIEZO1-HaloTag (n = 19 videos from 3 independent experiments) and grey trace indicates the average integrated intensity of PIEZO1-tdTomato (n = 20 videos from 3 independent experiments). The signals are scaled to the number of puncta in the first frame for each video. Black dashed curves represent exponential fits to the data with τ_HT_ = 38.1 ± 0.15 s and τ_tdT_ = 21.5 ± 0.13 s (p-value Mann-Whitney < 0.005 and the Cohen’s d value= -2.73). **B.** Distributions of localization errors, calculated as the deviation from the mean position of a trajectory, for trajectories extracted from PIEZO1-HaloTag endothelial cells labeled with JF549 HTL and PIEZO1-tdTomato mouse embryonic fibroblasts. Cells in each condition were imaged with identical acquisition settings. **C.** A migrating PIEZO1-HaloTag NSC imaged for 2 minutes with a frame rate of 10 fps. The cell is labeled with JF646 HTL. Top row shows the entire cell (Full FOV) and the bottom row shows a zoom-in of the inset box (red) on the rear of the cell. Note the channel enrichment at the rear of the cell throughout the 2-minute recording. **D.** Schematic depicting super-resolution localization, point linking, and tracking of PIEZO1-HaloTag channels labeled with JF646 HTL. Examples show two observed puncta motility behaviors: mobile (magenta) and immobile (cyan). **E.** Representative trajectories from 11 immobile and 11 mobile puncta. Starting positions of trajectories are scaled to the origin, with different trajectories indicated by color. **F.** Cumulative distribution functions (CDF) of Single Lag Displacements (SLD). Black dotted curve shows experimental data. Grey curve is a single-component exponential fit. Red curve is a two-component exponential fit. Residuals for single-component (grey) and two-component (red) fits are shown in the inset plot. **G.** Distribution of trajectory path lengths at 5 s (50 frames) derived from PIEZO1-HaloTag endothelial cells labeled with JF646 HTL imaged after fixation (number of tracks = 14,943 from 13 videos). Dashed red line indicates the 3 µm cutoff criterion used to define immobile trajectories. **H.** Corresponding distribution of trajectory path lengths for tracks at 5 s derived from live PIEZO1-HaloTag endothelial cells labeled with JF646 HTL (number of tracks = 889,971 from 40 videos). Grey curve represents a fit to a sum of two Gaussian curves; the individual Gaussian curves are shown in cyan and magenta. Dashed red line indicates cutoff to discriminate mobile and immobile trajectories **I.** Mean squared displacement (MSD) of immobile (cyan) and mobile puncta (magenta). Tracks that had a path length > 3 µm were characterized as mobile (magenta) and tracks which had a path length < 3 µm were characterized as immobile (cyan) in the Gaussian curve. Each gray trace represents mean MSD for all mobile puncta in a video (upper traces) and for all immobile puncta in a video (lower traces); data from 39 videos are plotted. Solid magenta (mobile) and cyan (immobile) curves are mean MSD curves across all videos. Solid black line is a linear fit to the initial MSD t < 2 s, dashed black line shows a linear extrapolation. All values are expressed as mean ± SEM. Data for panels E-I are from 4 independent experiments. See also Supplemental Video 4.

We labeled PIEZO1-HaloTag endothelial cells with Janelia Fluor 549 HaloTag ligand (JF549 HTL), enabling consistent experimental settings; i.e. the same 561 nm laser wavelength and power settings, filters, and camera acquisition settings as for tdTomato. Under the imaging conditions used, the initial integrated intensity shown in camera units (c.u.) (see Methods) of JF549 HTL-labeled PIEZO1-HaloTag puncta (4179 ± 268.8 c.u.) was approximately twice that of the PIEZO1-tdTomato puncta (2420 ± 108.6 c.u.), and they bleached more slowly, (time constant τ= 38.1 ± 0.15 s for PIEZO1-HaloTag vs. τ= 21.5 ± 0.13 s for PIEZO1-tdTomato puncta (Fig. 2A). Here and throughout all means are shown as ± SEM. The mean signal-to-background ratio was 5.62 ± 0.19 for PIEZO1-HaloTag puncta as compared to 3.14 ± 0.12 for PIEZO1-tdTomato puncta. Furthermore, the localization error, assessed by measuring frame-by-frame deviations from the mean position of puncta in fixed cells, was smaller for JF549 HTL-labeled PIEZO1-HaloTag channels (33 ± < 1 nm) compared to fixed PIEZO1-tdTomato channels (52 ± < 1 nm) (Fig. 2B). JF646 HTL labeled PIEZO1-HaloTag puncta had a localization error of 29 ± < 1 nm. Within our fixed sample data, we found a negligible contribution of x-y drift (<10 nm over 5 s). Thus, PIEZO1-HaloTag puncta were brighter, bleached more slowly, and enabled greater localization precision compared to PIEZO1-tdTomato puncta.

To illustrate the improved imaging capability of the PIEZO1-HaloTag system, we used TIRF microscopy to monitor a migrating PIEZO1-HaloTag neural stem cell labeled with JF646 HTL (Fig. 2C, Supplemental Video 4). We have previously shown that PIEZO1 is enriched at the rear of migratory cells; however, rapid photobleaching of the PIEZO1-tdTomato fluorophore precluded fast imaging of PIEZO1 dynamics during cell migration^13^. By imaging PIEZO1-HaloTag channels in a migrating neural stem cell over 2 minutes at a temporal resolution of 10 frames per second (fps) (Fig. 2C, Supplemental Video 4), we observed PIEZO1 channel enrichment at the trailing end of the cell throughout rear retraction, with a subset of PIEZO1 puncta displaying linear organization at the back of the cells. These observations highlight the enhanced imaging capacity of the PIEZO1-HaloTag system, allowing for high resolution visualization of PIEZO1 dynamics and organization during cellular processes.

### Super-resolution tracking of PIEZO1 puncta reveals distinct mobility modes

To analyze the mobility of PIEZO1 in the membrane, we used the FIJI plugin ThunderSTORM ^48,49^ for super-resolution localization of PIEZO1-HaloTag puncta in TIRF image stacks. Successive localizations were linked to form trajectories using the custom-built, open-source image processing software FLIKA ^50^ (Fig. 2D, Supplemental Video 5, see Methods). We compared the number of tracked puncta detected from PIEZO1-HaloTag knockout endothelial cells to PIEZO1-HaloTag endothelial cells, both labeled with JF646 HaloTag ligand. PIEZO1-HaloTag cells yielded 0.5 ± 0.05 tracked puncta/µm² while knockout cells yielded 0.004 ± 0.0006 tracked puncta/µm² (*** p < 0.005, Cohen’s d = -5.44), indicating that our analysis methods yielded minimal spurious trajectories. Visual inspection of PIEZO1-HaloTag trajectories revealed the presence of two distinct populations of puncta: those that appeared immobile and others that were highly motile (Fig. 2E, Supplemental Video 4), concordant with recent findings by Ly et al^39^. Trajectories superimposed on a fluorescent image of the endoplasmic reticulum (ER), showed immobile and mobile puncta, both coincident with the ER signal and in regions without ER (Supplemental Fig. 5). Thus, it appears that the motility of puncta is not exclusively governed by intracellular organelles like the ER.

We calculated Single Lag Displacements (SLD), i.e the distance covered by a punctum between consecutive frames, and generated a cumulative distribution function (CDF) of SLD^2^ from the individual PIEZO1 trajectories (Fig. 2F). In a population with a single diffusive behavior, the CDF should fit a single exponential function. However, our data was not adequately fit by a single-component exponential, whereas a two-component exponential fit well (Fig. 2F), consistent with the presence of two distinct motility behaviors. To segregate the two populations we separated the tracks of individual puncta based on the total path length traveled within a period of 5 seconds. We first determined how much apparent movement would result from localization error by labeling PIEZO1-HaloTag endothelial cells with JF646 HTL and imaging them after fixing the samples. The apparent path lengths over 5 s, representing a summation of localization errors over 50 frames, for these fixed puncta followed a roughly Gaussian distribution peaking at about 1.20 µm (Fig. 2G). In contrast, trajectories from live PIEZO1-HaloTag endothelial cells labeled with the same probe showed a distribution of path lengths that was fitted well by a two-component Gaussian distribution (Fig. 2H). The first component of the Gaussian fit (peak 1.78 µm) approximated that of the fixed cell data, suggesting that this fraction of PIEZO1 puncta is indeed almost immotile in live cells. To extract the second, mobile component, we selected a cutoff value of path length of >3 µm at 5 s to largely exclude ‘immotile’ puncta (see dashed red lines in Fig. 2G and Fig. 2H). Fig. 2I shows plots of the mean squared displacements (MSD) vs. time for classes of immotile and motile puncta as segregated by this criterion. Traces for these populations separated into two distinct groups (Fig. 2I). Puncta undergoing Brownian (random) diffusion would display a straight line on this plot, with the diffusion coefficient D = d^2^/4t, where d is the mean distance from origin at time t. A linear fit up to 2 s for the motile population yielded an apparent diffusion coefficient of 0.029 μm^2^ s^−1^, representing the upper limit of the diffusion coefficient. At longer times the relationship fell below linear (Fig. 2I), indicating sub-Brownian, anomalous diffusion, similar to our recent observations with PIEZO1-tdTomato channels^39^. In contrast, the diffusion coefficient for the immotile population based on the linear fit up to 2 s yielded a maximal diffusion coefficient of 0.003 μm^2^ s^−1^. Using the same path length cutoff value in PIEZO1-HaloTag neural stem cells also yielded two distinct populations of mobile and immobile PIEZO1 puncta (Supplemental Fig. 6). The diffusion coefficients for mobile PIEZO1 puncta (0.029 μm2 s^-1^ in endothelial cells and 0.026 μm2 s^-1^ in neural stem cells) fall within the expected diffusion range for large, plasma membrane ion channels as the diffusion coefficients for IP3R (0.064 μm2 s^-1^)^51^, TRAAK in a freestanding bilayer (0.8 μm2 s^-1^)^53^, and PIEZO1 in red blood cells (0.037 μm2 s^-1^)^52^ are similar to our reported values.

### Monitoring PIEZO1 activity with a Ca^2+^-sensitive HaloTag ligand

The availability of Ca^2+^-sensitive HTLs ^44^ enables measurement of channel activity using the PIEZO1-HaloT-ag system. We labeled PIEZO1-HaloTag endothelial cells with Janelia Fluor 646-BAPTA HaloTag Ligand (JF646-BAPTA HTL), a non-ratiometric Ca^2+^-sensitive fluorescent indicator ^44^. The location of the HaloTag at the C-terminus, near the cytosolic pore region of the channel (Fig. 1B), optimally places the probe to report on instantaneous Ca^2+^ influx as increases in fluorescence intensity (“flickers”).

To evaluate the Ca^2+^ sensitivity of the probe we transfected monomeric HaloTag into WTC-11 hiPSCs and labeled the cells with a 1:1 mixture of the Ca^2+^-sensitive HTL JF646-BAPTA and the non-Ca^2+^-sensitive JF549 (Supplemental Fig. 7). After cell fixation and permeabilization, we performed TIRF microscopy over a range of free Ca^2+^ concentrations (0-39 μM). In the absence of Ca^2+^, JF646-BAPTA-labeled puncta displayed barely detectable fluorescence, whereas JF549-labeled puncta showed stable fluorescence with little flickering (Supplemental Fig. 7). At a free Ca^2+^ concentration of 75 nM, JF646-BAPTA-labeled puncta exhibited increased flickering to a bright state, and at 39 µM showed largely persistent bright fluorescence (Supplemental Fig. 7). Given the high local Ca^2+^ concentration (> 15 µM) expected close to the pore of an open channel^54^, signals from JF646-BAPTA HTL PIEZO puncta would reflect channel gating, rather than random binding and unbinding of Ca^2+^ ions to the HTL.

We next imaged live ECs labeled with JF646-BAPTA HTL in a bath solution containing 3 mM Ca^2+^. We observed a relatively sparse density of puncta (0.07 ± 0.004 puncta per µm^2^) as compared to cells labeled with the non Ca^2+^-sensitive JF646 HTL (0.34 ± 0.01 puncta per µm^2^) (Fig. 3A). Endothelial cells derived from the PIEZO1-HaloTag Knockout hiPSCs had an even lower puncta density of 0.02 ± 0.002 per µm^2^. Taken together, these observations suggest that a small fraction of PIEZO1 puncta are active in unstimulated cells. To verify that detected JF646-BAPTA HTL signals reflect PIEZO1 activity, we applied 2 µM Yoda1, a chemical agonist of the channel that increases the open-state occupancy of PIEZO1^28,55^. Incubating PIEZO1-HaloTag endothelial cells with Yoda1 substantially increased the density of puncta to 0.22 ± 0.02 puncta per µm^2^ (Fig. 3A and B, Supplemental Video 6).

**Figure 3.**
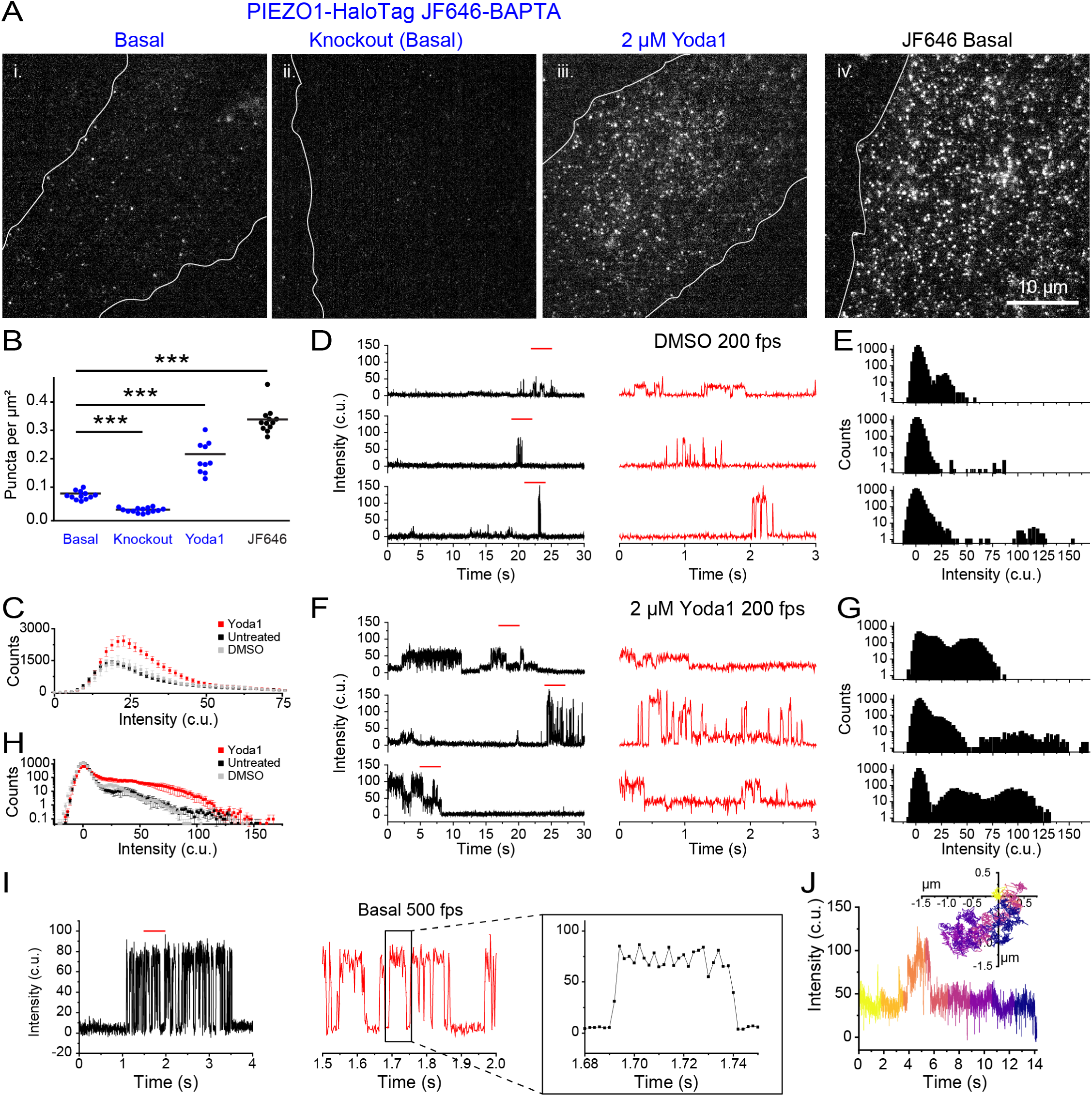
PIEZO1-HaloTag enables imaging of endogenous PIEZO1 activity with temporal resolution approaching that of patch-clamp electrophysiology. **A**. TIRF images of endothelial cells differentiated from i. PIEZO1-HaloTag and ii. PIEZO1-HaloTag Knockout lines. iii. PIEZO1-HaloTag treated with 2 µM Yoda1 samples. i-iii are all labeled with the Ca^2+^-sensitive HaloTag Ligand JF646-BAPTA HTL. iv. shows a PIEZO1-HaloTag endothelial cell labeled with the non-Ca^2+^-sensitive HaloTag Ligand JF646 HTL. **B.** Puncta densities with labeling by JF646-BAPTA HTL (blue) and JF646 (black). All values are expressed as mean ± SEM density of puncta per µm^2^. JF646-BAPTA HTL; Basal, 0.07 ± 0.004; PIEZO1-HaloTag KO, 0.02 ± 0.002; 2 µM Yoda 1, 0.22± 0.02; JF646 HTL, 0.34± 0.01. Data are from 4 independent experiments. All groups were significantly different from one another (*** p-value Mann-Whitney < 0.005 for all conditions). Cohen’s d effect sizes of PIEZO1-HaloTag treated with 2 µM Yoda1 compared to PIEZO1-HaloTag (2.28) and of PIEZO1-HaloTag Knockout compared to PIEZO1-HaloTag (-4.46). **C.** Average histogram of JF646-BAPTA puncta intensity in the bright state across every frame in 32 untreated, 12 DMSO, and 13 2 µM Yoda1 videos from 3 independent experiments. **D.** Representative background-subtracted fluorescence intensity traces of immobile PIEZO1-HaloTag puncta from TIRF imaging of hiPSC-derived endothelial cells labeled with the Ca^2+^-sensitive HaloTag Ligand JF646-BAPTA HTL and treated with vehicle control, DMSO. Right, expanded traces corresponding to the red line marked on the left. **E.** All-points amplitude histograms of intensity levels from the 30 s traces in C. Counts are shown on a log10 scale. **F.** Representative traces, as in C, of puncta from PIEZO1-HaloTag endothelial cells treated with 2 µM Yoda1. **G.** An all-points amplitude histogram of intensities from the 30 s traces in F. **H.** An average all-points amplitude histogram (i.e. including dark and bright states) from 21 immobile puncta representing 3 independent experiments of each condition (untreated, DMSO-treated or treated with 2 µM Yoda1) of PIEZO1-HaloTag JF646-BAPTA labeled ECs. The Yoda1 distribution was significantly different from both the Untreated and DMSO distributions, using a Two-Sample Kolmogorov-Smirnov test (*p* < 0.001). For individual fluorescence traces and all-points amplitude histogram of each punctum, please see Supplemental Fig. 8-10. **I.** Representative trace of a JF646-BAPTA HTL labeled immobile PIEZO1-HaloTag punctum imaged using TIRF at a frame rate of 500 fps. Other details are the same as in panels D and F. Note the improved temporal resolution of PIEZO1-mediated Ca2+ signals illustrated by the progressively expanded traces on the right. **J.** Representative trajectory and fluorescence intensity profile of a mobile punctum labeled with JF 646-BAPTA. The inset Cartesian coordinate plot illustrates the trajectory with the starting position normalized to the origin, and time depicted by progressively cooler colors. The graph shows the corresponding background-subtracted fluorescence intensity trace. Data for panels D - G are from 3 independent experiments. See also Supplemental Figs. 8, 9, and 10 and Supplemental Videos 7, 8, and 9.

### Monitoring activity of endogenous human PIEZO1 with high temporal resolution

To visualize channel activity dynamics, we imaged PIEZO1-HaloTag endothelial cells labeled with JF646-BAPTA HTL at 200 fps. The fluorescence intensity profiles of PIEZO1-HaloTag JF646-BAPTA puncta showed flickers above the baseline (Supplemental Video 7). We plotted puncta fluorescence intensity in the bright state across every frame for untreated cells, DMSO control, and 2 µM Yoda1-treated cells. The presence of Yoda1 resulted in both an increase in the number of bright-state puncta as well as an increase in their fluorescence intensity (untreated peak: 18.95 c.u, DMSO peak: 20.85 c.u., 2 µM Yoda1 peak: 22.75 c.u.) (Fig. 3C). To quantify fluctuations between the dim and bright state, we initially focused on immotile puncta for our analysis (Supplemental Videos 7-9). Flickers from individual puncta in untreated and vehicle control DMSO samples could be clearly resolved in fluorescence traces (Fig. 3D-G; Supplemental Videos 7, 9; Supplemental Figs. 8,9). All-points amplitude histograms showed the presence of multiple intensity levels (Fig. 3E-F; Supplemental Video 7, 9; Supplemental Figs. 8, 9). Between flickers, the signal at PIEZO1-HaloTag puncta sites was generally indistinguishable from the background fluorescence at surrounding regions, indicating that the intrinsic fluorescence of the BAPTA probe is very low at resting cytosolic [Ca^2+^]. The absolute fluorescence values corresponding to the peaks of each flicker level varied between different puncta (Fig. 3D, F; Supplemental Fig. 8-10) - possibly a result of different axial locations of the plasma membrane within the evanescent field. Treatment with Yoda1 increased the proportion of time that puncta exhibited increased fluorescence and increased the proportion of time at higher intensity levels (Fig. 3F-G; Supplemental Video 8; Supplemental Fig. 10).

We next plotted an average all-points amplitude histogram from untreated, DMSO-treated, and 2 µM Yoda1-treated immobile trajectories (Supplemental Fig. 8-10). The average all-points amplitude histogram from 21 puncta per condition showed that PIEZO1-HaloTag fluorescence had a dim state peak near 0 c.u. (background) across all conditions. The bright state intensity increased with Yoda1 treatment, consistent with its role in promoting PIEZO1 activation (Fig. 3H).

To further evaluate the temporal resolution of our recordings, we imaged JF646-BAPTA HTL signals in endothelial cells at a faster frame rate of 500 fps (Fig. 3I). Transitions on both the rising and falling phases of flickers were complete within 2 frames (Fig. 3I), indicating that the fluorescence recordings track channel gating with a time resolution of 4 ms or better.

Next, we extended our analysis to mobile JF646-BAPTA HTL-labeled puncta, achieving simultaneous detection of channel location and activity (Fig. 3J, Supplemental Video 9). However, a current limitation is that the dim basal fluorescence of the JF646-BAPTA HTL generally precluded the detection of a punctum when it was inactive, so tracks were restricted to durations when a punctum was in the bright state or could be interpolated across dim state gaps of a few frames (see Methods). Taken together with measurements from immobile puncta above, this demonstrates that both immobile and mobile PIEZO1 puncta can be active.

### Investigating cell type-specific PIEZO1 activity patterns

The functional roles of PIEZO1 in a given cell type may depend on the number and behavior of active channels specific to that particular lineage. To examine PIEZO1 behavior across two different cell types, we labeled ECs and NSCs with JF646-BAPTA HTL (Fig. 4A) and first quantified the density of active puncta in each (Fig. 4B). NSCs had close to half of the active puncta per unit area (mean = 0.13 ± 0.02 puncta per µm^2^) compared to ECs (mean = 0.23 ± 0.02 puncta per µm^2^). To assess whether cell type specific differences in PIEZO1 expression contribute to the decrease in the density of active NSC PIEZO1-HaloTag puncta, we labeled PIEZO-HaloTag ECs and NSCs with JF646 HTL. We noted that JF646 HTL-labeled NSCs had roughly 33% lower puncta density (mean = 0.30 ± 0.02 puncta per µm^2^) than ECs (mean = 0.45 ± 0.02 puncta per µm^2^) (Fig. 4C). However, this reduction in PIEZO1 expression was less pronounced than the nearly 50% decrease in PIEZO1 activity observed in NSCs compared to ECs, suggesting that our observed differences in active puncta density were at least partially, but not completely, due to different PIEZO1 expression levels in each cell type. We plotted the fluorescence intensity of JF646-BAPTA puncta in the bright state, showing that the peak intensity for ECs (17.05 c.u.) was higher than for NSCs (15.15 c.u.), with ECs also having more bright puncta at higher fluorescence intensity values (Fig. 4D). To better determine whether puncta in NSCs were less active, we imaged JF646-BAPTA-labeled cells at 500 fps and generated all-points amplitude histograms from these traces (Fig. 4E-F). We then plotted an average all-points amplitude histogram using 46 immobile puncta from ECs and NSCs each (Supplemental Figs. 11-12), which showed an increased proportion of counts EC PIEZO1 spends in the bright state relative to NSC PIEZO1 (Fig. 4G). Therefore, PIEZO1-HaloTag imaging revealed differences across the two cell types, with NSCs having fewer and less active puncta than ECs.

**Figure 4.**
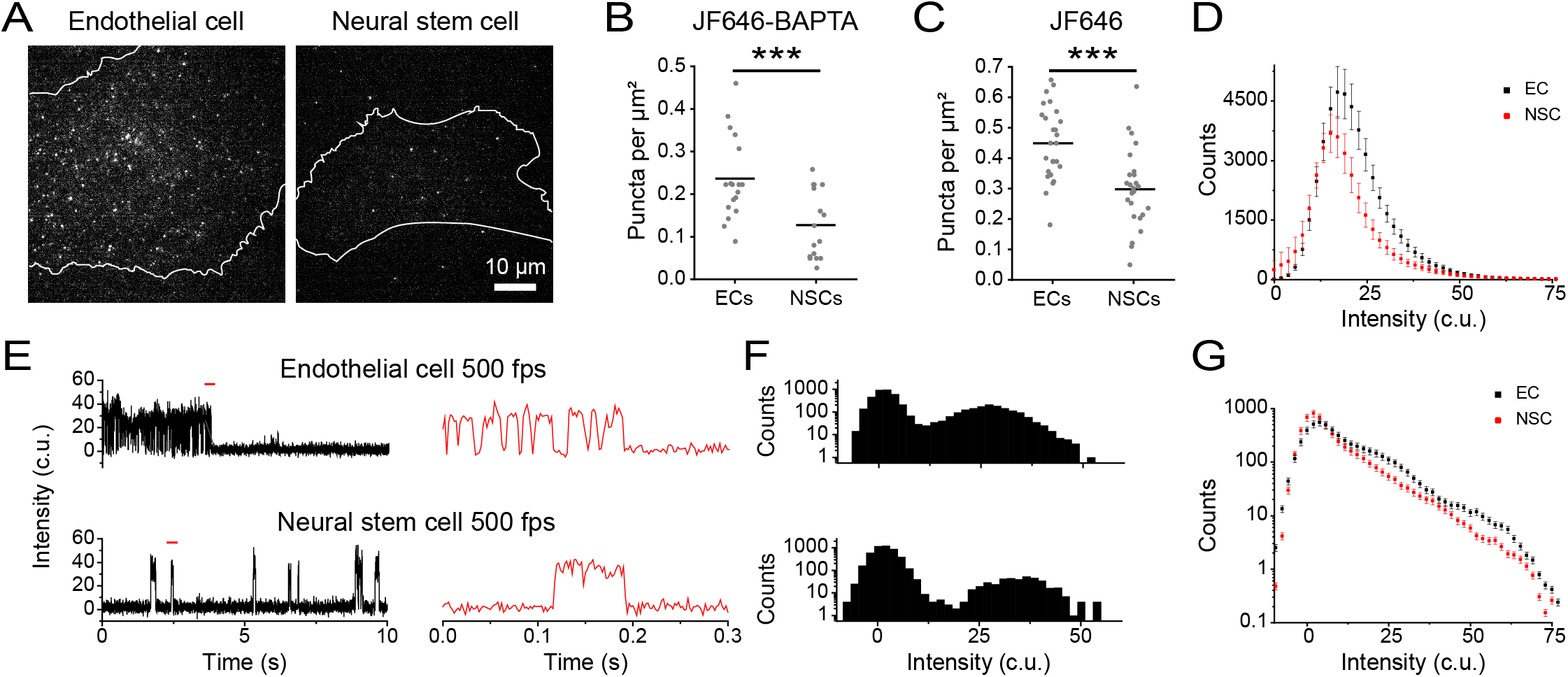
PIEZO1 activity monitored in endothelial and neural stem cells. **A.** Representative TIRF images of PIEZO1-HaloTag ECs and NSCs labeled with JF646-BAPTA HTL. **B.** JF646-BAPTA HTL puncta density in ECs (mean = 0.23 ± 0.02 puncta per µm^2^) and NSCs (mean = 0.13 ± 0.02 puncta per µm^2^). Each dot is representative of the puncta density in a cell (see Methods), n = 16 to 18 cells from 3 independent experiments. Means are indicated by black lines. The two groups were significantly different from one another, *** p-value Mann-Whitney < 0.005. The Cohen’s d effect size was -1.20. **C.** Same as B, for JF646 HTL puncta density in ECs (mean = 0.45 ± 0.02 puncta per µm2) and NSCs (mean = 0.30 ± 0.02 puncta per µm2), n = 26 cells from 3 independent experiments, *** p-value Mann-Whitney < 0.0001. The Cohen’s d effect size was -1.21. **D.** Average histogram of puncta intensity from all detected (i.e. bright state) JF646-BAPTA puncta across every frame in EC and NSC videos, recorded at 500 fps over 10 seconds, from 3 independent experiments. **E.** Representative background-subtracted fluorescence intensity traces of immotile PIEZO1-HaloTag puncta from 500-fps TIRF imaging of EC (top) and NSC (bottom) labeled with JF646-BAPTA HTL. *Right,* expanded traces from the sections marked with a red line on the left. **F.** Corresponding all-points amplitude histograms of intensity levels from the 10-s traces in E. Counts are shown on a log10 scale. **G.** An average all-points amplitude histogram from 46 immobile puncta representing 3 independent experiments of JF646-BAPTA-labeled PIEZO1-HaloTag ECs (black) and NSCs (red). The two distributions were significantly different from each other, using a Two-Sample Kolmogorov-Smirnov test (*p* < 0.01). For individual fluorescence traces and all-points amplitude histogram of each punctum, please see Supplemental Fig. 11-12.

### PIEZO1-HaloTag signals report on mechanically-evoked activity of PIEZO1

To assess the effectiveness of the PIEZO1-HaloTag system in reporting PIEZO1’s response to mechanical stimuli, we applied a hypotonic stimulus to PIEZO1-HaloTag ECs. The reduction in osmolarity is expected to increase membrane tension, thereby enhancing PIEZO1 activity ^56–58^. To evaluate the PIEZO1-HaloTag response to osmotic shock, we labeled PIEZO1-HaloTag ECs with JF646-BAPTA HTL and imaged them in solutions of different osmotic strengths: 314 mOsm/L control; 251 mOsm/L hypotonic (Supplemental Fig. 12A). We observed a 29% increase in the density of bright JF646-BAPTA-labeled puncta following the hypotonic stimulus (Supplemental Fig. 13B), increasing from 0.28 ± 0.02 puncta per µm^2^ in 314 mOsm/L to 0.36 ± 0.02 puncta per µm^2^ in 251 mOsm/L. We then imaged JF646-BAPTA-labeled cells in control and hypotonic solution at 200 fps and tracked puncta to obtain their intensity profiles and corresponding all-points amplitude histograms. Representative traces showed more channel activity in the hypotonic condition, which is also reflected in the all-points amplitude histogram (Supplemental Fig. 13C). We plotted an average all-points amplitude histogram of 71 tracked puncta in each condition (Supplemental Figs. 14-15), which showed that the hypotonic stimulus increased the proportion of PIEZO1 channels in the open state compared to the control solution (Supplemental Fig. 13D). Thus, the PIEZO1-HaloTag system can be used to study PIEZO1’s response to mechanical stress.

### Imaging PIEZO1 in a Tissue Organoid Model of Neural Development

The PIEZO1-HaloTag hiPSC line enables the investigation of PIEZO1 at the tissue scale using organoid models derived from hiPSCs. To demonstrate this, we examined PIEZO1 localization and activity in Micropatterned Neural Rosettes (MNRs)^59,60^, an hiPSC-derived human organoid model that mimics early neural development. MNRs exhibit a reproducible radial cell organization around a single central lumen, analogous to the cross-section of a neural tube (Fig. 5A). We generated MNRs from PIEZO1-HaloTag hiPSCs, labeled them with JF646 HTL, fixed the samples, and imaged them using confocal microscopy. Co-staining with SPY555-Actin revealed an actin-rich lumen and outer edge, with radial actin signal between these two regions. Punctate HTL signal was located at cell-cell interfaces, and was more pronounced closer to the central lumen than at the MNR’s outer edge (Fig. 5B).

**Figure 5.**
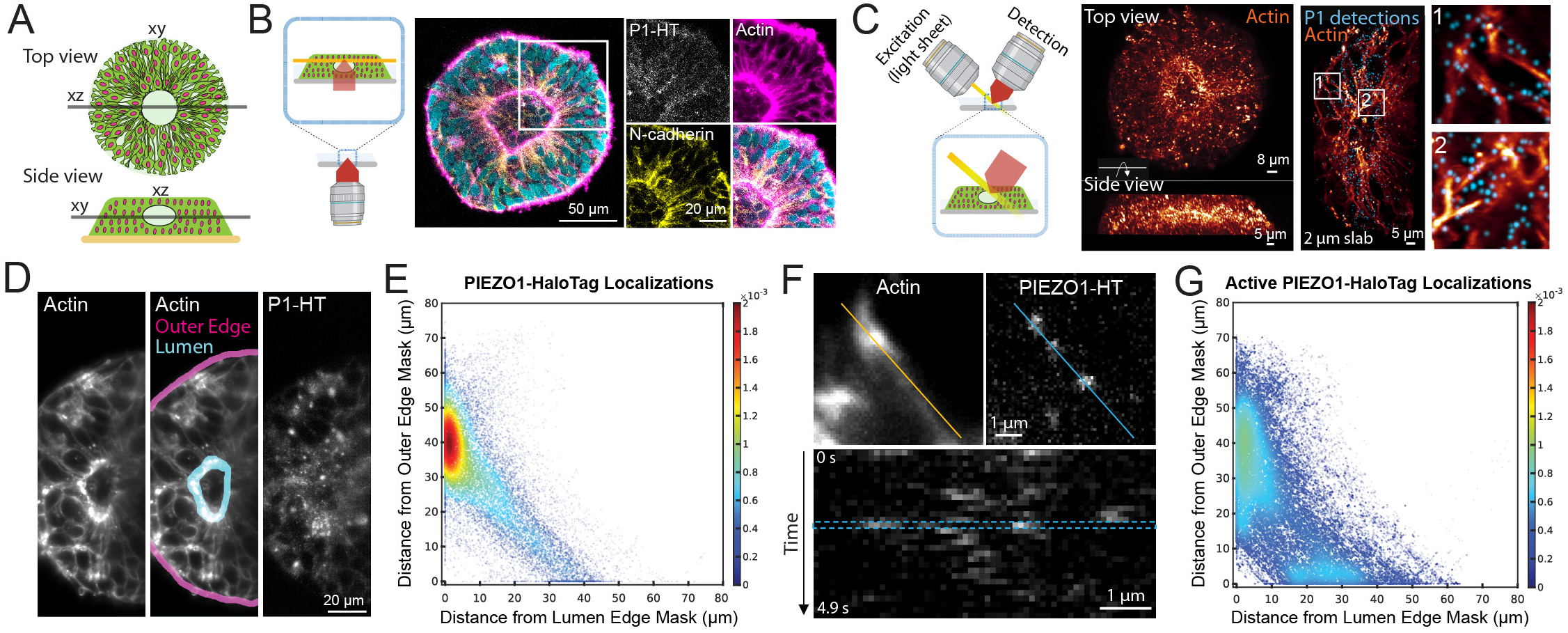
Visualizing the spatial distribution and activity of PIEZO1-HaloTag puncta in micropatterned neural rosettes (MNRs). **A.** Top and side view schematics of an MNR, illustrating cells organized radially around a central lumen. **B.** Schematic view on the left shows the confocal microscopy imaging plane parallel to the coverslip (in orange) in an MNR. Images show a representative confocal slice of a PIEZO1-HaloTag MNR labeled with JF646, fixed, and then stained with phalloidin (magenta), anti-N-cadherin antibody (yellow) and Hoechst (cyan). The section within the white square is shown zoomed in to the right for each channel. A gamma of 0.5 was applied on the zoomed-in actin image. Note the actin-rich regions at the lumen and outer edges, and the PIEZO1-HaloTag localization at the cell-cell interfaces. **C.** Schematic view on the left shows the orientation of the lattice light-sheet microscopy imaging plane in MNRs. Images on left show representative volumetric rendering of an actin-labeled MNR showing top-down and side views. The middle image panel shows a representative 2-µm slab projection of actin (orange) with JF635-labeled PIEZO1-HaloTag detections (cyan) and two zoomed insets at far right. **D.** Representative maximum intensity projection (MIP) image of 3 optical planes acquired 215 nm apart of an MNR labeled with actin (left); the same image with the lumen and outer edge masks marked for analysis (middle); and PIEZO1-HaloTag labeled with JF635 HTL (right). **E.** Density scatter plot of distances of JF635 HTL labeled puncta localizations to the lumen edge mask (x axis) and to the outer edge mask (y axis) of the MNR (n = 103 videos from 21 MNRs from 4 experiments). The color scale indicates the relative density of puncta at each position in the scatter plot, scaled to the total number of puncta represented in the plot. Note the enrichment of PIEZO1 channels near the lumen edge mask (red cluster). Density scatter plots for each individual MNR sample can be found in Supplemental Fig. 16. **F.** Top: Representative MIPs from 3-plane stacks of actin and of Ca^2+^-sensitive JF646-BAPTA labeled PIEZO1-HaloTag puncta in an MNR. Note how puncta along the blue line in the right panel localize with actin along the orange line in the left panel. The blue line in the PIEZO1-HaloTag panel indicates the region of interest used to generate the kymograph at the bottom. Note the flickering behavior of BAPTA-labeled puncta, indicating fluctuations in channel activity levels. **G.** Density scatter plot of distances of JF646-BAPTA HTL labeled active PIEZO1-HaloTag puncta localizations to the lumen edge mask (X axis) and to the outer edge mask (Y axis) of the MNR (n = 39 videos from 12 MNRs from 3 experiments). The color scale indicates the relative density of puncta at each position in the scatter plot, scaled to the total number of puncta represented in the plot. Active channels localize primarily near the actin-rich lumen, with a smaller cluster near the actin-rich outer edge of the MNR. Density scatter plots for each individual JF646-BAPTA HTL sample can be found in Supplemental Fig. 19. See also Supplemental Figs. 17, 18, 20, and 21 and Supplemental Video 10.

Due to the limitations of confocal microscopy for volumetric time series imaging, we employed adaptive optical lattice light-sheet microscopy (AO-LLSM)^61^ for rapid, high-sensitivity 3D imaging of live MNRs with minimal phototoxicity. We labeled live PIEZO1-HaloTag MNRs with JF635 HTL, as well as markers for actin and nuclei to delineate MNR morphology. We computationally separated the punctate PIEZO1 signals from the diffuse, non-specific autofluorescence also observed in PIEZO1-HaloTag Knockout MNRs and unlabeled PIEZO1-HaloTag MNRs (see Methods and Supplemental Results). AO-LLSM imaging revealed the central lumen, the radiating actin cytoskeleton and the PIEZO1-HaloTag JF635 HTL signal throughout the MNR volume (Fig. 5C, Supplemental Video 10).

For quantitative analysis of PIEZO1-HaloTag puncta distribution in live MNRs, we captured images of 3 optical planes over time to generate a time series consisting of 30 time points for a total average duration of 5.5 seconds. We then generated a maximum intensity projection image (MIP) over the 3 planes to obtain an image stack over time. We manually constructed masks representing the lumen and outer edges of the MNR in the MIP images (Fig. 5D), computationally identified PIEZO1-HaloTag puncta, and measured the distances of each punctum from the lumen and the outer edge mask (see Methods). Density scatter plots of the distances showed a clustering of PIEZO1-HaloTag puncta near the lumen edge (Fig. 5E, Supplemental Fig. 16). For the entire dataset of 103 videos from 21 MNRs, PIEZO1-HaloTag puncta distances were distributed with a mode close to 0 µm from the lumen edge mask and ∼40 µm from the outer edge mask (Supplemental Fig. 16A), further supporting lumen enrichment. Cumulative distribution plots of distance data showed that 38 ± 1% of detected PIEZO1 puncta were within 5 µm of the lumen edge mask, while only 9 ± 1% were within 5 µm of the outer edge mask (Supplemental Figs. 17A and 18A), indicating significant concentration of PIEZO1 channels at the lumen border.

To study PIEZO1 channel activity in MNRs, we labeled PIEZO1-HaloTag MNRs with the Ca^2+^-sensitive JF646-BAPTA HTL. Similar to TIRF measurements, we observed puncta exhibiting flickering behavior (Fig. 5F), albeit with lower temporal resolution of the lightsheet modality. We computationally thresholded active PIEZO1-HaloTag puncta (see Methods and Supplemental Results) and then quantified the distances of active PIEZO1-HaloTag puncta from the MNR lumen and outer edge masks, as for the JF635 HTL-labeled puncta above. We noted a diffuse cluster near the lumen edge and an additional smaller cluster near the outer edge (Fig. 5G, Supplemental Fig. 19). The localization of active PIEZO1 channels in the vicinity of the actin-rich lumen and edge regions suggests that channels in these regions are activated by cell-generated actomyosin forces, as we previously demonstrated in single cells^30^. Cumulative distribution plots of distance data from 39 videos in 12 MNRs revealed that 28 ± 3% of detected active PIEZO1 puncta localized within 5 µm of the lumen edge mask, while 20 ± 5% were within 5 µm of the outer edge mask (Supplemental Figs. 17B and 18B). Therefore, there were only 40% more active channels at the lumen border than near the outer edge, even though there were over four times as many channel puncta overall at the lumen, suggesting that tissue forces at the outer edge more efficiently activate PIEZO1 channels. These observations open future research avenues examining specific mechanisms by which channels are preferentially recruited to the lumen and outer edge regions, how cell- and tissue-level forces in MNRs activate PIEZO1, and the physiological impacts of this activity.

## Discussion

Here, we genetically engineered a human induced pluripotent stem cell (hiPSC) line by fusing a HaloTag protein to endogenous PIEZO1. By editing both alleles of the PIEZO1 gene, we ensured that all expressed PIEZO1 protein is tagged with the HaloTag. This modification, which preserves channel function, creates a chemogenetic tag for the channel that is compatible with a diverse array of specially designed HaloTag ligands (HTLs). We focus on imaging-based applications, utilizing the array of bright and photostable Janelia Fluor HTLs to visualize the localization and activity of individual PIEZO1 channels in a range of hiPSC-derived cell types and tissue organoids. Additional applications of the HaloTag technology, such as biochemical studies to identify interacting partners^62^, high-yield protein purification^46^, and Protac-mediated targeted protein degradation^63^, are also possible. Our PIEZO1-HaloTag hiPSC model thus synergizes the multifunctionality of HaloTag technologies with the versatility of hiPSCs, offering a significant leap over traditional methods in studying PIEZO1, a channel that has emerged as a critically important mechanotransducer in a wide range of physiological processes.

The PIEZO1-HaloTag hiPSC line allows for the visualization of individual mechanically-activated PIEZO1 puncta in the native cellular environment. This approach circumvents the limitations of overexpression systems, such as altered channel localization and activity due to changes in channel density. Our model also offers the flexibility to differentiate into multiple cell types and tissue organoids, providing a more comprehensive understanding of PIEZO1 dynamics compared to previous studies limited to single cell types. The superior brightness and photostability of the Janelia Fluor HTLs allow us to capture PIEZO1 localization dynamics with higher precision than previous PIEZO1-tdTomato reporter models^13,30,39^, revealing two distinct mobility behaviors and enabling imaging of the rear-enrichment of PIEZO1 in migrating cells with greater temporal resolution than previously possible^13^. Thus, our tool paves the way for a new generation of studies to examine PIEZO1 motility and localization under a variety of physiological conditions.

A further outcome of our study is the establishment of a novel single-channel PIEZO1 activity assay within the native cellular milieu. The HaloTag domain is attached to the PIEZO1 C-terminus, located directly below the pore domain; thus attachment of a Ca^2+^-sensitive HTL enables PIEZO1 activity measurements by detecting Ca^2+^ influx through the pore. This assay marks a significant advance over patch clamp electrophysiology, the standard method for measuring PIEZO1 activity. Whole-cell patch clamp dialyzes the cell, disrupting cellular structures and physiology, while cell-attached patch clamp imparts a large resting tension on the patch that may alter channel behavior. Neither modality provides spatial information on channel activation, whereas imaging approaches provide multiplexed activity measurements from dozens to hundreds of channel puncta simultaneously while maintaining cell integrity^64^. Additionally, our approach provides highly specific labeling of endogenous PIEZO1, single-channel measurement capabilities in intact cells, simultaneous readouts of PIEZO1 mobility and activity, spatial information on PIEZO1 activation, and high spatial and temporal resolution. These significant technical advancements over previous imaging-based approaches to measure PIEZO1 activity^26–28,30,32^ demonstrate our tool to be exceptionally suited for examining both PIEZO1 localization and activity dynamics under native cellular conditions in isolated cells as well as tissue organoids, and open new avenues to study regulation of PIEZO1.

Our findings advance existing methodologies, such as GenEPi65, a genetically-encoded fluorescent reporter of PIEZO1 activity, which relies on overexpression and is limited by poor kinetics and weaker signals. Activity measurements from PIEZO1-HaloTag revealed multiple levels of JF646-BAPTA HTL fluorescence intensity from single, stationary puncta. Although saturated labeling by the probe involves three HaloTag ligands per PIEZO1 trimer, the high [Ca^2+^] in the immediate proximity of a channel pore, estimated at > 15 µM ^54^, implies that the three HTLs would respond almost simultaneously to Ca^2+^ flux, given their proximity to the channel pore and high affinity for Ca^2+^ (Kd 0.14 µM)^44^. However, the observed “bright state” durations may overestimate channel open times as a consequence of the time courses of Ca^2+^ unbinding from the HTL and of Ca^2+^ diffusion away from the pore. We speculate that multiple levels of JF646 BAPTA HTL fluorescence may result if each diffraction-limited punctum represents a cluster of two or more PIEZO1 channels. Differences in amplitudes of signal from different PIEZO1-HaloTag puncta could thus arise from differences in number of PIEZO1 channels per punctum, or due to varying distances of the membrane within the exponentially decaying evanescent field, or sub-saturation labeling of some channels. Importantly, recent super-resolution studies using MINFLUX^56,66^ imaging of PIEZO1 demonstrate clustering of PIEZO1^67^.

Our studies also reveal the capability of the Ca^2+^-sensitive JF646-BAPTA HTL to spatially map PIEZO1 mobility in addition to activity, and we find that both stationary and mobile puncta can be active. However, the low resting fluorescence of the HTL in the absence of bound Ca^2+^ poses challenges in detecting and localizing PIEZO1-HaloTag channels when they are closed. This limitation may be addressed by the development of Ca^2+^-sensitive HTL variants with higher resting-state brightness, and by further engineering the hiPSC line to enable orthogonal dual labeling of PIEZO1 with spectrally distinct Ca^2+^-sensitive and insensitive HTLs.

The PIEZO1-HaloTag hiPSC also allows for the comparison of relative differences in PIEZO1 activity between distinct cell types. NSCs display less active PIEZO1 puncta than ECs which we speculate may be due to differences in cell type-specific contractility that activates PIEZO1^30^. We also provide proof-of-principle that the PIEZO1-HaloTag platform allows measurement of PIEZO1 activity in response to externally-applied mechanical stimuli such as osmotic stimulus. These activity measurements can be extended to other mechanical stimulus modalities in the future.

Beyond the single cell level, our PIEZO1-HaloTag hiPSC model offers a new approach to studying PIEZO1 at tissue scales using *in vitro* human organoid systems. Building upon prior research in a mouse model that reveals PIEZO1’s function in neural development^12^, we examine the channel’s localization and activity in micropatterned neural rosettes (MNRs), an *in vitro* model of early human neural tube development. Utilizing AO-LLSM for high-resolution live imaging, we observe a notable enrichment of PIEZO1 channels at the lumen edge of MNRs (Fig. 5E). Interestingly, channel activity is observed at both the lumen and outer edges (Fig. 5G), areas marked by high mechanical tension as evidenced by strong actin staining. Previous measurements in micropatterned neuroectoderm models showed that the outer edge experiences high cell-generated forces^68^, which are known to activate PIEZO1^30^. Our observations highlight the need to consider the site-specific biology of tissues and their unique geometrical and mechanical properties when studying physiological roles of PIEZO1.

In conclusion, our study establishes a novel and powerful approach to study PIEZO1 using hiPSCs. The adaptability of the hiPSC model, capable of differentiating into a variety of human tissue organoids, creates opportunities to biophysically and physiologically study PIEZO1 function at the tissue scale. Additionally, we can introduce pathogenic PIEZO1 Gainor Loss-of-Function mutations^15,18,19,69^ in the PIEZO1-HaloTag hiPSCs for human disease modeling of PIEZO1 channelopathies. Finally, the precision in measuring activity and localization measurements of wild-type and mutant PIEZO1 channels opens up new avenues in drug screening for novel therapeutic interventions targeting PIEZO1. Our approach provides a platform for enhancing our understanding of PIEZO1’s role in cellular mechanics and signaling in normal physiology as well as disease conditions, and holds significant potential for developing therapeutic strategies in PIEZO1-mediated human diseases. Future improvements, including continued developments in the rapidly evolving areas of HaloTag technologies as well as advanced imaging techniques, will further refine our ability to study PIEZO1’s intricate role in human health and disease. More broadly, our approach could be applied to other Ca^2+^-permeable membrane proteins.

## Methods

### hiPSC Culturing and Maintenance

All human stem cell experiments were carried out in accordance with approved Human Stem Cell Research Oversight (hSCRO) Committee of the University of California, Irvine guidelines. Stem cell lines used had no patient identifiers. hiPSCs were maintained in mTeSR^TM^ Plus Basal Medium (Cat. No.100-11300, STEMCELL Technologies) + Primocin (Cat. No.NC9392943, Invivogen) in an incubator at 37°C with 5% CO_2._. Cells were cultured on plates incubated with 10 µg/mL Vitronectin XF (Cat. No.07180, STEMCELL Technologies) for at least 1 hour at room temperature prior to passaging. Media was changed every other day, and cells were passaged mechanically or as single cells (10,000 cells/cm^2^) every 4-7 days. For generating single cell suspensions of hiPSCs, cells were dissociated with Accutase (Cat. No.# 07920_C, STEMCELL Technologies) for 3-5 minutes in a 37°C incubator. Accutase was diluted 1:1 with mTeSR Plus, 100ug/ml Primocin, and 10 μM Y-27632 (Cat. No.SM-0013-0010, Biological Industries, USA) and cells were centrifuged at 210 g for 5 minutes. For TIRF imaging, cells were seeded at 13,000 cell/cm^2^ density on Vitronectin XF coated MatTek dishes (Cat. No.P35G-1.5-14-C, MatTek Corporation Cells were regularly checked for mycoplasma contamination and karyotypic abnormalities.

### CRISPR Engineering

All PIEZO1 edits were outsourced to Sythego, Menlo Park, CA. and made in WTC-hiPSCs with normal karyotype. The ICE (Inference of CRISPR Edits) software analysis package developed by Synthego was used for analysis of CRISPR editing data. All clones had normal karyotype and passed pluripotency tests (OCT4 and SSEA immunostaining) after CRISPR editing. Karyotyping of clones was performed with Karystat analysis (ThermoFisher).

### PIEZO1-HaloTag hiPSCs

Briefly, the strategy employed for the CRISPR engineering was as follows. hiPSCs were transfected with Cas9, sgRNA (5’-GUGGACUCGUGAGAAGGAGU-3’) and knock-in template sequence which was designed with a 5’ homology arm (HA) of 496 bp upstream of the stop codon of PIEZO1, an 18 bp linker, the HaloTag coding sequence followed by the 3’ HA containing the TAG stop codon and 499 bp of the 3’ untranslated region of PIEZO1. The 18 bp linker, ggatccggtgcaggcgcc, encodes the amino acid sequence GSGAGA. HaloTag 7 is 297 amino acids encoded by 891 bp sequence. Transfected cells were screened for the presence of the insertion by PCR of the genomic DNA with FWD primer (5’-3’): GCCAAGCTCATCTTCCTCTAC and REV primer (5’-3’): GAACATGAAGGACTTGGTGAGTA which should yield a 1,669 bp product, while unedited gDNA would yield a 760 bp product.

### PIEZO1-HaloTag Knockout hiPSCs

PIEZO1-HaloTag hiPSC cells were edited to obtain homozygous indels in multiple exons within the PIEZO1 coding region in exons 6 and 43 and clonal lines were isolated. Cas9 and guide RNAs were transfected sequentially into PIEZO1-HaloTag hiPSC cells. Guide RNAs used were sgRNA (5’-UGGAUGCCAGCCCGAC-GGCA-3’) targeting exon 6 and sgRNA (5’-UCCGCCUACCAGAUCCGCUG-3’) targeting exon 43 of Piezo1. Forward primer 5’-AGGTAGACACTGGAGAGGGC-3’ and reverse primer 5’-CAGAGGAGCAGCTGTG-GATG-3’ were used in PCR amplification of genomic DNA from transfected cells. Sequencing of the PCR fragment from Clone E1 revealed a homozygous -1 indel in exon 6. Indels in exon 43 were identified using the forward primer 5’-ACCTTCTCTGTCTCTCGGCT-3’ and the reverse primer 5-ACCTTCTCTGTCTCTCGGCT-3’ for PCR amplification. Sequencing of the fragment revealed a homozygous a -5 indel in exon 43.

## hiPSC Differentiation

### Neural Stem Cell (NSC) Differentiation

Neural stem cells were differentiated from hiPSCs using STEMdiff™ Neural Induction Medium (Cat. No. 05839, STEMCell Technologies) monolayer culture protocol as per the manufacturer’s instructions. Briefly, hiPSCs were passaged using Accutase as described above and resuspended in STEMdiff™ Neural Induction Medium containing SMADi and 10 µM Y-27632. Cells were plated at 2 x 10^5^ cells/cm^2^ onto tissue culture plates coated with 10 µg/ml of CellAdhere™ Laminin-521 (Cat. No.77003, STEMCELL Technologies). Media changes (without Y-27632) were performed daily and cells were passaged at day 7, 14 and 21. NSCs were used from Day 21 and cultured in STEMDIFF™ Neural Progenitor Medium (Cat. No. 05833, STEMCell Technologies).

### Endothelial Cell (EC) Differentiation

Endothelial cells were differentiated following the S1-S2 method^70^. hiPSCs were dissociated into single cells using Accutase and plated on 10 µg/mL Vitronectin XF (Cat. No. 07180, STEMCELL Technologies)-coated plates with 10 µM of Y27632 (Cat. No.SM-0013-0010, Biological Industries, USA) in mTeSR^TM^ Plus Basal Medium (Cat. No.100-11300) + Primocin (Cat. No.NC9392943, Invivogen). 3 x 10^5^ cells/cm^2^ cells were seeded per well of a 6-well tissue culture plate. 24 hours after seeding, the media was changed to S1 media. S1 media was prepared first by making a Basal medium consisting of Advanced Dulbecco’s modified Eagle’s medium (DMEM)/F12 (Cat. No. 12334010, Thermo Fisher Scientific), 1x Glutamax supplement (Cat. No. 35050061, Thermo Fisher Scientific), 60 µg/ mL L-Ascorbic Acid (Cat. No. A8960, Sigma-Aldrich), and Primocin. Basal media was supplemented with 6 μM CHIR99021 (Cat. No. SML1046-5MG, Sigma-Aldrich) to create S1 media. Media was changed every 24h for two days. After two days in S1 media, media was changed to S2. S2 media was prepared by supplementing Basal media with 10µM SB431542 (Cat. No. S1067, Selleck Chem), 50 ng/mL bFGF-2 (Cat. No. 100-18B-100ug, PeproTech), 50 ng/mL VEGF-A (Cat. No. 100-20, PeproTech), and 10 ng/mL EGF (Cat. No. AF-100-15-100ug, PeproTech). Media was changed every 24h throughout the protocol. After a total of 48 h in S2 media, cells were purified using an immunomagnetic CD31 MACs sorting kit (Cat. No. 130-091-935, Miltenyi) and 15,000 cells were plated on 10 µg/mL Fibronectin (Cat. No. 356008, Corning)-coated MatTek dishes (Cat. No. P35G-1.5-14-C, MatTek Corporation) and cultured for 48 h in EGM-2 media (Cat. No.CC-3162, Lonza) before imaging.

### Keratinocyte Differentiation

Mechanically passaged hiPSCS were plated on 10 µg/mL Vitronectin XF in 6 well TC plates in 1 μM alltrans-RA (Cat. No. R2500-25MG, Sigma-Aldrich) to NutriStem® hPSC XF Medium (Growth Factor-Free)(Cat. No.06-5100-01-1A, Biological Industries Israel Beit-Haemek Ltd.). Daily media changes were performed with media containing 1 µM RA for 7 days. On day 7, differentiated cells were passaged with 2 mg/mL of Dispase (Cat. No. CnT-DNP-10, Cellntec) incubated for 30 min at 37C. Cells were resuspended in 1 mL of media CnT-Pr media (Cat. No. CNT-PR, CellnTec) and 1/10^th^ of total cells were plated on 10 µg/mL Fibronectin (Cat. No. 356008, Corning) coated MatTek dishes (Cat. No.P35G-1.5-20-C, MatTek Corporation). Cells were cultured in CnT-Prime Epithelial Proliferation Medium (Cat. No. CNT-PR, CellnTec) for 2-4 weeks prior to imaging. Media was exchanged to CnT-Prime Epithelial 2D Differentiation Medium (Cat. No. CnT-PR-D, CellnTec) for 2-4 days prior to imaging.

### Preparation and Differentiation of Micropatterned Neural Rosettes (MNRs)

MNRS were generated from hiPSCs using a method developed by Haremaki et al. 2019^60^ using Arena A CYTOOchips (Cat. No.10-020-00-18, CYTOO INC.). A CYTOO Arena A chip was placed on parafilm in a 10-cm petri dish (Cat. No. 08757100D, Fisher Scientific) the top side was coated with 10 µg/ml of CellAdhere™ Laminin-521 (Cat. No.77003, STEMCELL Technologies) for 3 hours at 37°C. Laminin was then removed via multiple washes in PBS + calcium + magnesium (PBS +/+) (Cat. No. PBL02-500ML, Caisson Laboratories Inc.). The CYTOOchip was then moved to a single well of a 6 well tissue culture plate with PBS +/+ prior to plating. PIEZO1-HaloTag and PIEZO1-HaloTag Knockout hiPSCs were washed with PBS +/+ and incubated with accutase at 37°C for 3-5 minutes to generate a single cell suspension. Accutase was diluted 1:1 with mTeSR Plus, containing 10 μM Y-27632 and spun at 210 g for 5 minutes. Cells were resuspended, counted and seeded at a concentration of 5×10^5^ hiPSCs in 2 mL of media on top of the micropattern in a single well of a 6 well plate. The media was changed after 3 hours into differentiation media comprising of a 1:1 mixture of DMEM/F12 and Neurobasal containing 1:100 Glutamax, 1:200 Non Essential Amino Acids, 1:200 N2 supplement, 1:100 B27 without vitamin A (all Invitrogen), 3.5 µL l^-1^ 2-mercaptoethanol (Sigma), 1:4000 insulin (Sigma), 10 μM SB431542 (Cat. No. S1067, Selleck Chem) and 0.2 μM LDN193189 (Cat. No. SM-0005-0010, Biological Industries, USA). Media was changed daily and MNRs were kept in culture until day 5 in a 37°C incubator, at which time central lumens were present. MNRs were incubated with HaloTag Ligand as detailed below and imaged on day 5. To label actin structures, MNRs were incubated with SPY555-actin (Cat. No. CY-SC202, Cytoskeleton, Cytoskeleton) at 1:1000 in culture media, 1:1000 SPY505-DNA (Cat. No. CY-SC101, Cytoskeleton) was added to label DNA, both for 1 hour prior to imaging. MNRs were washed 3 times in culture media prior to imaging.

Small adjustments were made to the above protocol to prepare MNRs for confocal microscopy. The CYTOO chip was placed in the bottom of a 35 mm glass-bottom dish without coverglass (Cat. No. D35-14, Cellvis). Thus, the micropatterned surface was positioned at the bottom in the hole. Subsequently, biosafe glue (KWIK-CAST silicon sealant, World Precision Instruments) was added to the bottom of the 35 mm dish around the edges of the CYTOOchip to seal it in place. The chip was rehydrated with 2mL PBS +/+ for 5 minutes. The same protocol as above was followed, coating the chip with laminin and seeding the cells directly into the 35 mm dish. MNRs were labeled with HaloTag Ligand on Day 5 as detailed below and imaged live for AO-LLSM or after fixation for confocal imaging.

### Mouse Liver Sinusoidal Endothelial (mLSEC) Isolation and Culture

All animal experiments were carried out in accordance with approved Institutional Animal Care and Use Committee protocols and University of California, Irvine guidelines. Livers from the PIEZO1-tdTomato mice^3^ were dissected and placed on a petri dish and minced with scalpel blades. Once the liver was minced, it was resuspended in a solution to further dissociate the tissue. This solution contained 1 mL 2.5 U ml-1 dispase, 9 mL 0.1% collagenase II, 1 µM MgCl_2_, and 1 µM CaCl_2_ in Hanks Buffer. The tissue was incubated with the dissociation mixture for 50 mins in a tube rotator with continuous agitation at 37°C. After the completion of the dissociation process, the dissociated tissue was filtered using 70 and 40 µm cell strainers. The dissociated cells were then washed twice in PEB buffer containing phosphate-buffered saline solution (PBS), 0.5% BSA, pH 7.2, EDTA 2 mM, and EDTA 2 mM. Once the pellets were washed, cells were placed in 1 mL PEB buffer and 30 µL CD146 microbeads (Cat. No. 130-092-007, Miltenyi Biotech) at 4°C for 15 min under continuous agitation. An LS column (Cat. No. 130-042-401, Miltenyi Biotech) was primed with PEB buffer during this incubation. Once the mLSECs were selected for using the microbeads, the solution was then placed in the primed LS column to separate the mLSECs (Miltenyi Biotech). After the cells passed through the column, the column was washed 3 times with 5 mL PEB buffer. Any CD146 positive cells were eluted from the column using 5 mL warmed EGM-2 growth medium supplemented with EGM-2 bullet kit (Lonza). The cells were pelleted with a 300 g spin for 5 min and counted in 1 mL EGM-2 media. Glass-bottom #1.5 dishes were prepared by coating with 10 µg/mL Fibronectin (Cat. No. 356008, Corning). After performing the cell count, approximately 30,000 cells were plated on each fibronectin coated glass-bottom dish. After 2 hours, a media change was performed and then media changed every 48 h until imaging 72 h later.

### Cell Lysis and Immunoblotting

iPSC cells were lysed in RIPA buffer (Thermo Scientific™ 89901) containing 1 µM Dithiothreitol, Thermo Scientific Halt Protease Inhibitor Cocktail (Cat. No. 78430) and Halt Phosphatase Inhibitor Cocktail (Part No. 78420)) for 15 min on ice. Lysates were sonicated (SONICS Vibra-cell, probe model CV184) at 100% power twice (10 seconds pulse on, 20 seconds off) then centrifuged 10 min x 16,100 x g at 4C. The supernatant was transferred to a new tube and the protein concentration was determined with Pierce BCA assays (Thermo Scientific™ 23225). Proteins were separated by SDS-PAGE by loading 15 micrograms of lysate on a NuPAGE 3-8% Tris Acetate gel, (ThermoFisher Cat. No. EA0375) in Tris Acetate SDS Running Buffer (Cat. No. LA0041), then transferred to 0.45 micron PDVF membrane (Thermo,FisherCat. No. 88518) with the Biorad Mini-PROTEAN Tetra System in 25 mM Tris, 192 mM Glycine, 20% Methanol overnight at 4C for 18 hrs at 30V. Western blotting was performed with 1:1000 mouse anti-HaloTag (Promega, Cat. No. G9211) or 1:1000 mouse anti-PIEZO1 (Novus Biologicals, Cat. No. NBP2-75617), followed by 1:1000 Goat anti-Mouse IgG-Secondary antibody, HRP (Invitrogen, Cat No. 32430) or 1:100,000 HRP-mouse anti-actin (ThermoFisher Cat. No. 5-15739-HRP) in TBS, 0.1% Tween20 in 5% NonFat Dried Milk (Carnation). Antigen detection was performed with Thermo SuperSignal West Femto Max sensitivity (ThermoFisher, Cat. No. 34095) with a Bio-Rad Gel Chemi-Doc Molecular Imager.

### HaloTag Ligand Treatment Protocol

After differentiation of the PIEZO1-HaloTag hiPSC lines into either NSCs, Endothelial cells, Keratinocytes, or MNRs, each respective cell type was incubated at 37 °C with 500 pM of Janelia Fluor® 646 HaloTag Ligand (NSCs, ECs, or keratinocytes) (Cat. No. GA1120, Promega) or Janelia Fluor® 635 HaloTag Ligand (MNRs) (Cat. No. CS315103, Promega) or Janelia Fluor® 646-BAPTA-3’AM-HaloTag ligand (requested from Lavis lab, Janelia Research Campus, HHMI) for incubation time outlined in Supplementary Methods Table 1, “Table of HaloTag Ligands”. MNRs were treated with JF635 HTL due to its improved fluorogenic properties and compatibility with existing filter sets in the AO-LLSM system^71^. Incubations with HTLs were performed in each respective cell type’s basal media outlined in their respective culturing sections. Cells were gently washed 5 times with DMEM/ F12 1:1 Cat. No. 25116001, Invitrogen) at room temperature prior to imaging using TIRF microscopy. MNRs were washed 3 times with culture media prior to fixation or live imaging.

### Immunofluorescence

Samples plated in MatTek dishes (Cat. No. P35G-1.5-14-C or P35G-1.5-20-C, MatTek Corporation) were first fixed in a 4% v/v paraformaldehyde solution, including 5 mM MgCl_2_, 10 mM EGTA, and 40 mg/mL sucrose in PBS (pH = 7.3) for 10 minutes at room temperature. Next, the fixed cells or micropatterned neural rosettes were permeabilized with 0.3% Triton X-100 in PBS for 5 minutes. 5% BSA in PBS (Jackson ImmunoResearch, Cat. 001-000-162) for 1 h at room temperature was used to block nonspecific binding of excess antibodies. After blocking was complete, the sample was incubated overnight at 4°C with primary antibody diluted in 1% BSA (Primary antibodies used and their concentrations are included in supplementary methods). Next, samples were washed several times to remove the primary antibody and then were incubated with secondary antibody for 1 hour at room temperature (Secondary antibodies used and their concentrations are included in supplementary methods). Samples were washed several times and subsequently labeled with Hoechst at 1 µg/mL for 5 minutes at room temperature. Samples were stored in 1× PBS prior to imaging.

### Cell-attached patch clamp

Cell-attached patch clamp experiments were made using an Axopatch 200B amplifier (Molecular Devices) at room temperature. Pipettes were made from thin walled borosilicate glass capillaries (Warner Instruments) and contained a solution consisting (in mM) of 130 NaCl,10 HEPES, 10 tetraethylammonium chloride, 8 Glucose, 5 KCl, 1 CaCl_2_, 1 MgCl_2_, (pH 7.3) with NaOH. Bath solution composition included (in mM) 140 KCl, 10 Glucose, 10 HEPES, 1 MgCl_2_, (pH 7.3 with KOH) was used to zero the membrane potential. The pipettes had a resistance of 0.6-1.1 MΩ when submerged in these solutions. Negative-suction application provided mechanical stimulation during recordings using a high-speed pressure clamp (HSPC-1; ALA Scientific) controlled using Clampex software. Suction pulses were applied using the patch pipette and membrane potential was held at -80 mV. Off-line leak subtraction was performed prior to quantification of maximum current. Maximum current was manually recorded and quantified for statistical differences using students t-test and mean effect size using Cohen’s *d*.

### Whole cell patch clamp

Whole cell patch clamp experiments were conducted according to the protocol described previously with slight modifications ^55^. Patch pipettes were pulled from thin-wall borosilicate capillaries with internal filament (GC 150TF-7.5; Harvard Apparatus). Recording pipettes were pulled to a resistance of 2 to 3 MΩ and fire polished using a microforge (MF2, Narishige. Standard HBSS (Cat. No. 14025092, GIBCO) was used as the bath saline. Patch pipettes were filled with 140 mM KCl, 10 mM HEPES, 10 mM TEA, and 2 mM EGTA (pH 7.4 with NaOH) saline. Cells were held at -60 mV after whole cell entry and during recording. Due to the flatter morphology of both neuronal and endothelial cells, which restricted the maximum poking distance, we implemented modifications to the poking protocol. Poking probes were made by fire-polishing the tip of glass pipettes to a relatively larger size of ∼4 to 6 to achieve sufficient mechanical stimulation despite the limited poking distance. Indentation stimuli were delivered by displacing poking probes with a piezoelectric actuator (P-841, Physik Instrumente) controlled by Clampex via an amplifier (E-625, Physik Instrumente). A shallow poking angle of ∼40° used to increase the horizontal poking distance and minimize the vertical poking distance. Before each experiment, the probe was slowly moved until visual deformation of the cell. The probe was then retracted ∼1 to 2 µm to set the initial probe position as close as possible to the cell surface without physical contact. Initial Poking distance was 3 µm. If poking induced current was not observed the poking distance was increased to 4 µm and then 6 µm until measurable current was observed.

### Total Internal Reflection Fluorescence (TIRF) Microscopy Imaging

TIRF microscopy was used to image endogenous PIEZO1-tdTomato and PIEZO1-HaloTag channels at 37°C. PIEZO1-HaloTag cells were incubated in accordance with the HaloTag Ligand Treatment Protocol above, and washed thrice with phenol red-free DMEM/F12 1:1 (Cat. No. 25116001, Invitrogen) and incubated in imaging solution, composed of 148 mM NaCl, 3 mM CaCl_2_, 1 mM KCl, 2 mM MgCl_2_, 8 mM Glucose, 10 mM HEPES, pH 7.30, and 316 mOsm/L osmolarity. The hypotonic imaging solution was made by diluting the control solution with 20% MQ water. PIEZO1-HaloTag and PIEZO1-tdTomato samples in Fig. 1, Supplemental Fig. 3, Supplemental Video 1, Supplemental Video 2, Supplemental Video 3, Fig. 2A, and Fig. 2B were all imaged using an Olympus IX83 microscope fitted with a 4-line cellTIRF illuminator, an environmental control enclosure and stage top incubator (Tokai Hit), programmable motorized stage (ASI), a PLAPO 60x oil immersion objective NA 1.45 (1 pixel is approximately 0.109 µm, no pixel binning), and a Hamamatsu Flash 4.0 v2+ scientific CMOS camera. The laser power for the 560 nm laser in these experiments was 0.43 mW at the back pupil of the objective. All other samples were imaged using an Olympus IX83 microscope fitted with a 4-line cellTIRF illuminator, an environmental control enclosure and stage top incubator (Tokai Hit), programmable motorized stage (ASI), a PLAPO 60x oil immersion objective NA 1.50 (1 pixel is approximately 0.108 µm, no pixel binning), and a Hamamatsu ORCA-Fusion BT Digital CMOS camera. To image JF646-BAPTA-labeled cells in this system with the 640 nm laser, the laser power at the back pupil of the objective was 3.5 mW for 100 fps videos; 12 mW for 200 fps videos; and 20.5 mW for 500 fps videos. All images were acquired using the open-source software Micro-Manager^72^. Cells were illuminated with either a 560 nm or a 640 nm laser, as appropriate for the fluorophore used, and images were acquired with a Hamamatsu ORCA-Fusion BT Digital CMOS camera. For TIRF experiments, 1 camera unit (c.u.) is equivalent to 0.229 photoelectrons.

### Analysis of PIEZO1-HaloTag Puncta Diffusion based on TIRF imaging

To assess the diffusion properties of PIEZO1, the single-molecule localization software package Thunder-STORM^48,49^ implemented in FIJI^48,49^ was used to detect and localize single PIEZO1-HaloTag puncta observed in TIRF recordings. ThunderSTORM was set to use multi-emitter fitting. PIEZO1-HaloTag trajectories were then generated by connecting puncta localization centroids over time using the image analysis software FLIKA^50^. The nearest punctum within three pixels (centroid-centroid distance) between adjacent frames was linked and assigned to a trajectory. If no puncta were detected within adjacent frames the next frame was also searched, and if no puncta could be detected within the search radius the trajectory was terminated. Tracks with a minimum of 4-links were analyzed to calculate signal-to-background, diffusion coefficients, trajectory path length, and mean squared displacement (MSD) values. Single lag displacements (SLD) were calculated using the distance moved by the punctum between two consecutive frames. In cases where the punctum was briefly undetected (a “gap frame”), punctum positions in the gap frames were interpolated.

### Evaluation x-y Drift in Immobilized PIEZO1-HaloTag

To assess the relative contribution of x-y drift in our puncta diffusion data, we labeled PIEZO1-HaloTag endothelial cells with JF646 HTL. Then we fixed the cells to immobilize the PIEZO1-HaloTag JF646 puncta using a 4% v/v paraformaldehyde solution, including 5 mM MgCl_2_, 10 mM EGTA, and 40 mg/mL sucrose in PBS (pH = 7.3) for 10 minutes at room temperature. Samples were washed 3 times with 1× PBS and imaged with TIRF microscopy. Images were acquired previously as described in the section “Total Internal Reflection Fluorescence (TIRF) Imaging.” Immobilized puncta data was then processed using the single-molecule localization software package ThunderSTORM^48,49^ implemented in FIJI^48,49^. Thunderstorm settings were as described in “Analysis of PIEZO1-HaloTag Puncta Diffusion based on TIRF imaging.” After processing, we ran the drift correction modulus to determine how much drift was present in the immobilized data (< 0.1 pixels, 10 nm, over 5 s). We found a negligible contribution of x-y drift, less than 10 nm over 5 s.

### Calculation of localization error using fixed PIEZO1-tdTomato and PIEZO1-HaloTag samples

To compare the relative localization error between PIEZO1-tdTomato and PIEZO1-HaloTag samples, we first harvested mouse PIEZO1-tdTomato mLSECs as described in the methods section “Mouse Liver Sinusoidal Endothelial (mLSEC) Isolation and Culture.” PIEZO1-tdTomato mLSECs and PIEZO1-HaloTag endothelial cells were plated on Matteks. Cells were then fixed using a 4% v/v paraformaldehyde solution, including 5 mM MgCl_2_, 10 mM EGTA, and 40 mg/mL sucrose in PBS (pH = 7.3) for 10 minutes at room temperature. Samples were washed 3 times with 1× PBS and imaged with TIRF microscopy. Images were acquired previously as described in the section “Total Internal Reflection Fluorescence (TIRF) Imaging.” Immobilized trajectories were extracted using methods section “Analysis of PIEZO1-HaloTag Puncta Diffusion based on TIRF imaging.” Localization error was determined by measuring the distance of each puncta’s localization from the mean position of the trajectory across records from fixed samples.

### Comparison of PIEZO1-tdTomato and PIEZO1-HaloTag signals

For each PIEZO1-tdTomato and PIEZO1-HaloTag JF549 HTL video, the sum of all pixel intensities in a region of interest (integrated intensity) was measured over the duration of the video in FIJI. This was done for 3 ROIs in each video. ROIs were 71×71 pixels (7.7×7.7 µm) These values were then background-subtracted using the integrated intensity from a ROI outside the cell border of the same video. ThunderSTORM was used to identify puncta in the first frame of each of the three ROIs and the background-subtracted integrated intensity was scaled for the number of puncta. These scaled integrated intensity values from each of the 3 ROIs within a video were then averaged to create a video-level average, which were then again averaged over 20 videos for PIEZO1-tdTomato and 19 videos for PIEZO1-HaloTag to yield an overall mean for each fluorophore.

To calculate the relative signal-to-background ratio for JF549-labeled PIEZO1-HaloTag and for PIEZO1-td-Tomato puncta (for Fig. 2A), puncta were detected in the first frame for each video. The signal intensity was measured from a 3×3 pixel ROI after subtracting camera black level. Background fluorescence was similarly determined from a 3×3 pixel ROI inside the cell devoid of puncta.

### Analysis of PIEZO1-HaloTag activity based on TIRF imaging

FLIKA software was used to build trajectories from PIEZO1-HaloTag cells labeled with JF646-BAPTA and to record puncta fluorescence intensity within a 3×3 pixel ROI for each frame ^50^. To account for the flickering behavior of the JF646-BAPTA HTL, the allowable gap time was 90 ms. Intensities and positions were interpolated over gap frames. Puncta intensities were background subtracted using intensities from a 3×3 pixel ROI inside the cell devoid of puncta. After trajectory building, videos were visually inspected to identify representative stationary and motile puncta.

### Puncta density analysis

To quantify the density of PIEZO1 puncta labeled with JF646-BAPTA or JF646 HTL, we first binned the videos acquired at either 100 fps (Fig. 3, Supplemental Figure 12) or 200 fps (Fig. 4) to 10 fps, then cropped a region of interest from the first frame of the video. Using the same settings as described in Methods section “Analysis of PIEZO1-HaloTag Puncta Diffusion based on TIRF imaging”, we detected and localized PIEZO1 puncta. The number of puncta in the cropped frame was recorded and scaled to the area of the cropped region of interest.

### Calcium Calibration Experiment in Transfected WTC-11 hiPSCs

One million WTC-11 iPSC cells were transfected with 3 µg of the plasmid pCDNA5/FRT/TO_HaloTag7_ T2A_EGFP (Addgene No.169325) using AMAXA Stem Cell Kit No.1 (Cat. No. VPH-5012) and Nucleofector II with the B-016 program. pCDNA5/FRT/TO_HaloTag7_T2A_EGFP was a gift from Kai Johnsson (Addgene plasmid No. 169325) ^73^). 50,000 cells were plated on MatTek plates in mTESR+ media containing CEPT cocktail (50 nM Chroman 1, 5 µM Emricasan, 0.7 µM trans-ISRIB (Captivate Bio, Cat. No. CET01B), 1× Polyamine Supplement (Sigma-Aldrich, Cat. No. P8483-5ml) and cultured overnight at 37°C in a humidified incubator with 5% CO_2_. GFP-positive cells indicated successful transfection and expression of HaloTag7 under the control of the CMV promoter. Transfected cells were subsequently labeled with a mixture of 0.5 nM JF646-BAPTA and 0.5 nM JF549 HTLs. Then, cells were fixed in a 4% v/v paraformaldehyde solution, including 5 mM MgCl_2_, 10 mM EGTA, and 40 mg/mL sucrose in PBS (pH = 7.3) for 10 minutes at room temperature, permeabilized in 0.3% Triton X-100 in PBS for 5 minutes, and imaged with sequential changes of bath solution containing 0 nM, 75 nM, and 39 µM Ca^2+^ solutions (Calcium Calibration Kit #1, Invitrogen, Cat. No. C3008MP) at 23°C. The Calcium Calibration Kit #1 includes two 50 mL solutions: 10 mM K₂EGTA and 10 mM CaEGTA. Both solutions are prepared in deionized water and contain 100 mM KCl and 30 mM MOPS, with a pH of 7.2. The two solutions are mixed to give the desired Ca^2+^ concentration. Cells were imaged with TIRF microscopy at 10 fps. Fixed puncta fluorescence intensity within a 3×3 pixel ROI was recorded and background subtracted, as previously described in the section “Analysis of PIEZO1-HaloTag activity based on TIRF imaging”.

### Live MNR Imaging

Micropatterned neural rosettes (MNRs) of 140 µm in diameter grown on 19 mm square Arena A CYTOO chips (Cat. No.10-020-00-18) were imaged on a modified adaptive optical lattice light-sheet microscope (AO-LLSM)^61^. As described previously^74^, prior to all imaging, the microscope was calibrated to correct for optical aberrations from the system. The Cytoo chip was mounted on a custom designed sample holder and immersed in 40 ml of phenol-free MNR culture medium. The excitation (Thorlabs 0.6 NA, TL20X-MPL) and detection (Zeiss 1.0 NA, 421452-9800-000) objectives were also immersed into the imaging medium. The MNRs were imaged with a multiBessel square lattice light sheet with the NA_sq_ of 0.35 and a 0.3/0.4 NA annular mask. The 488 nm, 560 nm and 642 nm lasers were used to visualize SPY505-DNA (DNA, Spirochrome Cat. No. CY-SC101), SPY555-actin (actin, Spirochrome Cat. No. CY-SC202), and Janelia Fluor® 635-HaloTag Ligand (Non-Bapta) or Janelia Fluor® 646-BAPTA-AM-HaloTag ligand (PIEZO1), respectively, with power at the back pupil of the excitation objective of 45 µW for 488 nm, 50 µW for 560 nm and ranging between 2.1-2.9 mW for 642 nm. To balance volumetric imaging speed, signal to noise and photobleaching, images were acquired using camera exposure between 30–50 milliseconds. All data from the AO-LLSM were collected on two Hamamatsu ORCA-Fusion sCMOS cameras. Emission light from actin, DNA, and JF635 or JF646 was separated by a dichroic mirror (Chroma T600DCRB) and passed to two cameras equipped with either Semrock FF01-600/37-25 emission filter for actin or Semrock FF01-538/685-25 filter for DNA and JF635 or JF646-BAPTA.

The 3D volumetric imaging of MNRs was performed by tiling AO-LLSM across the sample. Each tile (approximately 200 x 100 x 15 µm^3^) was scanned by moving the sample stage at 400 nm step sizes. Each neighboring tile had a 5 µm overlap between each adjacent tile. Mismatch between the excitation LLS and the detection focal plane caused by the MNR were corrected prior to every acquisition and was essential to ensure optimal imaging of PIEZO1, DNA and actin structures^75^. The AO-LLSM data was processed using MATLAB versions R2022b and R2023a. The large 3D MNR volumes were stitched in skewed space, deconvolved, deskewed and rotated on Advanced Bioimaging Center’s computing cluster at UC Berkeley using the computational pipelines published on GitHub (https://github.com/abcucberkeley/LLSM5DTools). The skewed space deconvolution was performed as described previously^74^ with experimentally measured point spread functions obtained from 200 nm fluorescent beads (Invitrogen FluoSpheres Carboxylate-Modified Microspheres, 505/515 nm, F8811). The nuclei were denoised using Content-Aware Image Restoration (CARE)^76^. The training data for the denoising model was collected using lattice light-sheet microscopy as previously described^73^ by volumetrically scanning LLC-PK1 cells expressing nuclear marker (H2B) to record low SNR and corresponding high SNR 3D stacks. The AO-LLSM instrument was controlled using a custom LabVIEW based image acquisition software (National Instruments, Woburn, MA) licensed from Janelia Research Campus, HHMI.

To observe the dynamics of PIEZO1-HaloTag in MNRs, 3-plane stack videos were acquired. For this, the sample stage (using the SmarAct MLS-3252 Electromagnetic Direct-Drive) was rapidly scanned across a 1 µm sample range (comprised of three image planes spaced 400 nm along the sample stage scan axis, corresponding to 215 nm along the optical z axis) at an interval between 160-210 ms per stack for 30 time points. The dynamic time series datasets were deskewed and maximum intensity projected prior to analysis^77^. The corresponding DNA and actin volumes were recorded at the same location prior to acquiring the PIEZO1-HaloTag time series. A new dataset of 30 time points was collected every ∼10 µm throughout the MNR and was repeated on multiple samples.

### Detection of PIEZO1-HaloTag JF635 puncta and separation from spurious detections

The maximum intensity projection (MIP) images of deskewed 3-plane stacks were used for puncta localization analysis. PIEZO1-HaloTag puncta localization was analyzed in JF635-labeled PIEZO1-HaloTag MNRs. JF635-labeled PIEZO1-HaloTag Knockout MNRs and unlabeled PIEZO1-HaloTag MNRs served as controls to characterize and subsequently filter out spurious detections due to autofluorescent spots, remains of unbound JF635 probe and local background fluctuations being detected as puncta, as described below.

The ImageJ plugin ThunderSTORM^48^ was used to detect PIEZO1 puncta, generating puncta localization maps for each frame of the acquisition video. A Gaussian function was fitted to the detected puncta spots to generate a centroid for each punctum, as described above for TIRF data. To distinguish labeled PIEZO1-HaloTag puncta from autofluorescent spots, which were much larger in size and also present in PIEZO1-HaloTag KO MNRs and unlabeled PIEZO1-HaloTag MNRs, a first filtering step was applied by setting a threshold based on the standard deviation (σ) of the Gaussian fit of each punctum. Puncta with σ higher than 280 nm were removed since experimental σ measured on fluorescent beads (diameter: 0.2 μm, excitation 642 nm) was ∼189 nm.

In order to remove any spurious detections due to unbound JF635 dye passing through the plane of imaging during acquisition, or to ThunderSTORM erroneously detecting small local background fluctuations as puncta, a time-based filtering step was applied to retain only objects that persisted over multiple frames. The custom-built FLIKA algorithm^50^ was used to link detected puncta across frames with a maximum linking distance between consecutive frames of 324 nm (3 pixels) and a linking gap of one frame, as done for TIRF data analysis above. Tracks composed of at least 3 segments (i.e. successfully linked puncta in at least 4 frames) were considered representative of bona fide PIEZO1-HaloTag puncta and retained for further analysis. See the Supplementary Results section for performance of these settings in detecting and filtering PIEZO1-HaloTag puncta.

### Detection of PIEZO1-HaloTag JF646-BAPTA signal and separation from spurious detections

Puncta in MNRs labeled with JF646-BAPTA HTL were detected using ThunderSTORM as described above for JF635-labeled MNRs. The same size-based filtering step as done for the JF635-labeled samples above was applied, which removed spurious detections in autofluorescent spots with a σ over 280 nm.

The fluorescence intensity of the JF646-BAPTA HTL depends on the local Ca^2+^ concentration, which changes with the channel’s activation state. Thus, the fluorescence intensity of the JF646-BAPTA HTL is higher when PIEZO1 channels are active/open than when in the resting/closed state; as channels flicker between open and closed, the fluorescence intensity of puncta varies over time (e.g. see kymograph in main Fig. 5F). Thus the time-based filtering used for the JF635 HTL above cannot be applied for image stacks of JF646-BAPTA HTL. Hence, Control JF 646-BAPTA PIEZO1-HaloTag KO and unlabeled PIEZO1-HaloTag MNRs were used to determine the threshold intensity values for separating puncta associated with active/open channels from unbound HTL and small autofluoresence puncta. Based on the distribution of integrated intensity of puncta in the Control samples, ∼77 photons (261 c.u.) was selected as the filtering threshold. Puncta in JF646-BAPTA PIEZO1-HaloTag MNRs below this threshold were filtered out while puncta with intensities above this value were retained for analysis.

### Determining distance of PIEZO1-HaloTag puncta to MNR lumen and outer edge

In order to determine the distances between detected PIEZO1 puncta and the lumen or outer edge of the MNR, we manually drew masks identifying the lumen border and the outer edge of the rosettes on ImageJ, using the actin labeling as reference (example in main Figure 5D).

For the JF635-labeled PIEZO1-HaloTag experiments, we computed the mean position coordinates of each PIEZO1-HaloTag track. We then computed distances of this mean position for each track to the MNR lumen and outer edge masks using Euclidean distance transform, and rounded the distances up to the nearest pixel coordinates. The puncta overlapping with the masks were assigned a distance of zero for the corresponding mask. For the JF646-BAPTA-labeled PIEZO1-HaloTag signal, we used each filtered punctum localization as described above, and Euclidean distances from these to lumen and outer edges were similarly calculated.

The calculated distance distributions were visualized using density scatter plots. We used the “ksdensity” function in MATLAB with a “normal” kernel having a bandwidth of [2.503 microns, 3.661 microns] for the distances to the lumen and outer edge respectively. Once the underlying distribution of the distances is obtained, the scatter plot was drawn using the “Scatter” object in MATLAB, with the colors corresponding to their probability densities. The transparency of the scatter points in the density scatter plots is based on the density values, where darker regions correspond to higher concentration of the puncta. This normalization method ensures similar smoothing and colormap scaling of the underlying distributions across all experiments, thus allowing us to compare distribution patterns across all the conditions tested.

The individual relative frequency distributions were plotted using histograms computed from the corresponding distance distributions with a bin size of 2 microns. For the histogram plots, the actual frequency of the detections at a particular distance from the corresponding mask was normalized using the total count, along with the bin size, so that the total area is 1. These plots were generated using the “hist” and “stairs” functions in MATLAB. Also, for further analysis, the respective cumulative distribution frequency (CDF) graphs were plotted using the “ecdf” function in MATLAB on the distance distributions, which computes the empirical CDF values from the data.

### 3D Detection and filtering of PIEZO1-HaloTag puncta in volumetric samples

The PIEZO1-HaloTag puncta localization in the volumetric data was inspected by numerically fitting a model of the PSF approximated by a 3D Gaussian function, as described previously^77^. JF635 PIEZO1-HaloTag KO MNRs were used as a control to determine filtering parameters to exclude autofluorescent blobs present in the data. For filtering, thresholding was applied on the fitted amplitude of the intensity above the local background, as well as the fitted local background to extract the detected diffraction limited puncta. This approach ensured that most puncta in the Knockout sample were removed, while retaining most of the puncta in JF635 PIEZO1-HaloTag MNRs.

## Statistical Analysis

Sample sizes and number of biological replicates are indicated in corresponding figures. An online estimation stats tool (https://www.estimationstats.com)^78^ was used to calculate Cohen’s d. All p value calculations and graphs were performed using OriginPro 2020 (OriginLab Corporation). Statistical tests for p value calculations are indicated in legends.

## Acknowledgements

We thank Dr. Ian Smith, Ms. Elaine Lai and Ms. Vivian M. Leung for technical support and members of the lab for comments on the manuscript. We gratefully acknowledge Dr. Luke Lavis, Janelia Research Campus, and the Janelia Materials project team for sharing Janelia Fluor HaloTag Ligands. We thank Dr. Francesco Tombola, University of California, Irvine for the use of his patch clamp equipment; and Ms. Allia Fawaz and the Core Facilities of the Sue and Bill Gross Stem Cell Research Center (supported in part through a California Institute of Regenerative Medicine Shared Research Lab Grants (SRL): Cl1-00520). We thank Dr. Xiongtao Ruan and Matthew Mueller, UC Berkeley for helpful discussions on data analysis and visualization.

This work was supported by National Institutes of Health grants DP2AT010376 and R01NS109810 to M.M.P.. G. A. B. was supported in part by NIH T32 NS082174 and the University of California, Irvine Stanley Behrens Fellowship in Medicine, E.L.E. was supported in part by California Institute of Regenerative Medicine Training Grant EDUC4-12822, A.T.L. was supported in part by NIH F31 1F31NS127594 and the University of California, Irvine Graduate Dean’s Dissertation Fellowship, and J.R.H. by the HHMI Gilliam Diversity Fellowship. G.L., S.S. and S.U. are funded by the Philomathia Foundation. S.U. is funded by the Chan Zuckerberg Initiative Imaging Scientist program. S.U. is a Chan Zuckerberg Biohub – San Francisco Investigator.

## Supplementary Methods Information

### Reagents

#### Table of Antibodies

| Antibody | Host | Company, Cat. # | Use | Dilution |
| --- | --- | --- | --- | --- |
| anti-PIEZO1 | Mouse | Novus Biologicals, NBP2-75617 | Western Blotting | 1:1000 |
| anti-HaloTag | Mouse | Promega, G9211 | Western Blotting | 1:1000 |
| HRP-mouse anti actin | Mouse | ThermoFisher, MA5-15739-HRP | Western Blotting | 1:100,000 |
| anti-N-cadherin | Mouse | BD Biosciences, 610920 | IF, Confocal | 1:500 |
| anti-Nestin | Mouse | R&D Systems, MAB1259 | IF, Confocal | 1:500 |
| anti-Sox2 | Rabbit | Millipore, AB5603 | IF, confocal | 1:500 |
| anti-Sox2 | Goat | R&D Systems, AF2018 | IF, Confocal | 1:300 |
| anti-VE Cadherin | Rabbit | Abcam, ab33168 | IF, Confocal | 1:500 |
| anti-CD31 | Mouse | Agilent Technologies, M082329-2 | IF, Confocal | 1:500 |
| anti-PAXILLIN | Mouse | Millipore, 05-41 | IF, TIRF | 1:200 |
| anti-KERATIN 14 | Rabbit | Biolegend, 905301 | IF, TIRF | 1:100 |
| anti-Rabbit IgG | Goat | Invitrogen, A32731 | IF | 1:500 |
| anti-Mouse IgG | Goat | Invitrogen, A32727 | IF | 1:500 |
| anti-Goat IgG | Donkey | Invitrogen, A21432 | IF | 1:500 |

#### Table of HaloTag Ligands

| **HaloTag Ligand** | **Catalog No.** | **Concentration** | **Incubation Time** |
| --- | --- | --- | --- |
| JF549 | Cat. No. GA1110, Promega | 500 pM | Overnight |
| JF635 | Requested from Janelia Materials, Luke Lavis Lab | 500 pM | Overnight |
| JF646 | Cat. No. GA1120, Promega | 500 pM | Overnight |
| JF646-BAPTA-3'AM | Requested from Janelia Materials, Luke Lavis Lab | 500 pM | 15 minutes |

#### Chemical Reagents Table

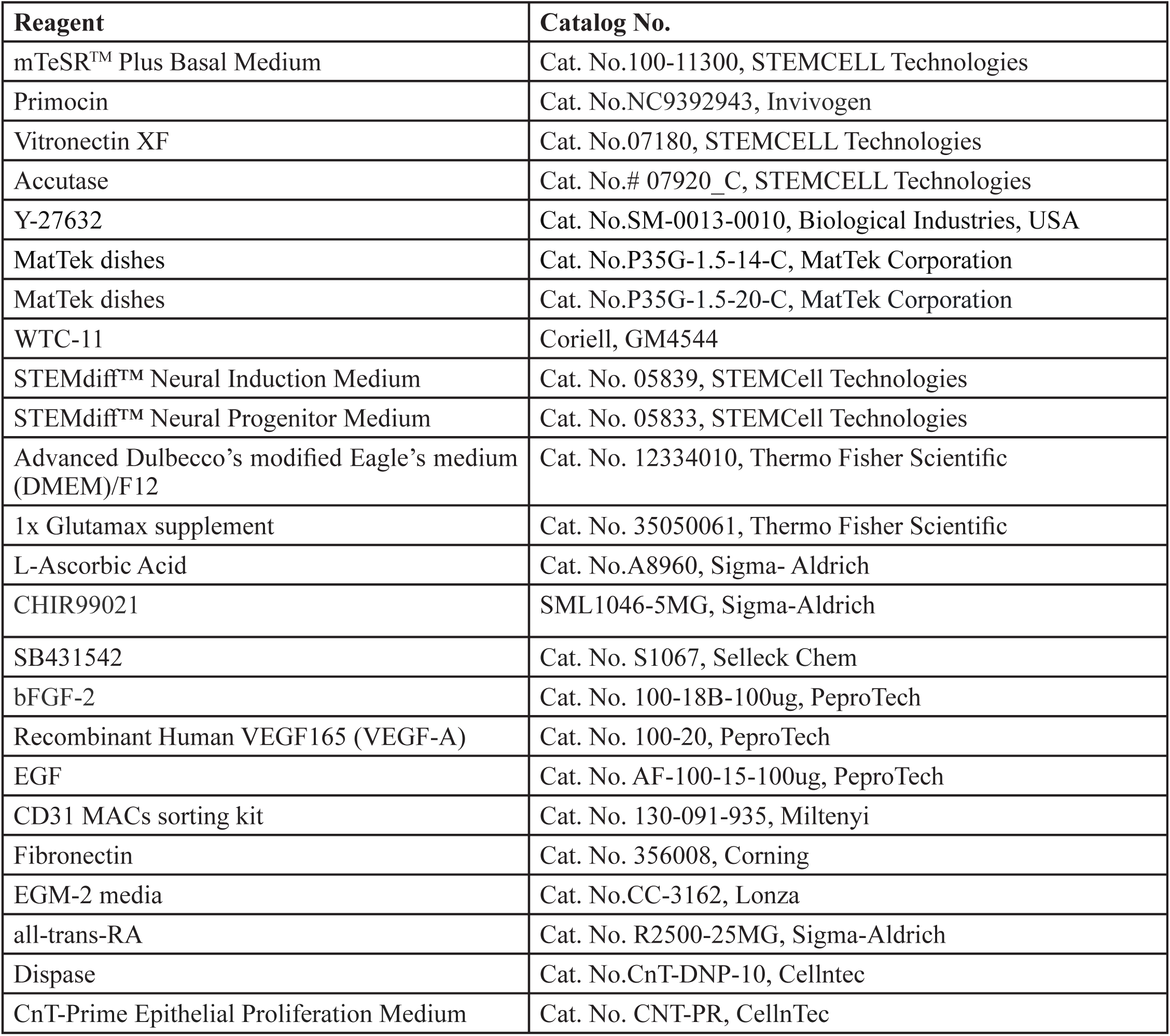
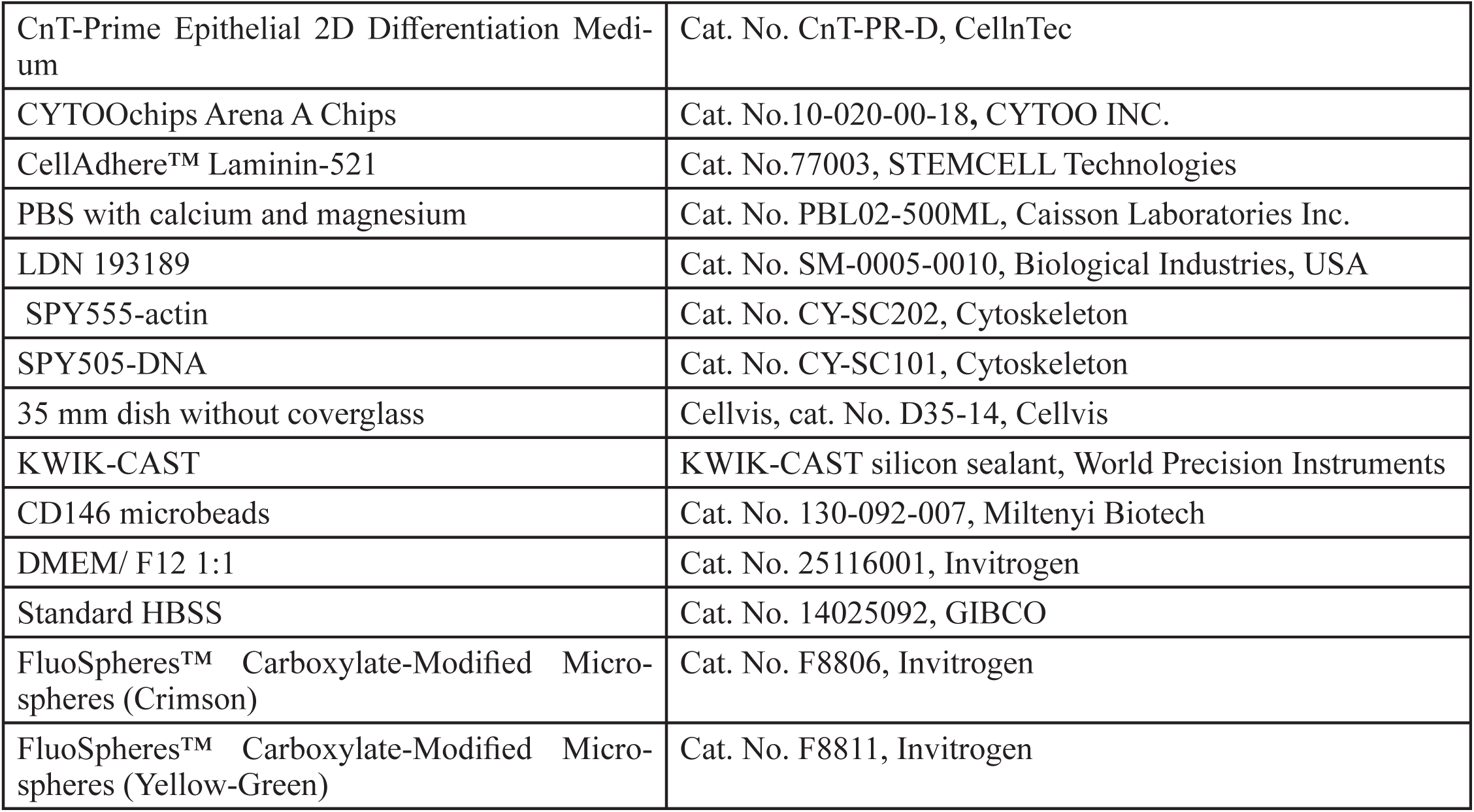

#### Table of Microscope information

| **Microscope** | **Objective information**<br>**(Magnification, Numerical Aperture)** | **Image Resolution Pixel Size**<br>**(µm/pixel)** | **Filter Cubes**<br>**Fluorescence**<br>**Excitation/ Emission** |
| --- | --- | --- | --- |
| Keyence bZ-X810 Widefield Microscope | 10x, NA 0.45<br>20x, NA 0.75<br>60x, NA 1.4 oil | 1.50977<br>0.75488<br>0.12581 | Ex 340-380/Em 435-485<br>Ex 465-495/Em 510 and up<br>Ex 527-553/Em 577-633<br>Ex 590-65/Em 662-737 |
| Olympus FV3000S laser-scanning confocal microscope | 40x, NA 1.25 | 0.3877 | U-FUW Ex 365/50/Em 420 LP<br>U-FBN Ex 482/25/Em 510 LP<br>UFGW Ex 540/20/Em 575 LP |
| Olympus IX83 CellTIRF | 60x, NA 1.45 | 0.1092 | Ex FF01-563/9-25<br>Em FF01-624/40-25<br>Ex FF01-640/14-25<br>Em<br>FF01-676/29-25 |
| Olympus IX83 CellTIRF | 60x, NA 1.50 | 0.1082 | Ex<br>ET-488/561nm Laser Dual TIRF Set for 486-492nm and 556-563nm lasers in BX3 cube<br>TRF49909-OL3; Single Bandpass ET-561nm laser TIRF set. For 557-563nm lasers. BX3 Cubes.<br>Em<br>E0431923; 455nm longpass cellTIRF Dichroic for 400-450nm lasers. Reflects: 400-450nm.<br>Transmits: 480-640nm.<br>E0431921; 525nm longpass cellTIRF Dichroic for 440-500nm lasers. Reflects: 440-500nm.<br>Transmits: 530-640nm.<br>E0431919; 595nm longpass cellTIRF Dichroic for 460-565nm lasers. Reflects: 460-565nm.<br>Transmits: 600-640nm. |
| AO-LLSM | 20x, NA 1.0 | 0.108 | CamA: Semrock FF01-600/ 37-25<br>CamB: Semrock FF01-538/685-25 |

## Data and Code Availability

Methods, representative movie files, and supplementary information are included in the manuscript. Source data for graphs in the main figures are available as a downloadable Supplementary Data file. Raw datasets, including source images and analyzed trajectories, included in this study can be requested from the corresponding author. No novel software tools were developed. Custom analysis code used has been deposited on Github at https://github.com/Pathak-Lab/PIEZO1-LocalizationTools and at https://github.com/abcucberkeley/piezo1_analysis_pipeline.

## Software used

- OriginPro 2022 (64-bit) SR1 9.9.0.225
- FLIKA version 4.8.1
- ImageJ 2.9.0/1.53t ; Java 1.8.0_322
- Micro-Manager 2.0
- cellTIRF 1.4
- Python 3.11.5
- Imaris x64: 10.0.0
- Amira 3D 2021.1
- MATLAB versions R2022b and R2023a

## References

1. Coste, B. et al. Piezo1 and Piezo2 are essential components of distinct mechanically activated cation channels. Science 330, 55–60 (2010).

2. Murthy, S. E., Dubin, A. E. & Patapoutian, A. Piezos thrive under pressure: mechanically activated ion channels in health and disease. Nat. Rev. Mol. Cell Biol. 18, 771–783 (2017).

3. Ranade, S. S. et al. Piezo1, a mechanically activated ion channel, is required for vascular development in mice. Proc. Natl. Acad. Sci. U. S. A. 111, 10347–10352 (2014).

4. Li, J. et al. Piezo1 integration of vascular architecture with physiological force. Nature 515, 279–282 (2014).

5. Rode, B. et al. Piezo1 channels sense whole body physical activity to reset cardiovascular homeostasis and enhance performance. Nat. Commun. 8, 1–11 (2017).

6. Cui, C.-P. et al. Piezo1 channel activation facilitates baroreflex afferent neurotransmission with subsequent blood pressure reduction in control and hypertension rats. Acta Pharmacol. Sin. 1–11 (2023).

7. Wang, S. et al. Endothelial cation channel PIEZO1 controls blood pressure by mediating flow-induced ATP release. J. Clin. Invest. 126, 4527 (2016).

8. Zeng, W. Z. et al. PIEZOs mediate neuronal sensing of blood pressure and the baroreceptor reflex. Science 362, (2018).

9. Cahalan, S. M. et al. Piezo1 links mechanical forces to red blood cell volume. Elife 4, (2015).

10. Faucherre, A., Kissa, K., Nargeot, J., Mangoni, M. E. & Jopling, C. Piezo1 plays a role in erythrocyte volume homeostasis. Haematologica 99, 70–75 (2014).

11. Pathak, M. M. et al. Stretch-activated ion channel Piezo1 directs lineage choice in human neural stem cells. Proc. Natl. Acad. Sci. U. S. A. 111, 16148–16153 (2014).

12. Nourse, J. L. et al. Piezo1 regulates cholesterol biosynthesis to influence neural stem cell fate during brain development. J. Gen. Physiol. 154, (2022).

13. Holt, J. R. et al. Spatiotemporal dynamics of PIEZO1 localization controls keratinocyte migration during wound healing. Elife 10, (2021).

14. Chen, J., Holt, J. R., Evans, E. L., Lowengrub, J. S., & Pathak, M. M. (2024). PIEZO1 regulates leader cell formation and cellular coordination during collective keratinocyte migration. PLoS Computational Biology, 20(4), e1011855.

15. Bae, C., Gnanasambandam, R., Nicolai, C., Sachs, F. & Gottlieb, P. A. Xerocytosis is caused by mutations that alter the kinetics of the mechanosensitive channel PIEZO1. Proc. Natl. Acad. Sci. U. S. A. 110, E1162– 8 (2013).

16. Zarychanski, R. et al. Mutations in the mechanotransduction protein PIEZO1 are associated with hereditary xerocytosis. Blood 120, 1908–1915 (2012).

17. Beneteau, C. et al. Recurrent mutation in the PIEZO1 gene in two families of hereditary xerocytosis with fetal hydrops. Clin. Genet. 85, 293–295 (2014).

18. Albuisson, J. et al. Dehydrated hereditary stomatocytosis linked to gain-of-function mutations in mechanically activated PIEZO1 ion channels. Nat. Commun. 4, 1884 (2013).

19. Lukacs, V. et al. Impaired PIEZO1 function in patients with a novel autosomal recessive congenital lymphatic dysplasia. Nat. Commun. 6, 8329 (2015).

20. Fotiou, E. et al. Novel mutations in PIEZO1 cause an autosomal recessive generalized lymphatic dysplasia with non-immune hydrops fetalis. Nat. Commun. 6, 8085 (2015).

21. Ma, S. et al. A role of PIEZO1 in iron metabolism in mice and humans. Cell 184, (2021).

22. Ma, S. et al. Common PIEZO1 Allele in African Populations Causes RBC Dehydration and Attenuates Plasmodium Infection. Cell 173, 443–455.e12 (2018).

23. Lacroix, J. J., Botello-Smith, W. M. & Luo, Y. Probing the gating mechanism of the mechanosensitive channel Piezo1 with the small molecule Yoda1. Nat. Commun. 9, 1–13 (2018).

24. Lewis, A. H. & Grandl, J. Mechanical sensitivity of Piezo1 ion channels can be tuned by cellular membrane tension. Elife 4, (2015).

25. Poole, K., Herget, R., Lapatsina, L., Ngo, H. D. & Lewin, G. R. Tuning Piezo ion channels to detect molecular-scale movements relevant for fine touch. Nat. Commun. 5, (2014).

26. Blythe, N. M. et al. Mechanically activated Piezo1 channels of cardiac fibroblasts stimulate p38 mitogen-activated protein kinase activity and interleukin-6 secretion. J. Biol. Chem. 294, 17395 (2019).

27. Jiang, F. et al. The mechanosensitive Piezo1 channel mediates heart mechano-chemo transduction. Nat. Commun. 12, 1–14 (2021).

28. Syeda, R. et al. Chemical activation of the mechanotransduction channel Piezo1. (2015) doi:10.7554/eLife.07369.

29. Wang, Y. et al. A lever-like transduction pathway for long-distance chemical- and mechano-gating of the mechanosensitive Piezo1 channel. Nat. Commun. 9, 1–12 (2018).

30. Ellefsen, K. L. et al. Myosin-II mediated traction forces evoke localized Piezo1-dependent Ca2+ flickers. Commun Biol 2, 298 (2019).

31. Richardson, J., Kotevski, A. & Poole, K. From stretch to deflection: the importance of context in the activation of mammalian, mechanically activated ion channels. FEBS J. 289, (2022).

32. Yao, M. et al. Forceand cell state-dependent recruitment of Piezo1 drives focal adhesion dynamics and calcium entry. Sci Adv 8, eabo1461 (2022).

33. Chen, X. et al. A Feedforward Mechanism Mediated by Mechanosensitive Ion Channel PIEZO1 and Tissue Mechanics Promotes Glioma Aggression. Neuron 100, 799–815.e7 (2018).

34. Chuntharpursat-Bon, E. et al. PIEZO1 and PECAM1 interact at cell-cell junctions and partner in endothelial force sensing. Commun Biol 6, 358 (2023).

35. Wang, J. et al. Tethering Piezo channels to the actin cytoskeleton for mechanogating via the cadherin-β-catenin mechanotransduction complex. Cell Rep. 38, 110342 (2022).

36. Gudipaty, S. A. et al. Mechanical stretch triggers rapid epithelial cell division through Piezo1. Nature 543, 118–121 (2017).

37. Yang, S. et al. Membrane curvature governs the distribution of Piezo1 in live cells. Nat. Commun. 13, 7467 (2022).

38. Ridone, P. et al. Disruption of membrane cholesterol organization impairs the activity of PIEZO1 channel clusters. J. Gen. Physiol. 152, (2020).

39. Ly, A. T., Freites, J. A., Bertaccini, G. A., Evans, E. L., Dickinson, G. D., Tobias, D. J., & Pathak, M. M. (2024). Single-particle tracking reveals heterogeneous PIEZO1 diffusion. In Biophysics (No. biorxiv;2022.09.30.510193v3). bioRxiv. https://www.biorxiv.org/content/10.1101/2022.09.30.510193v3

40. Lewis, A. H. & Grandl, J. Piezo1 ion channels inherently function as independent mechanotransducers. Elife 10, (2021).

41. Zheng, Q. et al. Mechanical properties of the brain: Focus on the essential role of Piezo1-mediated mechanotransduction in the CNS. Brain Behav. 13, e3136 (2023).

42. Baxter, S. L. et al. Investigation of associations between Piezo1 mechanoreceptor gain-of-function variants and glaucoma-related phenotypes in humans and mice. Sci. Rep. 10, 19013 (2020).

43. Grimm, J. B., Brown, T. A., English, B. P., Lionnet, T. & Lavis, L. D. Synthesis of Janelia Fluor HaloTag and SNAP-Tag Ligands and Their Use in Cellular Imaging Experiments. Methods Mol. Biol. 1663, 179– 188 (2017).

44. Deo, C., Sheu, S.-H., Seo, J., Clapham, D. E. & Lavis, L. D. Isomeric Tuning Yields Bright and Targetable Red Ca2+ Indicators. J. Am. Chem. Soc. 141, 13734–13738 (2019).

45. Grimm, J. B. et al. A general method to improve fluorophores for live-cell and single-molecule microscopy. Nat. Methods 12, 244–50, 3 p following 250 (2015).

46. Los, G. V. et al. HaloTag: a novel protein labeling technology for cell imaging and protein analysis. ACS Chem. Biol. 3, 373–382 (2008).

47. Romero, L. O. et al. Dietary fatty acids fine-tune Piezo1 mechanical response. Nat. Commun. 10, 1200 (2019).

48. Ovesný, M., Křížek, P., Borkovec, J., Svindrych, Z. & Hagen, G. M. ThunderSTORM: a comprehensive ImageJ plug-in for PALM and STORM data analysis and super-resolution imaging. Bioinformatics 30, 2389–2390 (2014).

49. Schindelin, J. et al. Fiji - an Open Source platform for biological image analysis. Nat. Methods 9,.

50. Ellefsen, K. L., Lock, J. T., Settle, B., Karsten, C. A. & Parker, I. Applications of FLIKA, a Python-based image processing and analysis platform, for studying local events of cellular calcium signaling. Biochim. Biophys. Acta Mol. Cell Res. 1866, 1171–1179 (2019).

51. Smith, I. F., Swaminathan, D., Dickinson, G. D. & Parker, I. Single-molecule tracking of inositol trisphosphate receptors reveals different motilities and distributions. Biophys. J. 107, 834–845 (2014).

52. Vaisey, G., Banerjee, P., North, A. J., Haselwandter, C. A. & MacKinnon, R. Piezo1 as a force-through-membrane sensor in red blood cells. Elife 11, (2022).

53. Pérez-Mitta, G., Sezgin, Y., Wang, W., & MacKinnon, R. (2024). Freestanding bilayer microscope for single-molecule imaging of membrane proteins. Science Advances, 10(25), eado4722.

54. Shuai, J. & Parker, I. Optical single-channel recording by imaging Ca2+ flux through individual ion channels: theoretical considerations and limits to resolution. Cell Calcium 37, 283–299 (2005).

55. Wijerathne, T. D., Ozkan, A. D. & Lacroix, J. J. Yoda1’s energetic footprint on Piezo1 channels and its modulation by voltage and temperature. Proc. Natl. Acad. Sci. U. S. A. 119, e2202269119 (2022).

56. Mulhall, E. M. et al. Direct observation of the conformational states of PIEZO1. Nature 620, 1117–1125 (2023).

57. Desplat, A. et al. Piezo1-Pannexin1 complex couples force detection to ATP secretion in cholangiocytes. J. Gen. Physiol. 153, (2021).

58. Syeda, R. et al. Piezo1 Channels Are Inherently Mechanosensitive. Cell Rep. 17, 1739–1746 (2016).

59. Knight, G. T. et al. Engineering induction of singular neural rosette emergence within hPSC-derived tissues. Elife 7, (2018).

60. Haremaki, T. et al. Self-organizing neuruloids model developmental aspects of Huntington’s disease in the ectodermal compartment. Nat. Biotechnol. 37, 1198–1208 (2019).

61. Liu, T.-L. et al. Observing the cell in its native state: Imaging subcellular dynamics in multicellular organisms. Science 360, (2018).

62. Zhou, Z. et al. MyoD-family inhibitor proteins act as auxiliary subunits of Piezo channels. Science 381, 799–804 (2023).

63. Buckley, D. L. et al. HaloPROTACS: Use of Small Molecule PROTACs to Induce Degradation of HaloTag Fusion Proteins. ACS Chem. Biol. 10, 1831 (2015).

64. Parker, I. & Smith, I. F. Recording single-channel activity of inositol trisphosphate receptors in intact cells with a microscope, not a patch clamp. J. Gen. Physiol. 136, 119–127 (2010).

65. Yaganoglu, S. et al. Highly specific and non-invasive imaging of Piezo1-dependent activity across scales using GenEPi. Nat. Commun. 14, 4352 (2023).

66. Gwosch, K. C. et al. MINFLUX nanoscopy delivers 3D multicolor nanometer resolution in cells. Nat. Methods 17, 217–224 (2020).

67. Verkest, C., Roettger, L., Zeitzschel, N. & Lechner, S. G. Cluster nanoarchitecture and structural diversity of PIEZO1 in intact cells. bioRxiv 2024.11.26.625366 (2024) doi:10.1101/2024.11.26.625366.

68. Xue, X. et al. Mechanics-guided embryonic patterning of neuroectoderm tissue from human pluripotent stem cells. Nat. Mater. 17, 633–641 (2018).

69. Zhou, Z., Li, J. V., Martinac, B. & Cox, C. D. Loss-of-Function Piezo1 Mutations Display Altered Stability Driven by Ubiquitination and Proteasomal Degradation. Front. Pharmacol. 12, 766416 (2021).

70. Wang, K. et al. Robust differentiation of human pluripotent stem cells into endothelial cells via temporal modulation of ETV2 with modified mRNA. Science Advances (2020) doi:10.1126/sciadv.aba7606.

71. Grimm, J. B. et al. A general method to fine-tune fluorophores for live-cell and in vivo imaging. Nat. Methods 14, 987–994 (2017).

72. Edelstein, A., Amodaj, N., Hoover, K., Vale, R. & Stuurman, N. Computer control of microscopes using µManager. Curr. Protoc. Mol. Biol. Chapter 14, Unit14.20 (2010).

73. Frei, M. S. et al. Engineered HaloTag variants for fluorescence lifetime multiplexing. Nat. Methods 19, 65–70 (2022).

74. Liu, G. et al. Characterization, comparison, and optimization of lattice light sheets. Sci Adv 9, eade6623 (2023).

75. Wong, K. K. L. et al. Origin of wiring specificity in an olfactory map revealed by neuron type–specific, time-lapse imaging of dendrite targeting. (2023) doi:10.7554/eLife.85521.

76. Weigert, M. et al. Content-aware image restoration: pushing the limits of fluorescence microscopy. Nat. Methods 15, 1090–1097 (2018).

77. Aguet, F. et al. Membrane dynamics of dividing cells imaged by lattice light-sheet microscopy. Mol. Biol. Cell 27, 3418–3435 (2016).

78. Ho, J., Tumkaya, T., Aryal, S., Choi, H. & Claridge-Chang, A. Moving beyond P values: data analysis with estimation graphics. Nat. Methods 16, 565–566 (2019).

