## Supplemental Information for "Visualizing PIEZO1 Localization and Activity in hiPSC-Derived Single Cells and Organoids with HaloTag Technology"

#### Supplementary Information

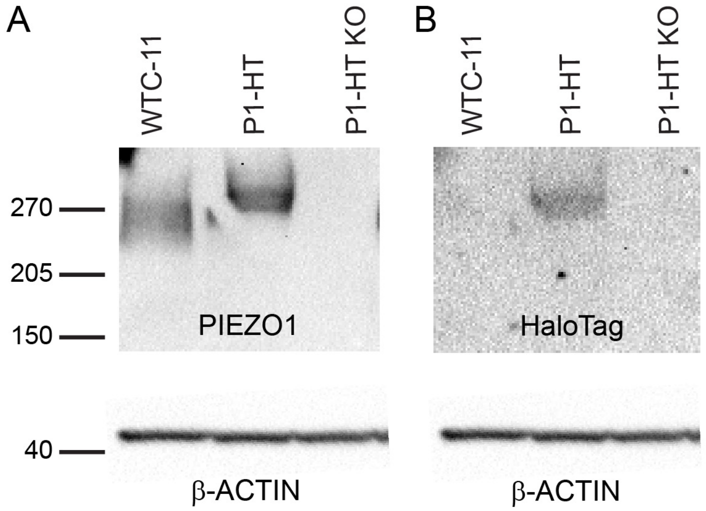


**Supplemental Figure 1. Western blots demonstrate successful PIEZO1-HaloTag fusion.** Immunoblot of hiPSCs (upper panels) probed with anti-PIEZO1 **(A)** or anti-HaloTag **(B)** antibody. The untagged PIEZO1 from WTC-11 hiPSCs (predicted molecular weight 289 kDa), PIEZO1-HaloTag (P1-HT, predicted molecular weight 319 kDa) migrated at the expected sizes based on molecular weight standards with no signal in PIEZO1-HaloTag Knockout (P1-HT KO) lanes. (Lower panels) Immunoblot probed with anti-β-ACTIN demonstrates equal loading of samples. n = 4 independent repeats.

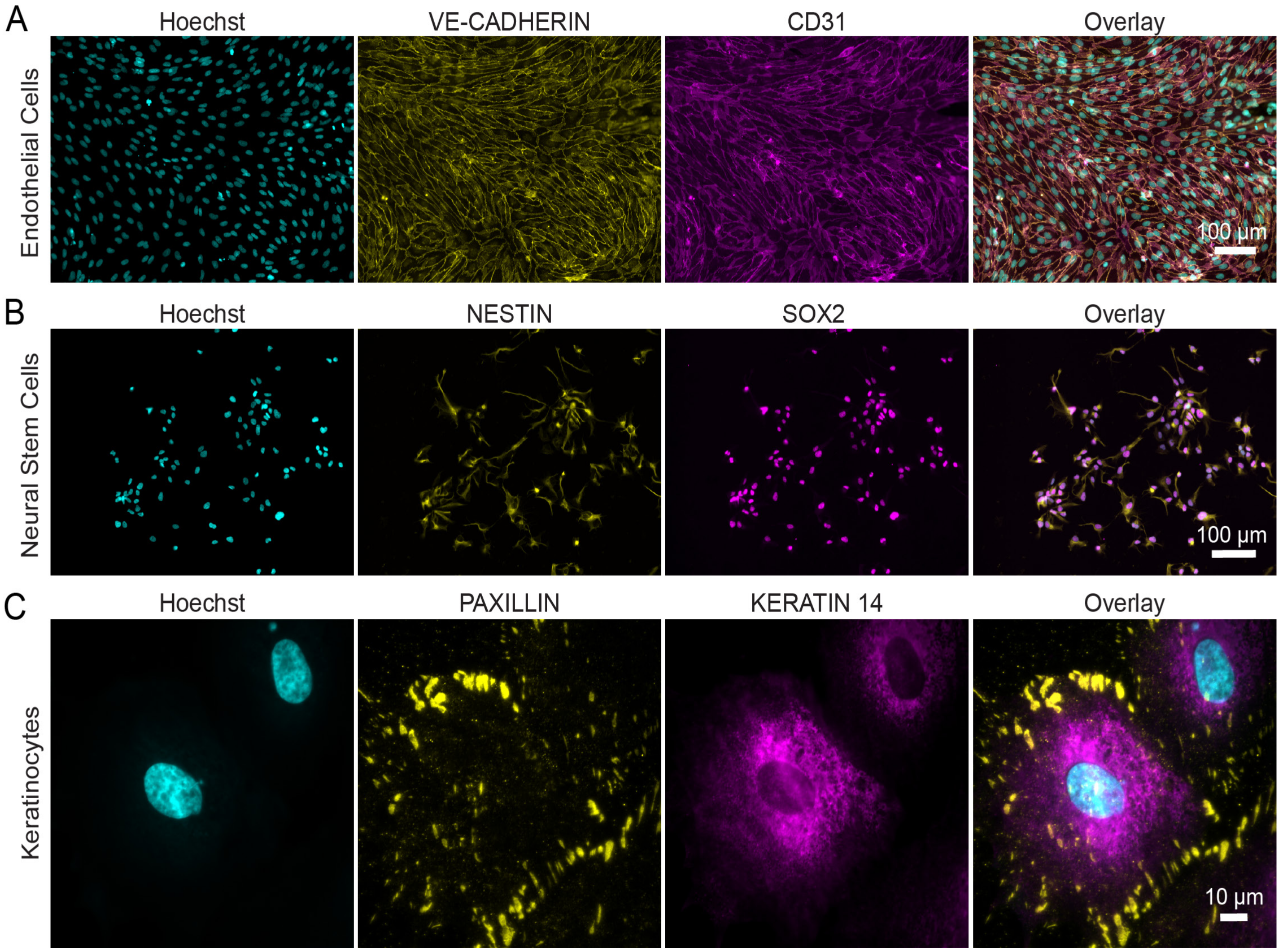


**Supplemental Figure 2. Validation of endothelial cells, neural stem cells, and keratinocytes differentiated from PIEZO1-HaloTag hiPSCs. A.** Endothelial cells differentiated from PIEZO1-HaloTag hiSPCs expressed endothelial cell markers VE-CADHERIN (Yellow) and CD31 (Magenta). **B.** Neural stem cells differentiated from PIEZO1-HaloTag hiPSCs expressed neural stem cell markers NESTIN (Yellow) and SOX2 (Magenta). **C.** Keratinocytes differentiated from PIEZO1-HaloTag hiPSCs expressed keratinocyte marker KERATIN 14 (Magenta) and displayed prominent focal adhesion marker PAXILLIN (Yellow). See the Methods section for differentiation protocols.

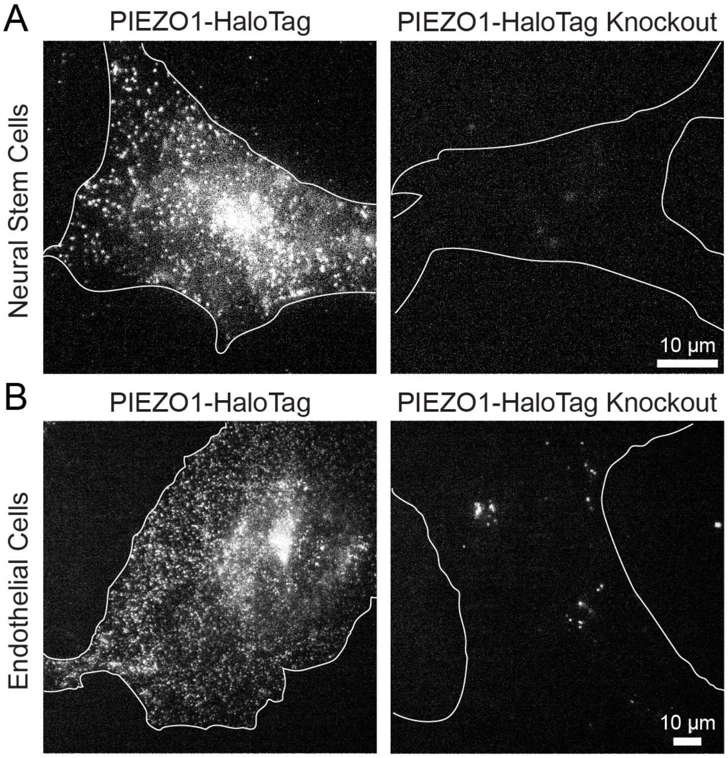


**Supplemental Figure 3. Specific labeling of PIEZO1-HaloTag channels. A.** Differentiated PIEZO1-HaloTag and PIEZO1-HaloTag Knockout neural stem cells labeled with JF646 HTL and imaged using TIRF. **B.** Differentiated PIEZO1-HaloTag and PIEZO1-HaloTag Knockout endothelial cells labeled with JF646 HTL and imaged using TIRF. Note the lack of HTL signal in Knockout cells.

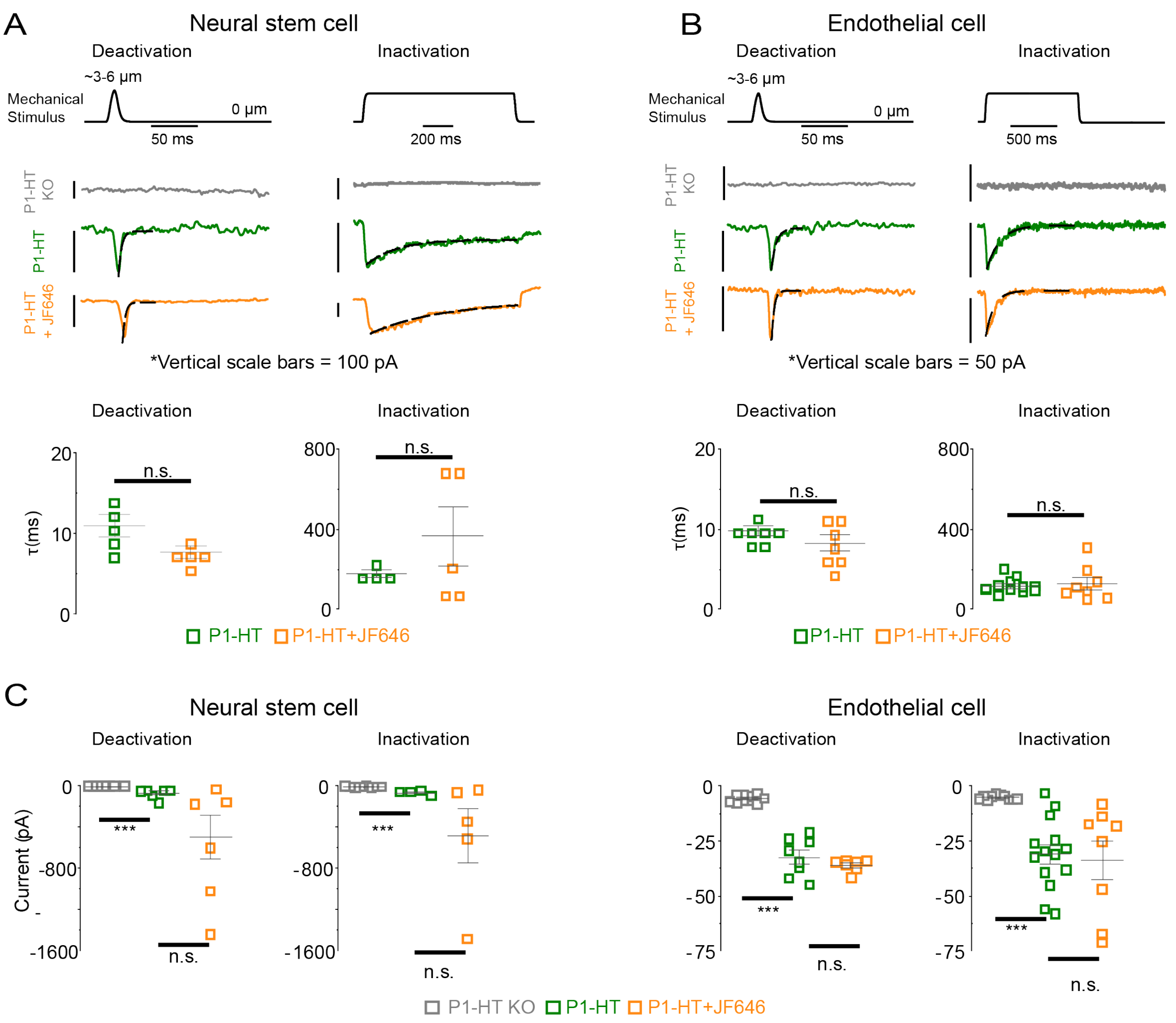


**Supplemental Figure 4. Attachment of HaloTag ligand does not affect PIEZO1-HaloTag channel function.** Whole-cell poking assays on PIEZO1-HaloTag Knockout (grey), PIEZO1-HaloTag (green), and PIEZO1-HaloTag + JF646 HTL (orange) from **A.** differentiated neural stem cells ( $\tau_{deactivation}$ p-value: 0.07, $\tau_{inactivation}$ p-value 0.30) and **B.** differentiated endothelial cells ($\tau_{deactivation}$ p-value: 0.21, $\tau_{inactivation}$ p-value 0.68). n.s. denotes no statistically significant difference between P1-HT and P1-HT + JF646. All values are expressed as mean ± SEM (NSC P1-HT deactivation mean:10.96±1.35 ms, n=5; NSC P1-HT+JF646 deactivation mean:7.71±0.75 ms, n=5; NSC P1-HT inactivation mean:178.66±18.62 ms, n=4; NSC P1-HT+-JF646 inactivation mean: 363.69±146.63 ms, n=5; EC P1-HT deactivation mean: 9.82±0.62 ms, n=7; EC P1-HT+JF646 deactivation mean: 8.28±0.99 ms, n=7; EC P1-HT inactivation mean:155.85±10.70 ms, n=12; EC P1-HT+JF646 inactivation mean:127.60±30.46 ms, n=8.) **C.** Maximal poking-evoked current amplitudes recorded from P1-HT KO (grey), P1-HT (green), and P1-HT + JF646 HTL (orange) conditions of NSC and EC. All values are expressed as mean ± SEM (NSC P1-KO deactivation mean: -3.10±0.20 pA, n=8; NSC P1-HT deactivation mean:-70.32±17.86 pA, n=7; NSC P1-HT+JF646 deactivation mean:-496.66±207.73 pA, n=7; NSC P1-KO inactivation mean:-9.83±1.62 pA, n=8; NSC P1-HT inactivation mean:-64.91±10.76 pA, n=4; NSC P1-HT+JF646 inactivation mean:-488.06±464.10 pA, n=5; EC P1-KO deactivation mean:-5.87±0.53 pA, n=10; EC P1-HT deactivation mean: -32.39±3.08 pA, n=8; EC P1-HT+JF646 deactivation mean: -36.12±1.10 pA, n=7; EC P1-KO inactivation mean:-5.29±0.32 pA, n=10; EC P1-HT inactivation mean:-31.0±4.27 pA, n=14; EC P1-HT+JF646 inactivation mean:and -33.62±8.79 pA, n=8.) p-values were calculated using unpaired two-sample t-Tests. (*** p-value < 0.005)

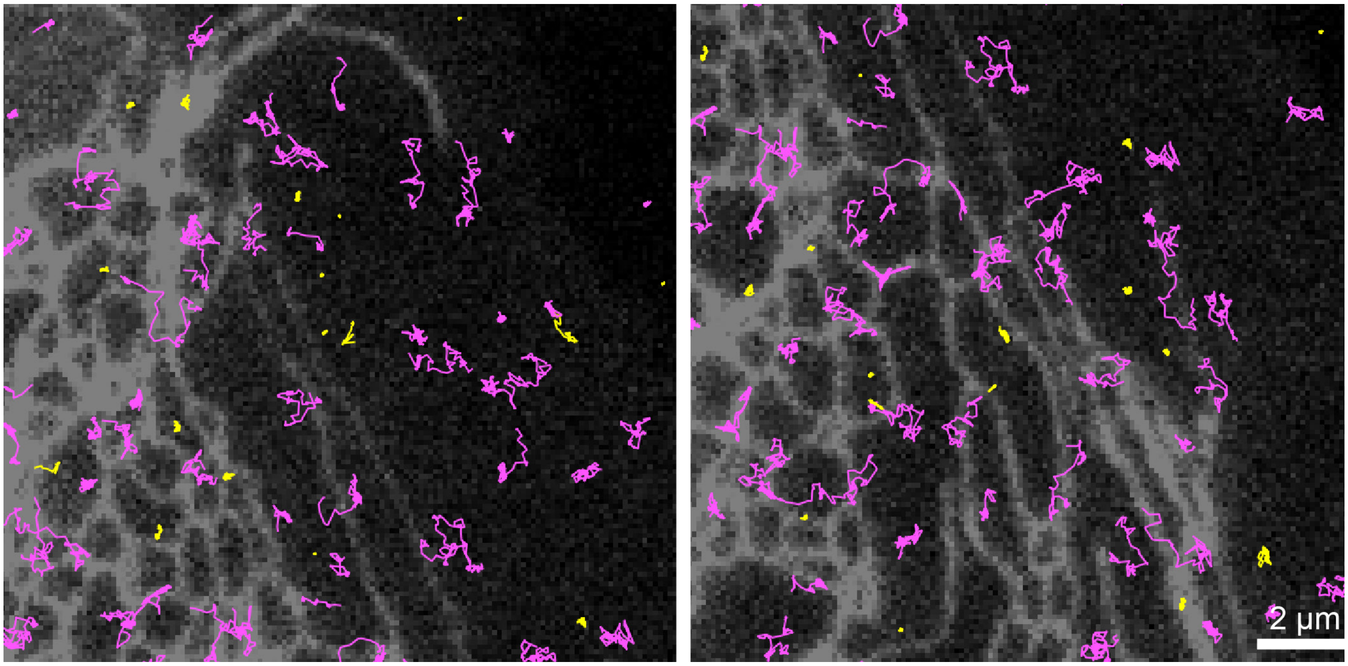


**Supplemental Figure 5. Motility of PIEZO puncta is not determined by association with the ER.** Panels show representative TIRF images of multiple positions within PIEZO1-HaloTag endothelial cells labeled with ER tracker 488 dye (grayscale) overlaid with trajectories of JF646 HTL PIEZO1-HaloTag puncta. Trajectories are color-coded by path length as in Fig. 2 (magenta: mobile, yellow: immobile). Some mobile and immobile trajectories overlap with the ER signal while others localize to ER-free regions.

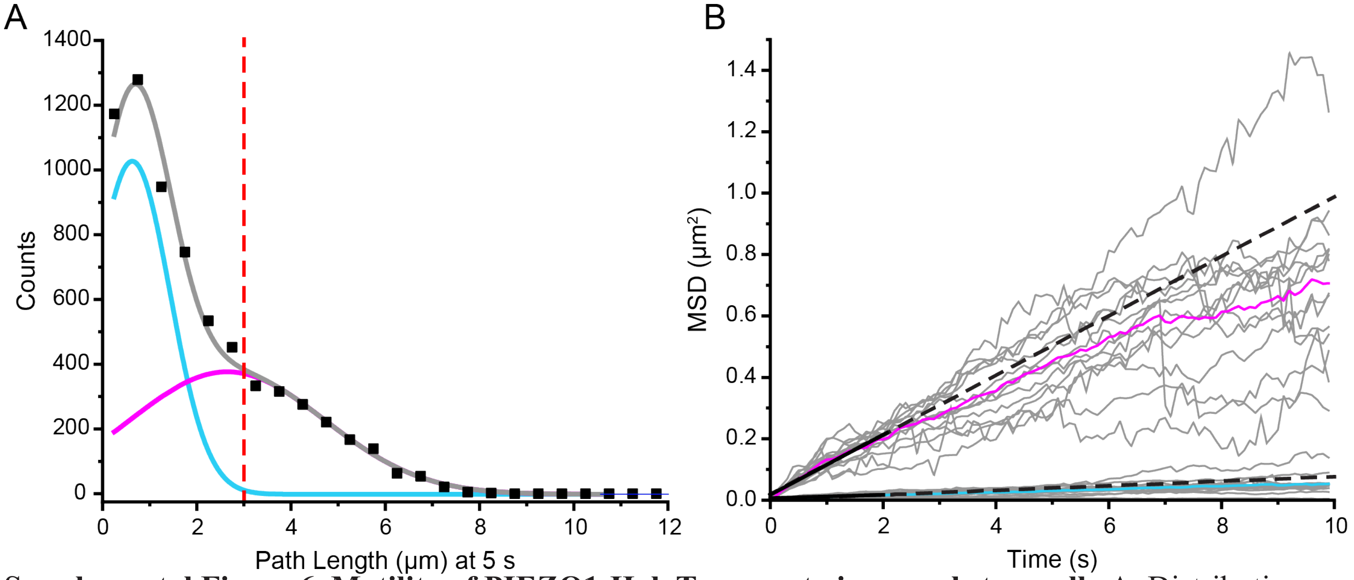


**Supplemental Figure 6. Motility of PIEZO1-HaloTag puncta in neural stem cell. A.** Distribution of trajectory path lengths for tracks at 5 s derived from live PIEZO1-HaloTag neural stem cells labeled with JF646 HTL (n = 15 videos from 3 independent experiments). Grey curve represents a fit to a sum of two Gaussian curves; the individual Gaussian curves are shown in cyan and magenta. **B**. Mean squared displacement (MSD) of immobile (cyan) and mobile puncta (magenta). Tracks that had a path length > 3 µm were characterized as mobile (magenta) and tracks which had a path length < 3 µm were characterized as immobile (cyan) in the Gaussian curve. Each gray trace represents mean MSD for all mobile puncta in a video (upper traces) and for all immobile puncta in a video (lower traces); data from 15 videos from 3 independent experiments are plotted. Solid magenta (mobile) and cyan (immobile) curves are mean MSD curves across all videos. Solid black line is a linear fit to the initial MSD t < 2 s, dashed black line shows a linear extrapolation.

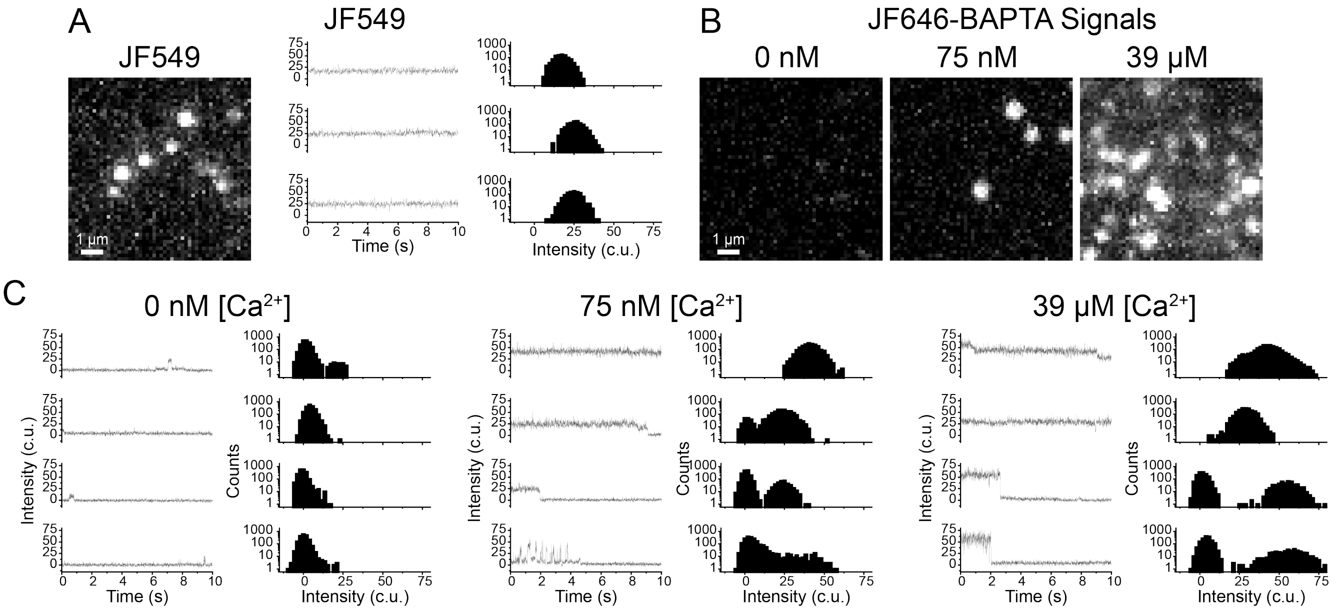


**Supplemental Figure 7. Fluorescence signals from monomeric cytoplasmic HaloTag imaged at different free $Ca^{2+}$ concentrations.** WTC-11 hiPSCs were transfected with plasmids expressing cytoplasmic HaloTag and then labeled with a 1:1 mixture of JF549 and JF646-BAPTA. Labeled cells were fixed, permeabilized and imaged for 10 s at 200fps. **A.** Representative TIRF maximum intensity projection image of JF549 HTL (left). Fluorescence intensities over time for three representative JF549 puncta along with all-point amplitude histograms (right). Note the stable fluorescence intensity over time. **B.** Fixed and permeabilized samples were exposed to increasing levels of free $Ca^{2+}$. Representative TIRF maximum intensity projection images of JF646-BAPTA HTL at different $Ca^{2+}$ concentrations. **C.** Fluorescence intensities over time for four representative JF646-BAPTA puncta for different $Ca^{2+}$ concentrations, along with all-point amplitude histograms. Note the increased occupancy at higher fluorescence intensity with increase in $Ca^{2+}$.

Supp. Fig.8

### Untreated

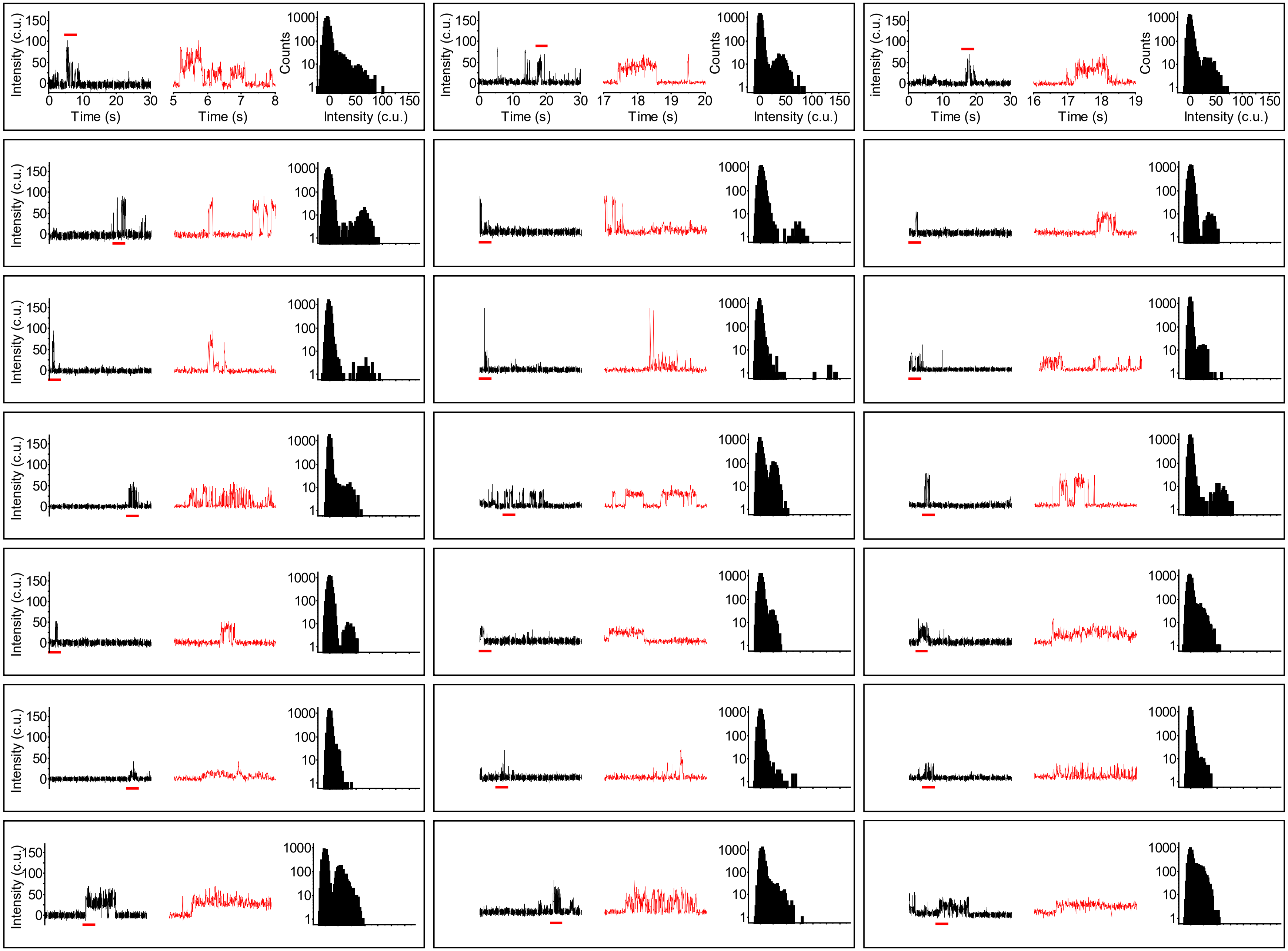

**Supplemental Figure 8. Representative activity traces from untreated PIEZO1-HaloTag.** Representative background-subtracted fluorescence intensity traces of 21 untreated immobile PIEZO1-HaloTag puncta from 3 independent TIRF imaging experiments of hiPSC-derived endothelial cells labeled with the $Ca^{2+}$-sensitive JF646-BAPTA HTL. Images were acquired at a frame rate of 200 fps. Intensity profiles were plotted from tracked immobile puncta. All traces are 30 s long (black) with a zoom in of a 3-s portion (red). Rightmost panels show an all-points histogram of intensity levels for the entirety of the 30-s recording.

Supp. Fig.9

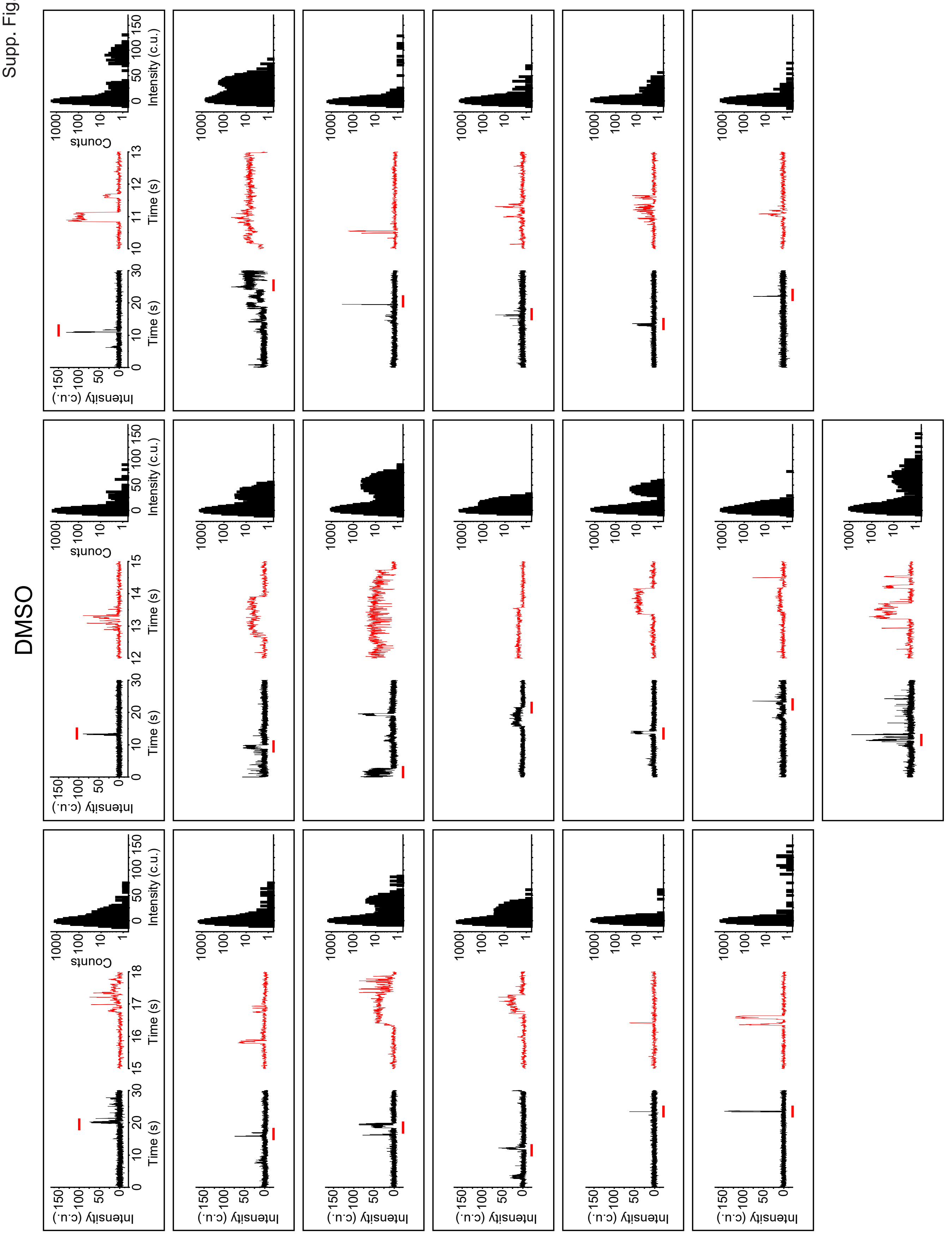

**Supplemental Figure 9. Representative activity traces from PIEZO1-HaloTag treated with vehicle control, DMSO.** Representative background-subtracted fluorescence intensity traces of 21 immobile PIEZO1-HaloTag puncta from 3 independent TIRF imaging experiments of hiPSC-derived endothelial cells labeled with the $Ca^{2+}$-sensitive JF646-BAPTA HTL and treated with vehicle control, DMSO. Intensity profiles were plotted from tracked immobile puncta. All traces are 30 s long (black) with a zoom in of a 3-s portion (red). Rightmost panels show an all-points histogram of intensity levels for the entirety of the 30-s recording.

Supp. Fig.10

### 2 µM Yoda1

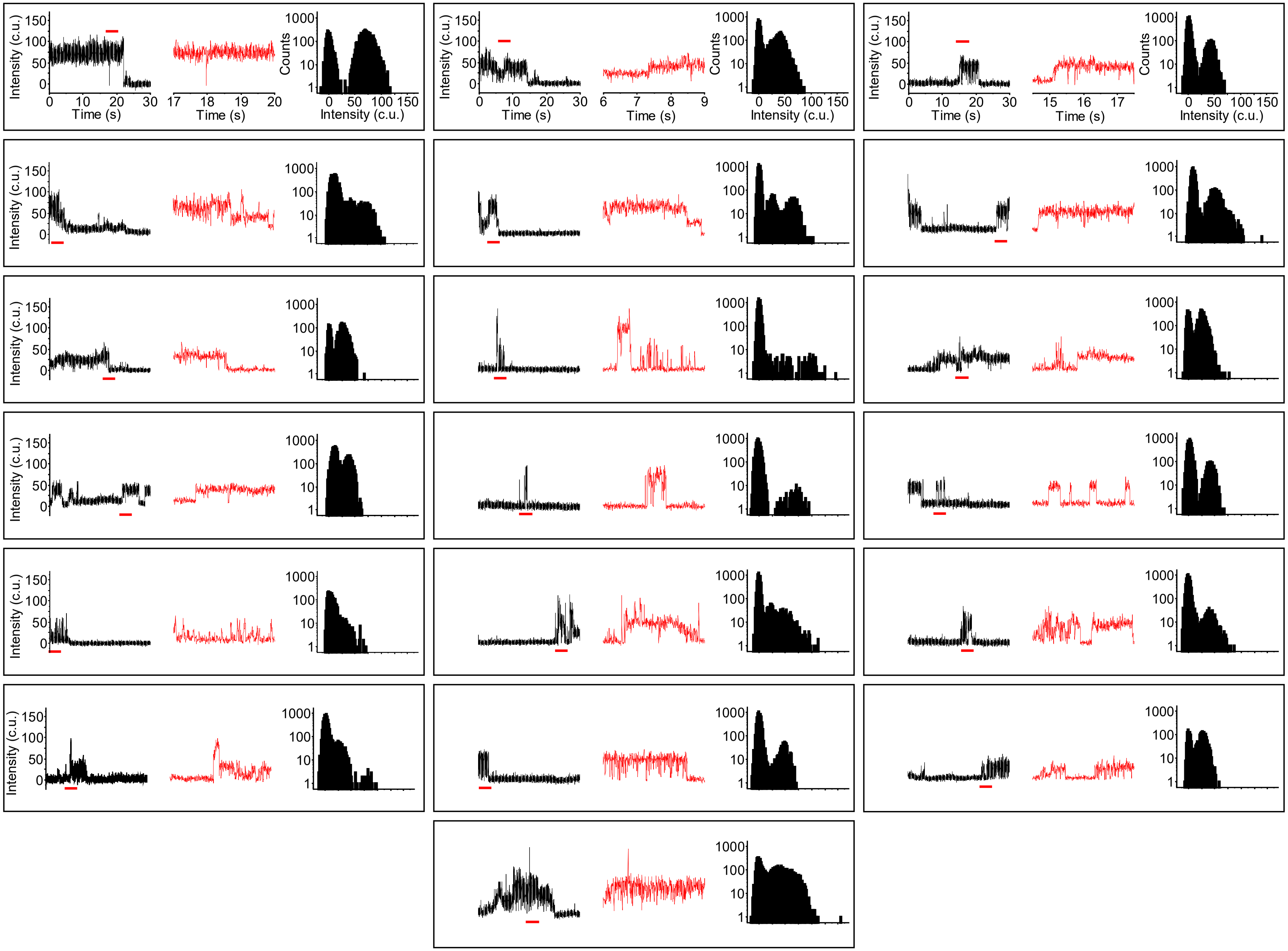

**Supplemental Figure 10. Representative activity traces from PIEZO1-HaloTag treated with 2 µM of Yoda1.** Representative background-subtracted fluorescence intensity traces of 21 immobile PIEZO1-HaloTag puncta from 3 independent TIRF imaging experiments of hiPSC-derived endothelial cells labeled with the $Ca^{2+}$-sensitive JF646-BAPTA HTL and treated with 2 µM Yoda1. Images were acquired at a frame rate of 200 fps. Intensity profiles were plotted from tracked immobile puncta. All traces are 30 s long (black) with a zoom in of a 3-s portion (red). Rightmost panels show an all-points histogram of intensity levels for the entirety of the 30-s recording.

Supp. Fig.11, page 1

### Endothelial cells 500 fps

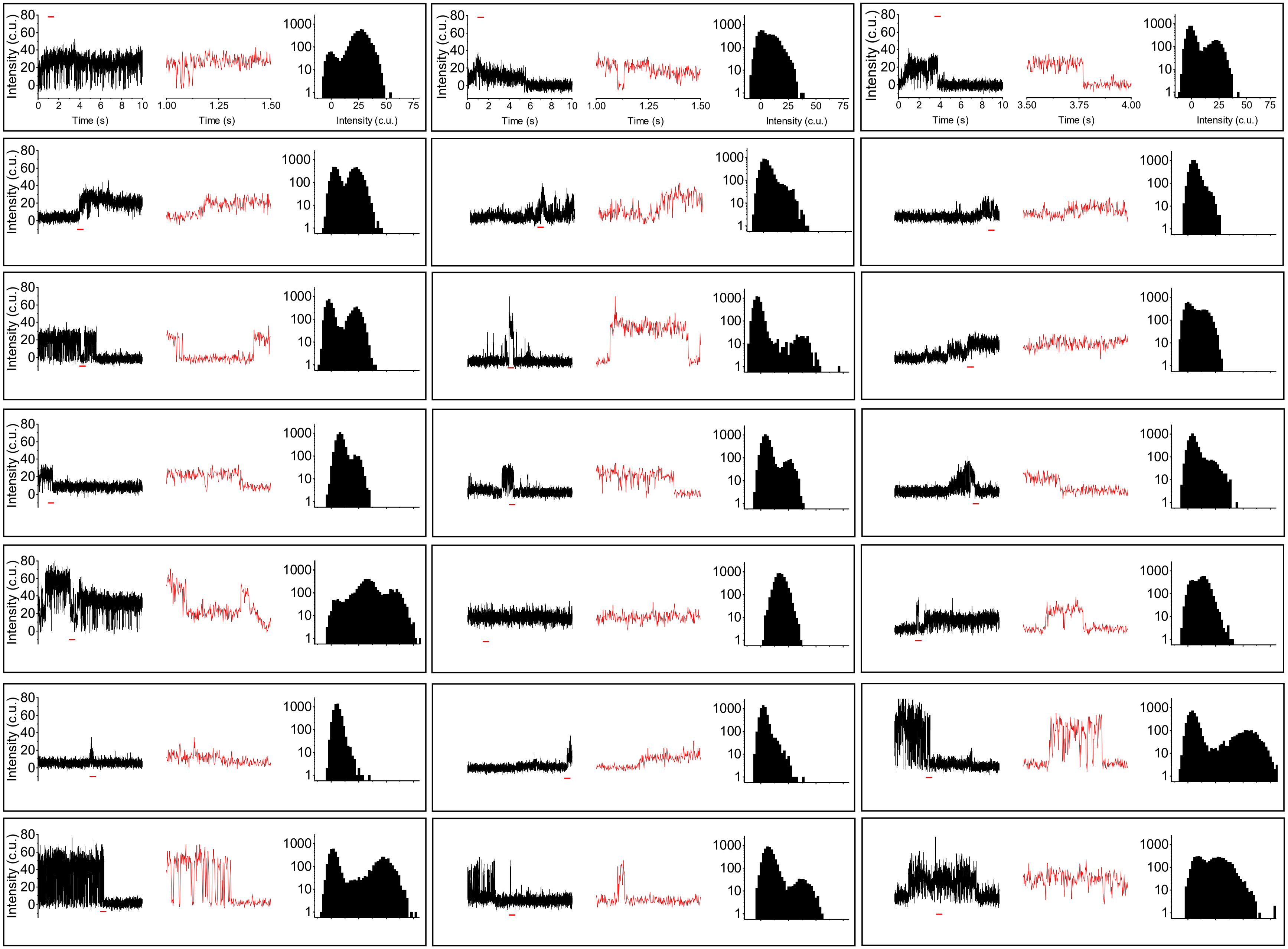

### Endothelial cells 500 fps

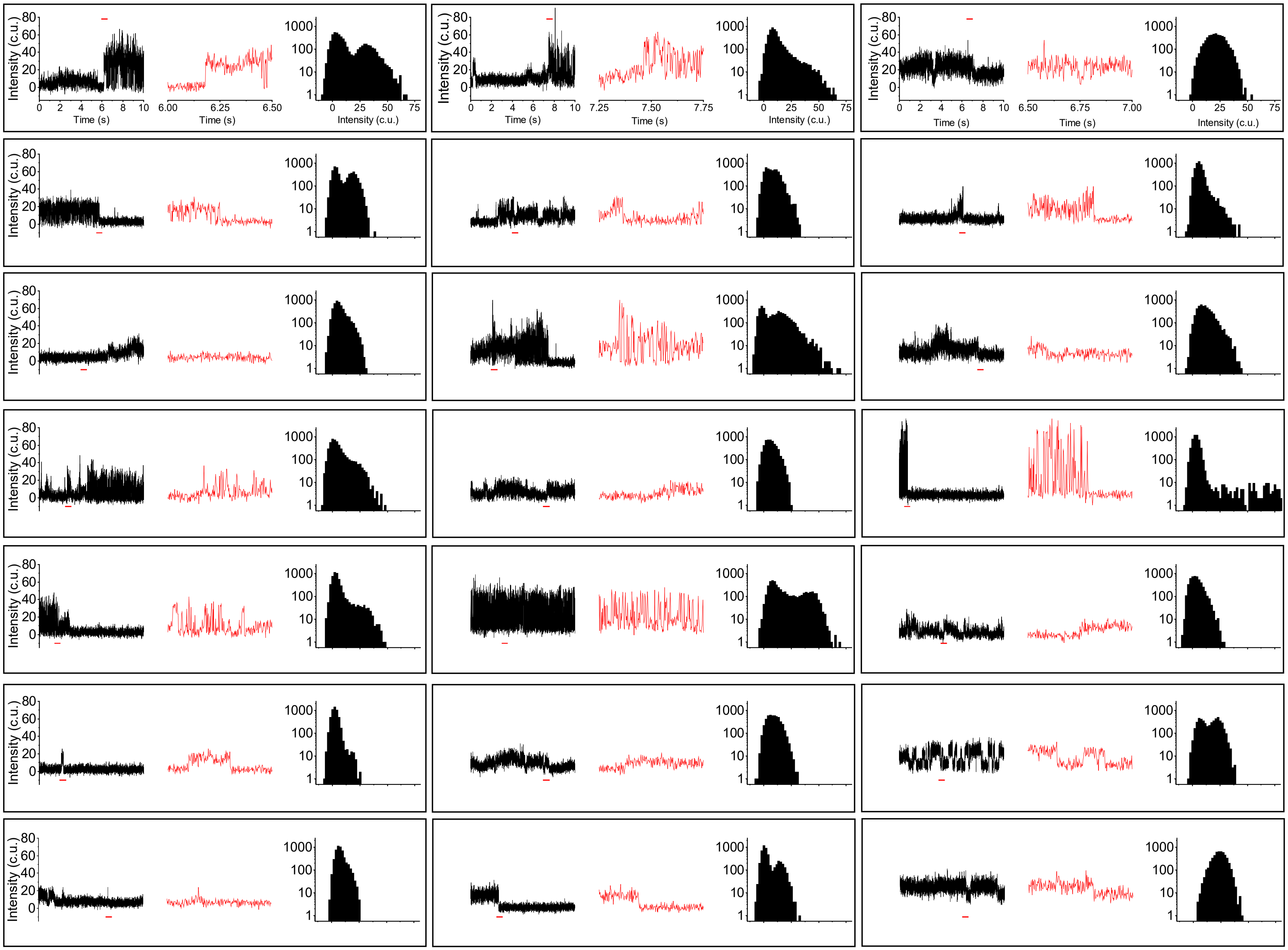

### Endothelial cells 500 fps

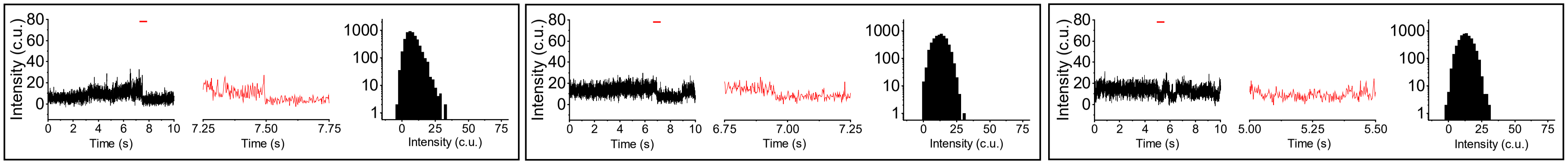

**Supplemental Figure 11. Representative 500-fps activity traces from PIEZO1-HaloTag endothelial cells.** Representative background-subtracted fluorescence intensity traces of 45 immobile PIEZO1-HaloTag puncta from TIRF imaging of hiPSC-derived endothelial cells labeled with the $Ca^{2+}$-sensitive JF646-BAPTA HTL. Intensity profiles were plotted from tracked immobile puncta. All traces shown are 10 seconds long (black) with a zoom in of a 0.5-second portion (red). Rightmost panels show an all-points histogram of intensity levels for the entirety of the 10-second recording.

### Neural stem cells 500 fps

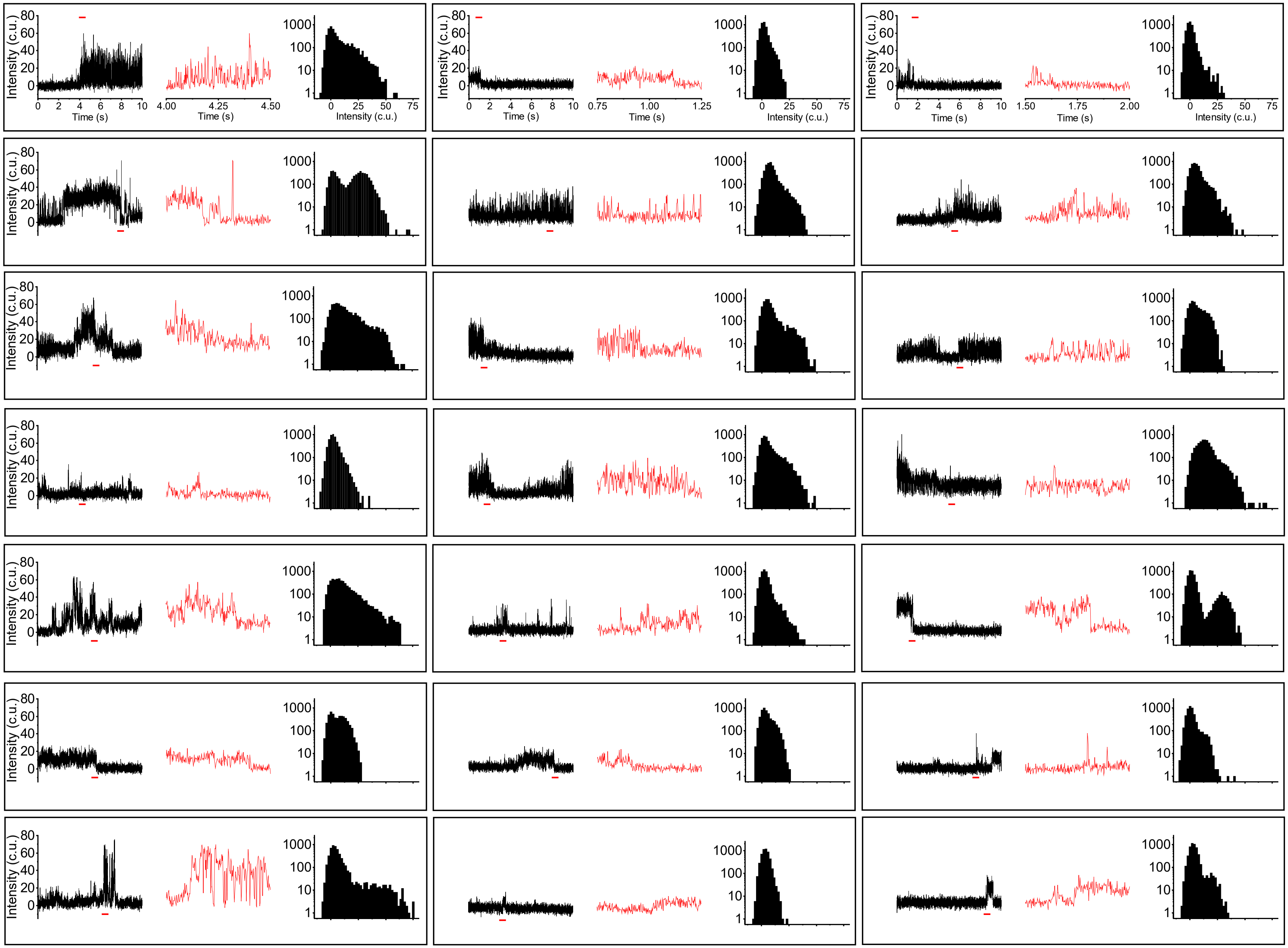

Supp. Fig.12, page 2

### Neural stem cells 500 fps

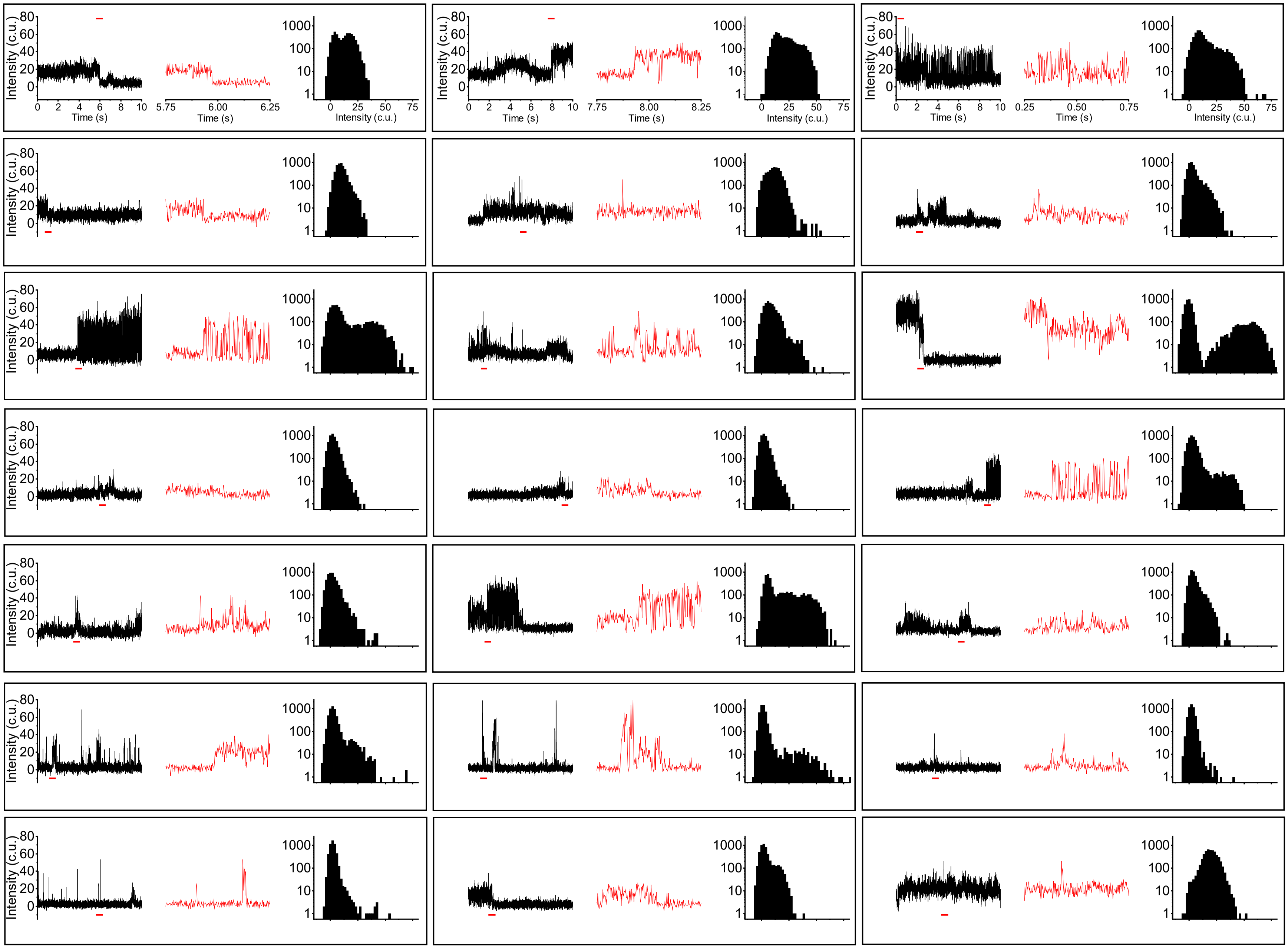

### Neural stem cells 500 fps

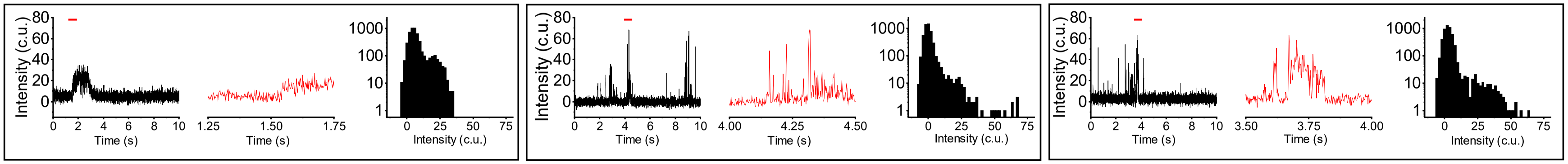

**Supplemental Figure 12. Representative 500-fps activity traces from PIEZO1-HaloTag neural stem cells.** Representative background-subtracted fluorescence intensity traces of 45 immobile PIEZO1-HaloTag puncta from TIRF imaging of hiPSC-derived neural stem cells labeled with the $Ca^{2+}$-sensitive JF646-BAPTA HTL. Intensity profiles were plotted from tracked immobile puncta. All traces shown are 10 seconds long (black) with a zoom in of a 0.5-second portion (red). Rightmost panels show an all-points histogram of intensity levels for the entirety of the 10-second recording.

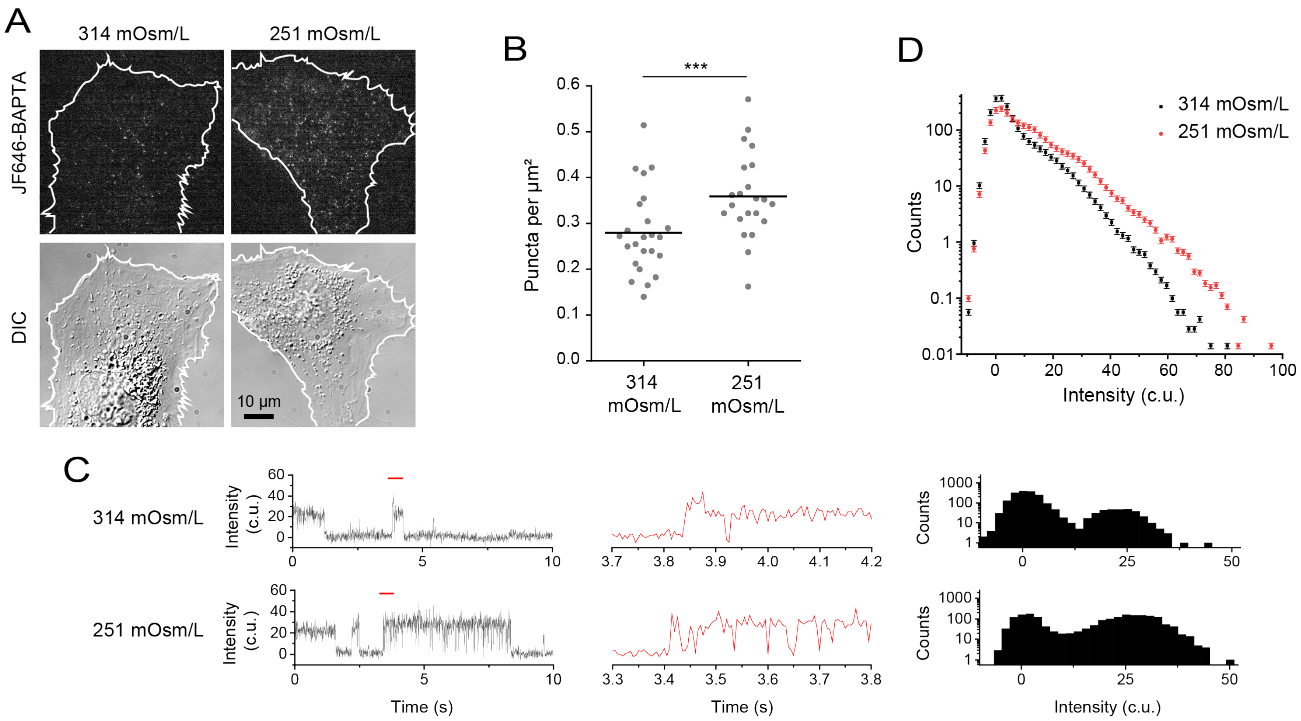


**Supplemental Figure 13. Hypotonic stimulus increases PIEZO1 activity. A.** Representative TIRF images of $Ca^{2+}$-sensitive HaloTag Ligand JF646-BAPTA puncta (top) and corresponding differential interference contrast (DIC) images (bottom) of hiPSC-derived PIEZO1-HaloTag endothelial cells, in control solution (314 mOsm/L) or under hypotonic stimulus (251 mOsm/L). **B**. Quantitation of puncta density of JF646-BAPTA HTL-labeled PIEZO1 puncta (see Methods). Each dot is the puncta density in one cell (n = 22 to 24 cells from 3 independent experiments). 314 mOsm/L mean puncta density: 0.28 ± 0.02 puncta per $\mu m^2$. 251 mOsm/L mean puncta density: 0.36 ± 0.02 puncta per $\mu m^2$. Means are indicated by black lines. The two groups were significantly different from one another (p-value Mann-Whitney < 0.005). The Cohen's d effect size between control and hypotonic conditions was 0.86. **C.** Representative background-subtracted fluorescence intensity traces of immobile PIEZO1-HaloTag puncta from 200-fps TIRF imaging of ECs imaged in 314 or 251 mOsm/L solutions, labeled with JF646-BAPTA HTL. *Center*, expanded traces from the sections marked with a red line on the left. *Right,* corresponding all-points amplitude histograms of intensity levels from the 10 s traces on the left. Counts are shown on a log10 scale. **D.** An average all-points amplitude histogram from 71 immobile puncta representing 3 independent experiments of each condition (black: 314 mOsm/L, red: 251 mOsm/L). For individual fluorescence traces and all-points amplitude histogram of each punctum, please see Supplemental Fig. 13-14.

Supp. Fig.14

Control (314 mOsm/L) 200 fps

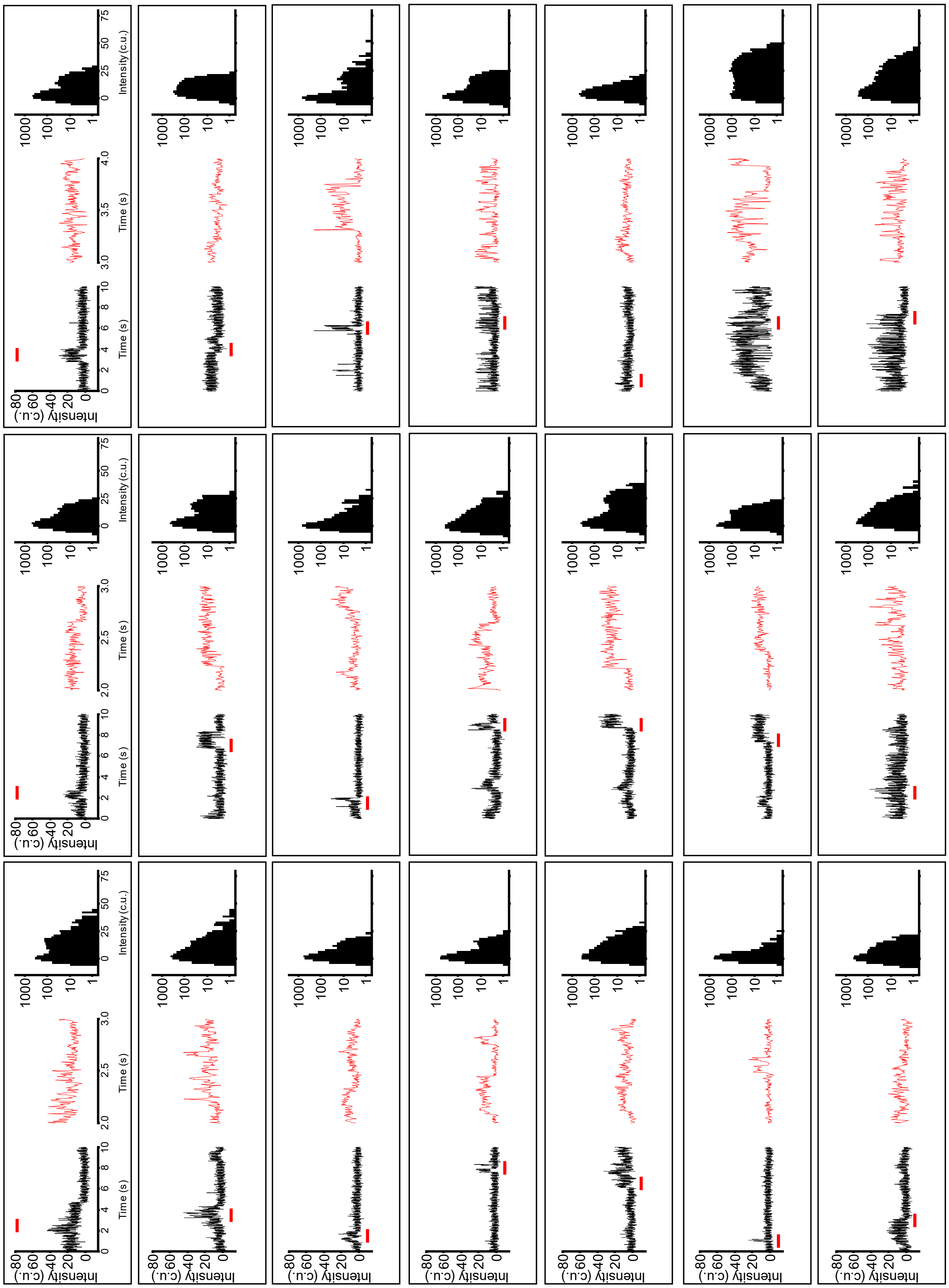

**Supplemental Figure 14. Representative 200-fps activity traces from PIEZO1-HaloTag endothelial cells in control solution (314 mOsm/L).** Representative background-subtracted fluorescence intensity traces of 21 immobile PIEZO1-HaloTag puncta from TIRF imaging of hiPSC-derived endothelial cells labeled with the $Ca^{2+}$-sensitive JF646-BAPTA HTL. Intensity profiles were plotted from tracked immobile puncta. All traces shown are 10 seconds long (black) with a zoom in of a 1-second portion (red). Right-most panels show an all-points histogram of intensity levels for the entirety of the 10-second recording.

Supp. Fig.15

### Hypotonic (251 mOsm/L) 200 fps

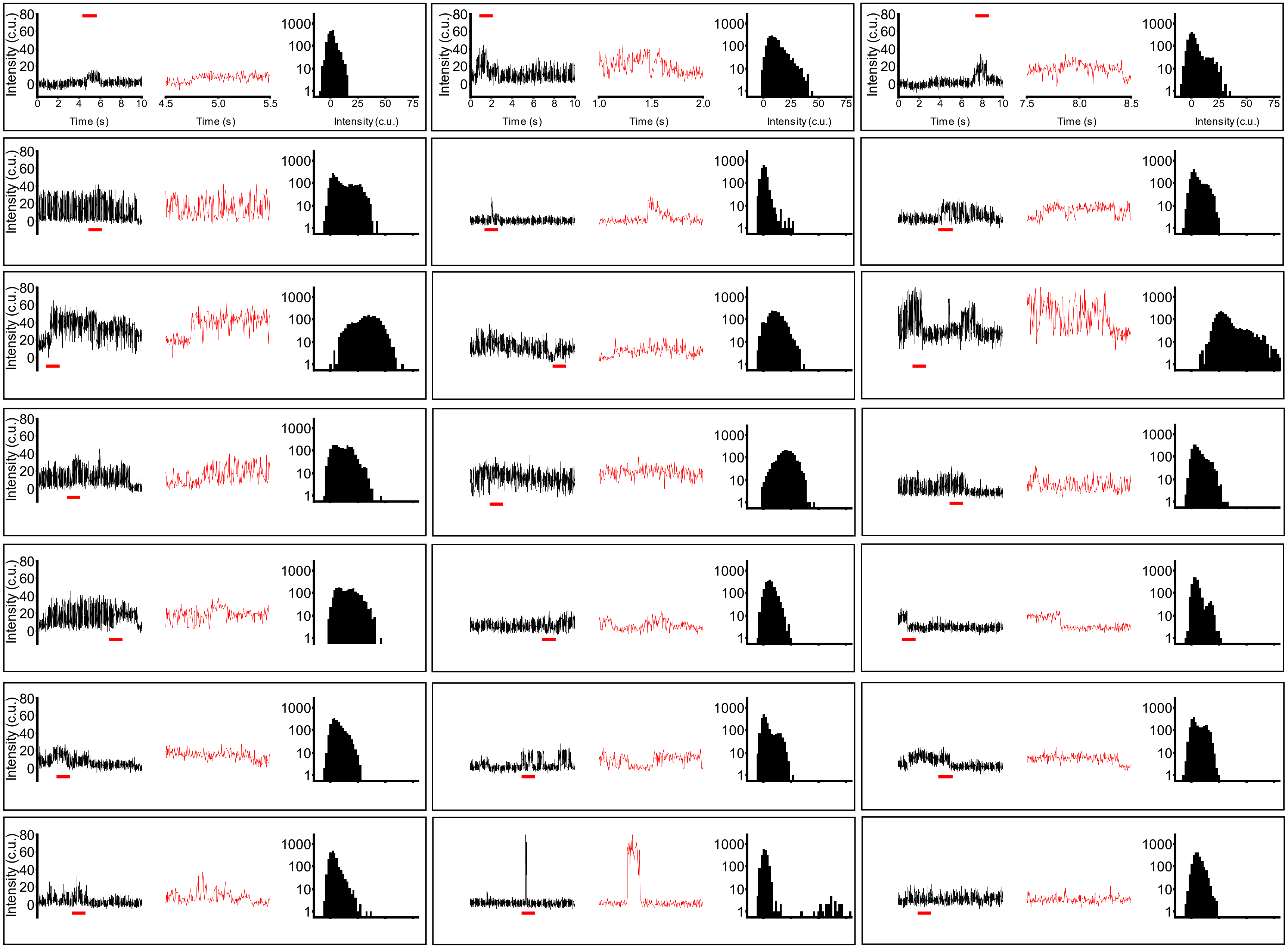

**Supplemental Figure 15. Representative 200-fps activity traces from PIEZO1-HaloTag endothelial cells in hypotonic solution (251 mOsm/L).** Representative background-subtracted fluorescence intensity traces of 21 immobile PIEZO1-HaloTag puncta from TIRF imaging of hiPSC-derived endothelial cells labeled with the $Ca^{2+}$-sensitive JF646-BAPTA HTL. Intensity profiles were plotted from tracked immobile puncta. All traces shown are 10 seconds long (black) with a zoom in of a 1-second portion (red). Right-most panels show an all-points histogram of intensity levels for the entirety of the 10-second recording.

Density scatter plots of PIEZO1-HaloTag localizations

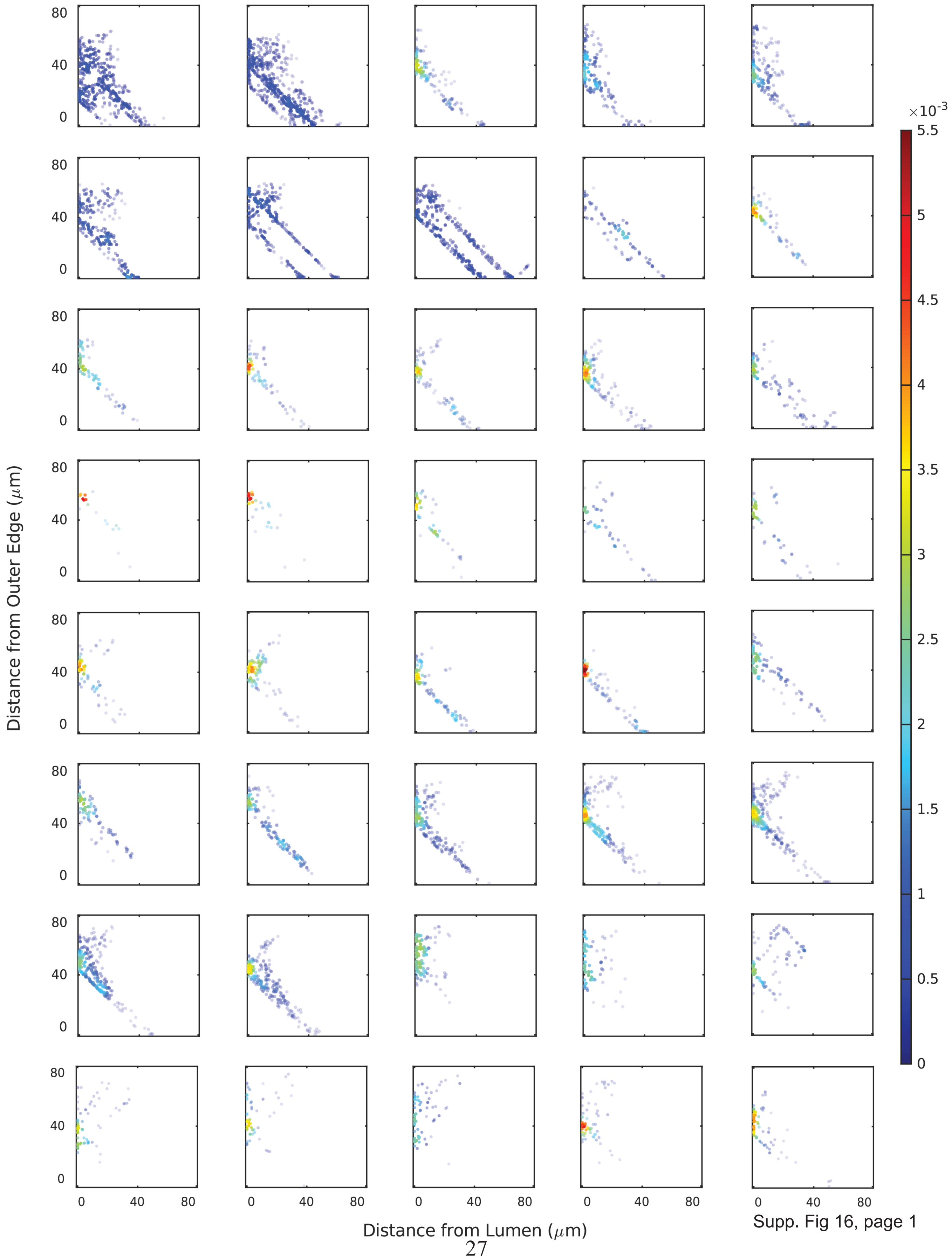


Supp. Fig 16, page 1

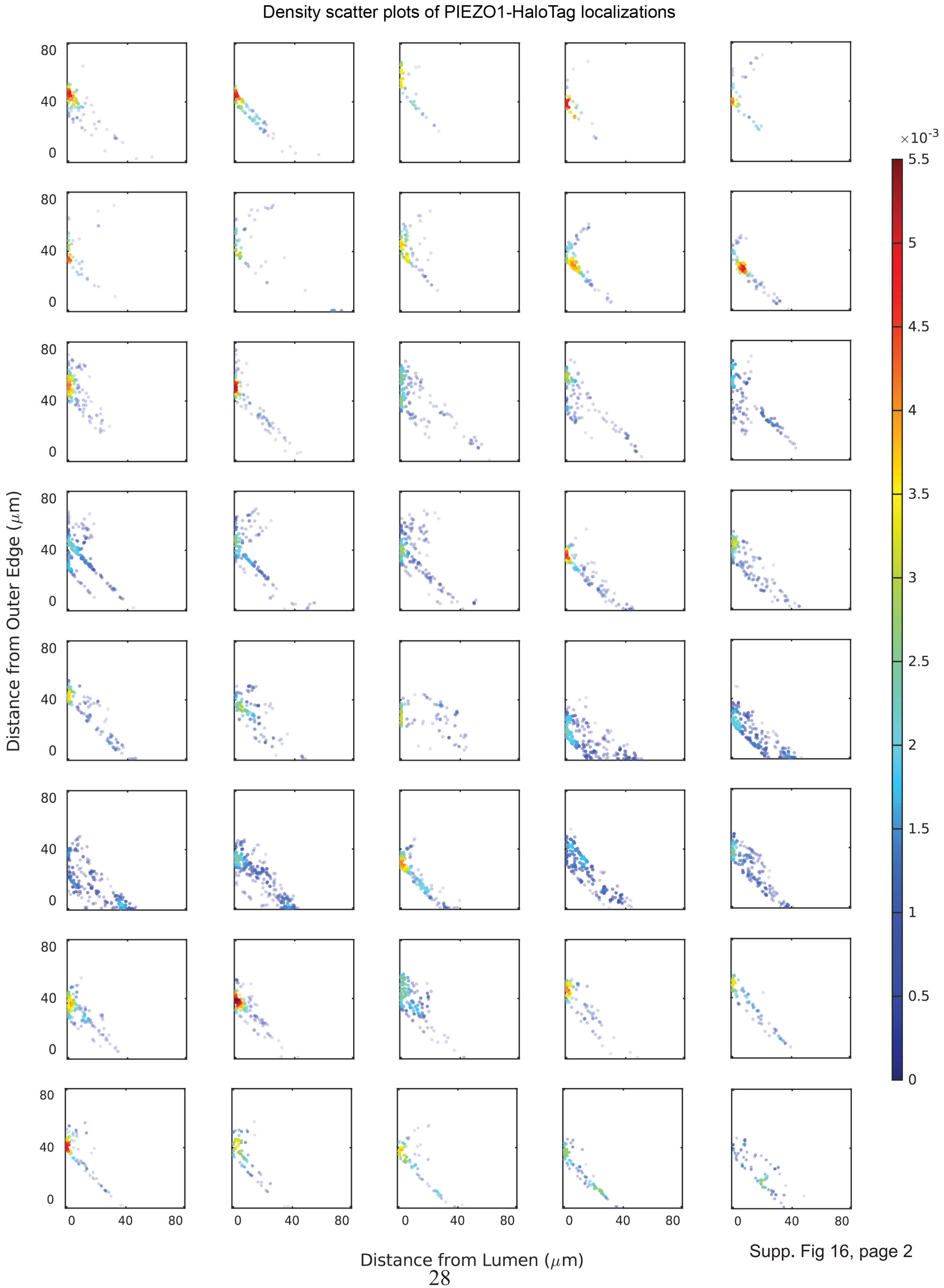


Supp. Fig 16, page 2

Density scatter plots of PIEZO1-HaloTag localizations

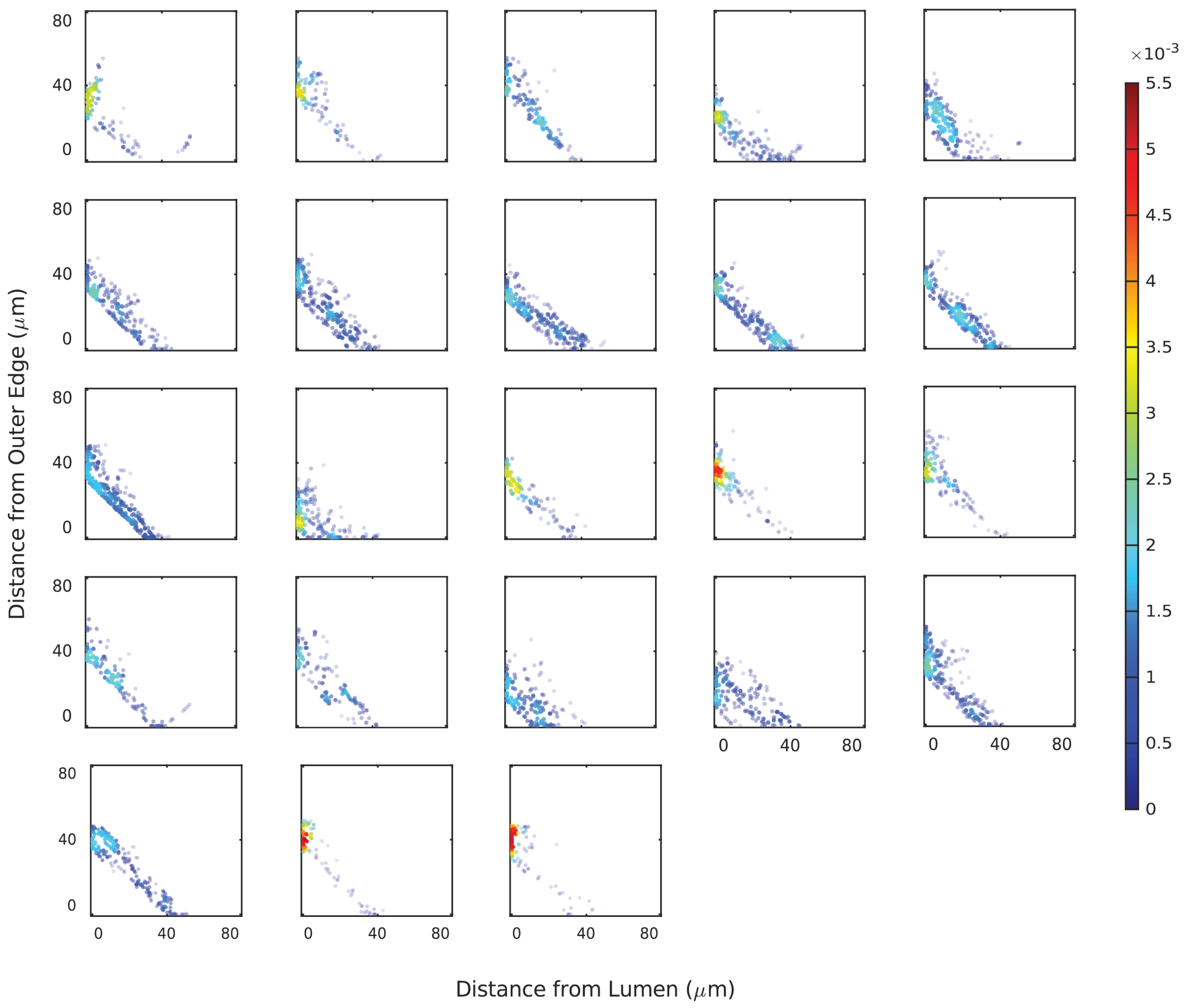

**Supplemental Figure 16. Density scatter plots for individual MNRs labeled with JF635 HTL**. The color scale indicates the relative density of puncta at each position in the scatter plot, normalized to the total number of puncta represented in the plot. n = 103 videos from 21 MNRs from 4 experiments.

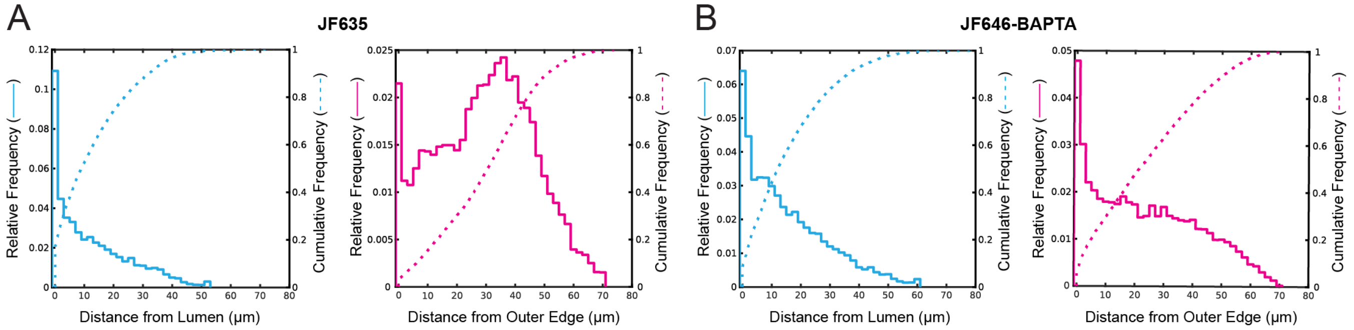


**Supplemental Figure 17. Relative frequency plots for JF635 or JF646-BAPTA labeled MNRs. A.** Relative frequency (solid lines, left axis) and cumulative frequency (dashed lines, right axis) of puncta distances to the lumen border (cyan) and outer edge (magenta) for JF635 labeled PIEZO1-HaloTag MNRs (n = 103 videos from 21 MNRs from 4 experiments). **B.** The same as A, for JF646-BAPTA labeled MNRs (n = 39 videos from 12 MNRs). Cumulative frequency plots for each individual MNR can be found in Supplemental Fig. 18.

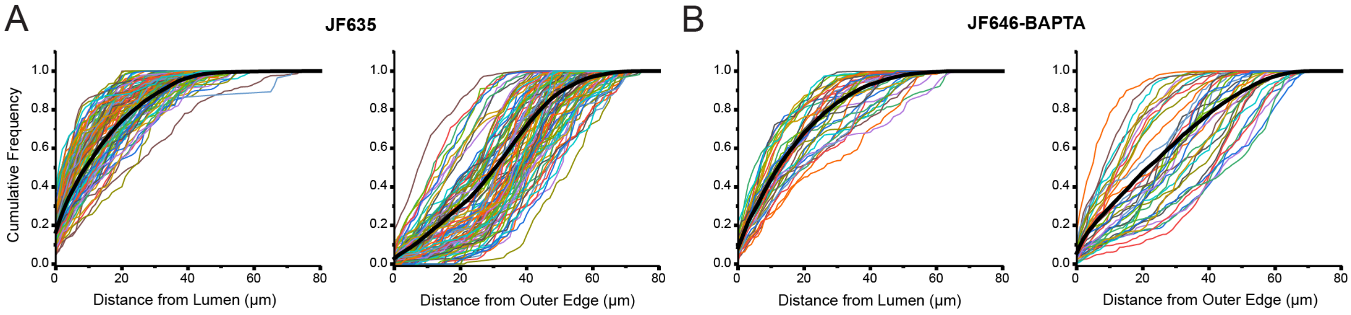


**Supplemental Figure 18. Cumulative frequency plots for individual experiments of JF635 and JF646-BAPTA labeled MNRs. A.** Left: Cumulative frequency plots showing distance of PIEZO1-HaloTag puncta from the MNR lumen border (left) or outer edge (right) for each individually imaged JF635 labeled MNR (n = 103 videos from 21 MNRs from 4 experiments). Each color line denotes a different imaging video. The thick black line indicates the mean cumulative frequency for all experiments. **B.** As for A but for active PIEZO1-HaloTag puncta labeled with JF646-BAPTA (n = 39 videos from 12 MNRs from 3 experiments).

Density scatter plots of active PIEZO1-HaloTag localizations

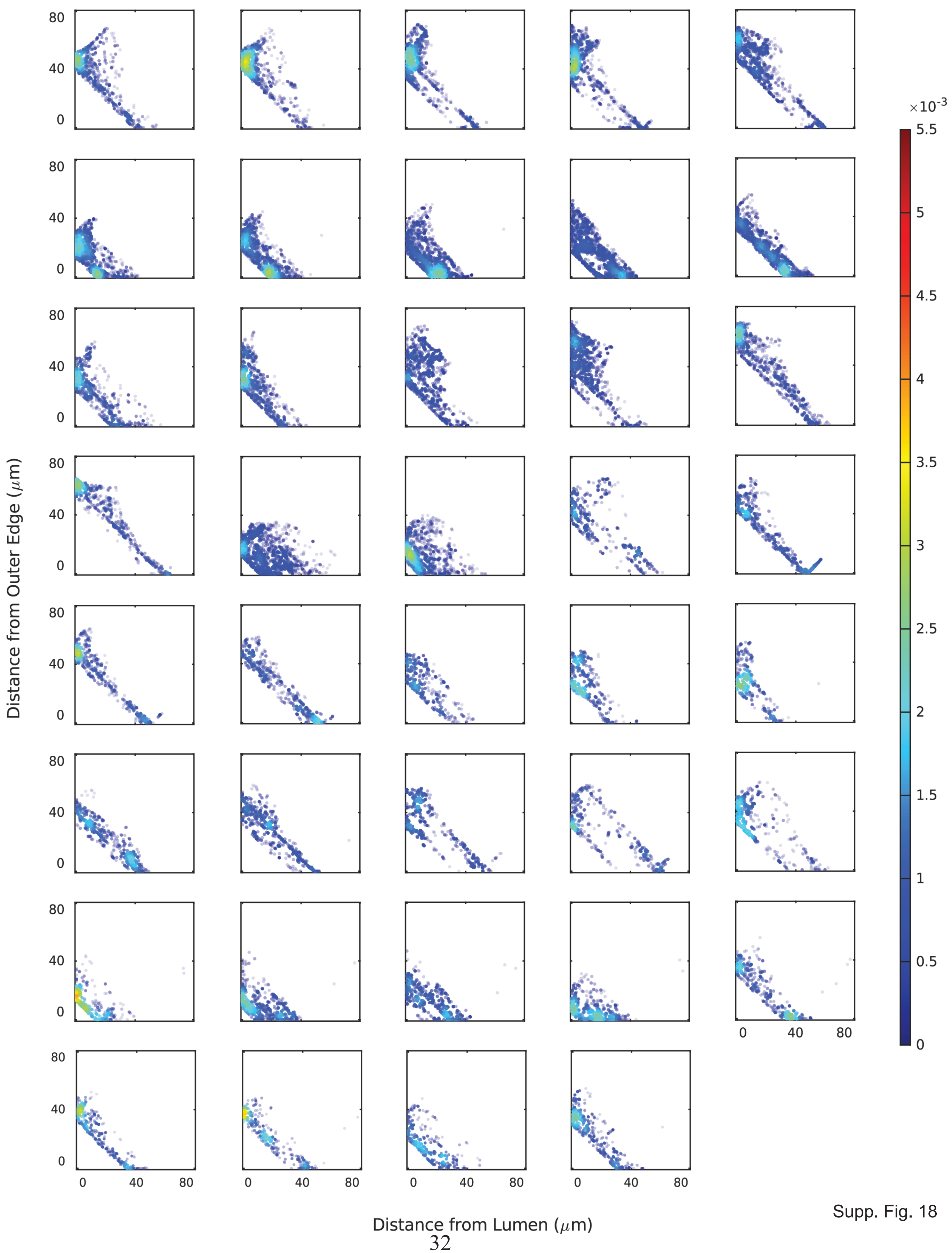


Supp. Fig. 18

**Supplemental Figure 19. Density scatter plots for individual 140-µm MNRs labeled with JF646-BAPTA HTL.** The color scale indicates the relative density of of puncta at each position in the scatter plot, normalized to the total number of puncta represented in the plot. n = 39 videos from 12 MNRs from 3 experiments.

#### SupplementaryVideos

**Supplemental Video 1. PIEZO1 puncta imaged in hiPSC-derived PIEZO1-HaloTag endothelial cells.** A representative PIEZO1-HaloTag endothelial cell labeled with the JF646 HTL and imaged using TIRF microscopy for a duration of 2 minutes at 10 fps.

**Supplemental Video 2. PIEZO1 puncta imaged in hiPSC-derived PIEZO1-HaloTag keratinocytes.** A representative PIEZO1-HaloTag keratinocyte labeled with the JF646 HTL and imaged using TIRF microscopy for a duration of 2 minutes at 10 fps.

**Supplemental Video 3. PIEZO1 puncta imaged in hiPSC-derived PIEZO1-HaloTag neural stem cells.** A representative PIEZO1-HaloTag neural stem cell labeled with the JF646 HTL and imaged using TIRF microscopy for a duration of 2 minutes at 10 fps.

**Supplemental Video 4. Localized enrichment of PIEZO1 at the rear of migrating NSC.** A representative migrating PIEZO1-HaloTag neural stem cell labeled with the JF646 HTL and imaged using TIRF microscopy for 2 minutes at 10 fps.

**Supplemental Video 5. Tracking of PIEZO1-HaloTag endothelial cells.** PIEZO1-HaloTag endothelial cell imaged at 10 fps for 10 seconds. Trajectories were extracted using the protocol detailed in the Methods section, "Analysis of PIEZO1-HaloTag Puncta Diffusion Based on TIRF Imaging."

**Supplemental Video 6. PIEZO1-HaloTag endothelial cells labeled with JF646-BAPTA HaloTag ligand.** TIRF images of JF646-BAPTA labeled PIEZO1-HaloTag and PIEZO1-HaloTag Knockout endothelial cells. Rightmost video shows a PIEZO1-HaloTag endothelial cell treated with 2 μM Yoda1. Videos were acquired at a frame rate of 100 fps.

**Supplemental Video 7. PIEZO1-HaloTag endothelial cells labeled with JF646-BAPTA HaloTag ligand.** TIRF images of JF646-BAPTA labeled PIEZO1-HaloTag and PIEZO1-HaloTag Knockout endothelial cells. Rightmost video shows a PIEZO1-HaloTag endothelial cell treated with 2 μM Yoda1. Videos were acquired at a frame rate of 100 fps.

**Supplemental Video 8. Flickers from a trapped 2 μM of Yoda1 treated PIEZO1-HaloTag JF646-BAPTA puncta.**TIRF video a trapped punctum from a PIEZO1-HaloTag endothelial cell (left) treated with 2 μM of Yoda1. Green dot indicates centroid of trajectory and green box indicates 3x3 pixel region used to generate the fluorescence intensity trace populating on the right. Data were acquired at a frame rate of 200 fps

**Supplemental Video 9. Simultaneous imaging of PIEZO1-HaloTag JF646-BAPTA puncta mobility and activity.** TIRF video of PIEZO1-HaloTag endothelial mobile puncta (left). Green dot indicates centroid of trajectory and the green line shows the trajectory of the puncta movement used to generate the fluorescence intensity trace populating on the right. Data were acquired at a frame rate of 200 fps.

**Supplemental Video 10. 3D Visualization of actin and PIEZO1-HaloTag in MNR.** PIEZO1-HaloTag in MNRs labeled with JF635 HTL imaged using AO-LLSM and visualized using both 2D orthoslices and 3D volumetric views. The video shows actin (white), denoised nuclei (cyan), PIEZO1-HaloTag (green, raw data; magenta, computationally detected PIEZO1 puncta), followed by a 3D local density map of the PIEZO1-HaloTag detections color-coded for the total number of detected PIEZO1-HaloTag puncta within a 3.6 µm search radius or 200 µm3 search volume centered around each detected PIEZO1 puncta.

#### Supplementary Results

##### A size- and time-based detection filtering method removed spurious puncta detections in JF635-labeled MNRs

Lightsheet imaging of PIEZO1-HaloTag in micropatterned neural rosettes (MNRs) allowed us to study PIEZO1 localization and activity in an in vitro model of human neural development. Specifically, we used this system to determine the localization of PIEZO1-HaloTag puncta relative to the lumen border or the outer edge of the MNR (see main Figure 5D-G).

To study puncta localization, we first used the ThunderSTORM detection algorithm (Ovesný et al. 2014) to detect puncta in JF635-labeled PIEZO1-HaloTag MNRs; JF635-labeled PIEZO1-HaloTag KO MNRs as well as unlabeled PIEZO1-HaloTag MNRs served as controls (see Methods). PIEZO1-HaloTag MNRs displayed 1.9 times more detections per MNR than PIEZO1-HaloTag KO MNRs, and a similar number of detections as unlabeled PIEZO1-HaloTag MNRs (calculated from the median values of each condition, see Supplemental Fig. 20A). However, many of the detected puncta were attributed to spurious detections, which were common across all three conditions. We identified three types of such spurious detections: detections within autofluorescence spots, detections due to unbound JF ligand remaining after wash steps, and false positive detections within the ThunderSTORM detection algorithm. Therefore, we developed a method to computationally filter out these spurious detections and enrich the pool of detections representing bona fide PIEZO1-HaloTag puncta.

We observed spurious detections within large autofluorescence spots, i.e. spots also present in control KO and unlabeled MNRs (Supplemental Fig. 20B-D). These spots were immobile, presented stable intensity profiles over time (except for a progressive intensity decrease due to photobleaching), and were much larger than PIEZO1-HaloTag puncta. Therefore, we applied a first filtering step based on the size of the detected puncta, in which we removed any initial detections that had a standard deviation (**σ**) of their Gaussian fit larger than 280 nm. This step successfully removed most detections that were located within autofluorescence spots (Supplemental Fig. 20B-D). After size-based filtering, there were 2.8 times more detections retained in JF635 PIEZO1-HaloTag MNRs than in JF635 PIEZO1-HaloTag KO MNRs; and 3.6 times more than in unlabeled PIEZO1-HaloTag MNRs (Supplemental Fig. 20E). Therefore, the size-based filtering step already removed a large number of spurious detections.

Besides detections in autofluorescence spots, we observed spurious detections corresponding either to unbound JF635 HTL or to small local background fluctuations being detected as puncta (Supplemental Fig. 20B-D). These detections generally appeared for only one or two frames, while bona fide PIEZO1-HaloTag puncta observed in JF635 PIEZO1-HaloTag MNRs tended to last several frames (Supplemental Fig. 20F). We thus used the FLIKA algorithm to track puncta over time and then applied a second, time-based filtering step designed to remove spurious detections that lasted only for one or two frames of imaging (Supplemental Fig. 20B-D).

After these two filtering steps, JF635 PIEZO1-HaloTag MNRs contained 9.6 times more puncta than JF635 PIEZO1-HaloTag KO MNRs, and 6.9 times more than unlabeled PIEZO1-HaloTag MNRs (Supplemental Fig. 20G). Therefore, our puncta filtering method allowed us to remove detections that were not representative of bona fide PIEZO1 puncta, and to retain detections representative of PIEZO1 puncta for further analysis.

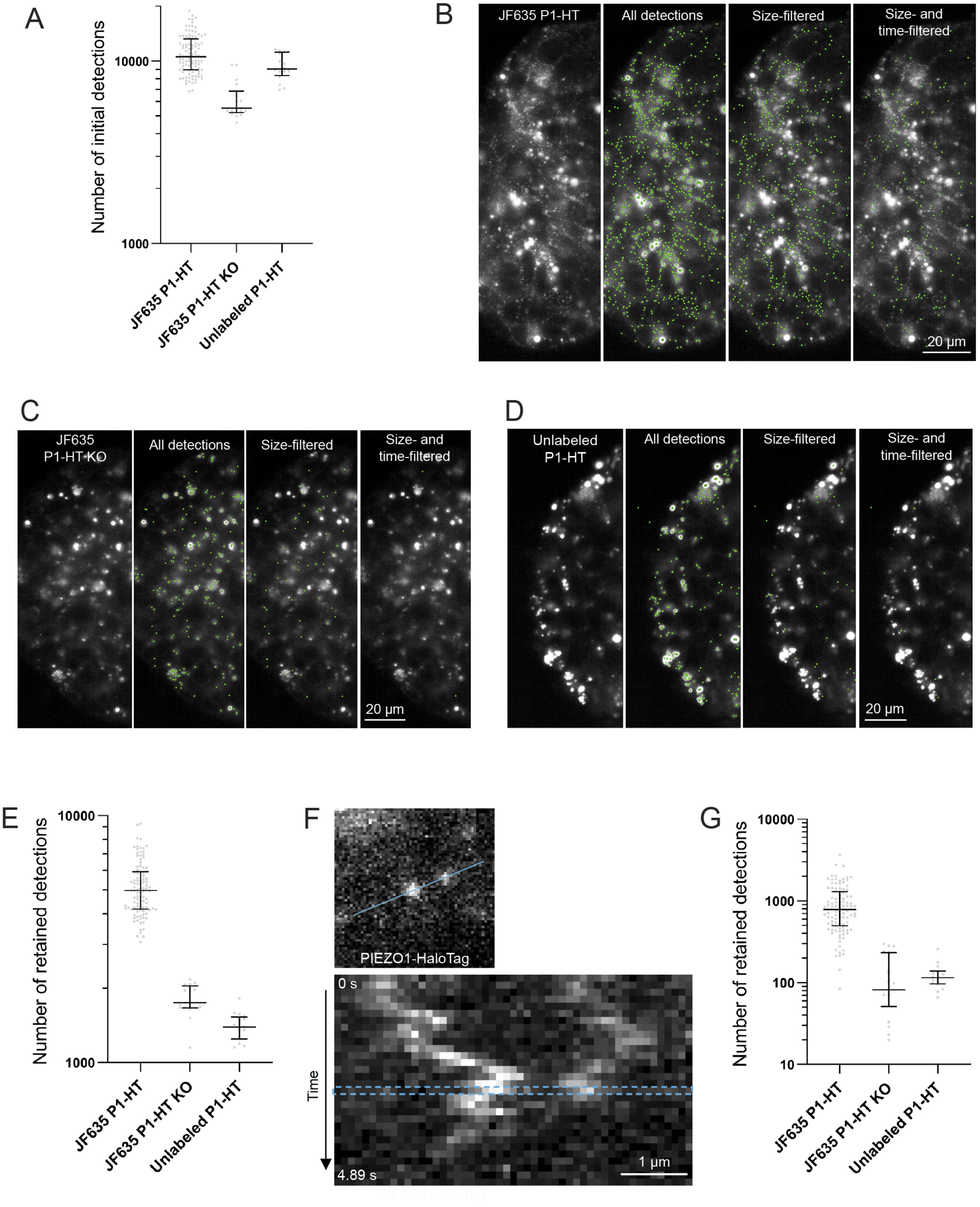
A
Number of initial detections
10000
1000
JF635 P1-HT
JF635 P1-HT KO
Unlabeled P1-HT
B
JF635 P1-HT
All detections
Size-filtered
Size- and time-filtered
20 µm
C
JF635 P1-HT KO
All detections
Size-filtered
Size- and time-filtered
20 µm
D
Unlabeled P1-HT
All detections
Size-filtered
Size- and time-filtered
20 µm
E
Number of retained detections
10000
1000
JF635 P1-HT
JF635 P1-HT KO
Unlabeled P1-HT
F
PIEZO1-HaloTag
0 s
Time
4.89 s
1 µm
G
Number of retained detections
10000
1000
100
10
JF635 P1-HT
JF635 P1-HT KO
Unlabeled P1-HT

**Supplemental Figure 20. Size- and time-based filtering removes spurious localizations in JF635-labeled MNRs. A.** Number of puncta detected by ThunderSTORM in each sample type. Median values: JF635 PIEZO1-HT: 10579 detections, JF635 PIEZO1-HT KO: 5528 detections, unlabeled PIEZO1-HT: 9046 detections. **B.** Representative images of puncta detections remaining after each step in the puncta filtering process in the first data frame of a JF635 PIEZO1-HaloTag MNR. From left to right: PIEZO1-HaloTag data, initially detected puncta, remaining puncta after size-based filtering, and remaining puncta after time-based filtering. Green dots denote the detected puncta. **C.** As in B, but for a JF635 PIEZO1-HaloTag Knockout MNR. **D.** As in B and C, but for an unlabeled PIEZO1-HaloTag MNR. **E.** Number of remaining puncta in each sample type after size-based filtering. Median values: JF635 PIEZO1-HT: 4975 detections, JF635 PIEZO1-HT KO: 1746 detections, Unlabeled PIEZO1-HT: 1393 detections. **F.** Representative 3-plane stack image of JF635 PIEZO1-HaloTag puncta in an MNR. The blue line indicates the region of interest used to generate the kymograph on the right. The dashed blue rectangle in the kymograph indicates the frame shown in the left panel. Note how JF635 PIEZO1-HaloTag puncta remain bright over several frames, in contrast to the similar kymograph shown in Fig. 4F for JF646-BAPTA HTL which displays flickering. **G.** Number of remaining puncta in each sample type after size- and time-based filtering. The two-step filtering process resulted in a substantial reduction of detections in JF635 PIEZO1-HT KO MNRs and unlabeled PIEZO1-HaloTag MNRs, with a smaller reduction of puncta in JF635 PIEZO1-HaloTag MNRs. Median values: JF635 PIEZO1-HT: 788 detections, JF635 PIEZO1-HT KO: 82 detections, Unlabeled PIEZO1-HT: 115 detections. All plots: Each dot is one video, bars are the median with interquartile range for each condition. JF635 PIEZO1-HaloTag: n=103 videos from 21 MNRs from 4 experiments; JF635 PIEZO1-HaloTag KO: n=19 videos from 6 MNRs from 3 experiments; unlabeled PIEZO1-HaloTag: n=16 videos from 2 MNRs from 1 experiment.

**A size- and intensity-based detection filtering method removed spurious puncta detections in JF646-BAPTA-labeled NMRs**

Light-sheet imaging of MNRs labeled with JF646-BAPTA HTL also enabled us to quantify the spatial distribution of active puncta in the neural organoids. Before analyzing the data, we conducted two filtering steps to remove spurious detections. The first step was identical to the size-based step used for JF635 samples above, while the second step was tailored to the highly variable intensity profiles displayed by BAPTA HTL ligands.

Upon initial puncta detection with ThunderSTORM, JF646-BAPTA PIEZO1-HaloTag MNRs presented 1.6 times more detections than JF646-BAPTA PIEZO1-HaloTag KO MNRs, and a similar number of detections as unlabeled PIEZO1-HaloTag MNRs (Supplemental Fig. 21A). Many of these arose from sample "autofluorescent spots" independent of PIEZO1. As above, we filtered out these out by excluding puncta with **σ** over 280 nm (Supplemental Fig. 21B-D). After this size-based filtering, there were 1.5 times more detections retained in JF636-BAPTA PIEZO1-HaloTag MNRs than in JF646-BAPTA PIEZO1-HaloTag KO MNRs; and 1.8 times more than in unlabeled PIEZO1-HaloTag MNRs (Supplemental Fig. 21E). Therefore, the size-based filtering step removed a large number of spurious initial detections from all three conditions, and in particular from unlabeled PIEZO1-HaloTag MNRs.

Because BAPTA probes flicker quickly over time (see kymograph in main Fig. 5F), the second filtering step was based on puncta integrated intensity (the sum of intensity values for all pixels under the Gaussian curve fitted to each detected puncta) rather than track duration (Supplemental Fig. 21B-D). Here we removed puncta with a value of integrated intensity lower than ~77 photons (corresponding to 261 c.u.), based on the distribution of integrated intensity values retained after the size-based filtering step (Supplemental Fig. 21F). Relative frequency histograms of integrated intensity values showed large proportions of puncta contained under the 77-photon threshold, in particular for negative control samples. After filtering these spurious puncta out based on this intensity threshold, JF646-BAPTA PIEZO1-HaloTag MNRs had 2.6 times more localizations than JF646-BAPTA PIEZO1-HaloTag KO MNRs, and 2 times more than unlabeled PIEZO1-HaloTag MNRs (Supplemental Fig. 21G). Only detections that were both small enough in spatial spread and bright enough to represent active PIEZO1 puncta were then retained for analysis of active punta localization in MNRs. Therefore, while this approach did not remove as many spurious detections as the time-based filtering applied to JF635-labeled samples above, intensity-based filtering also successfully retained bona fide active PIEZO1-HaloTag puncta.

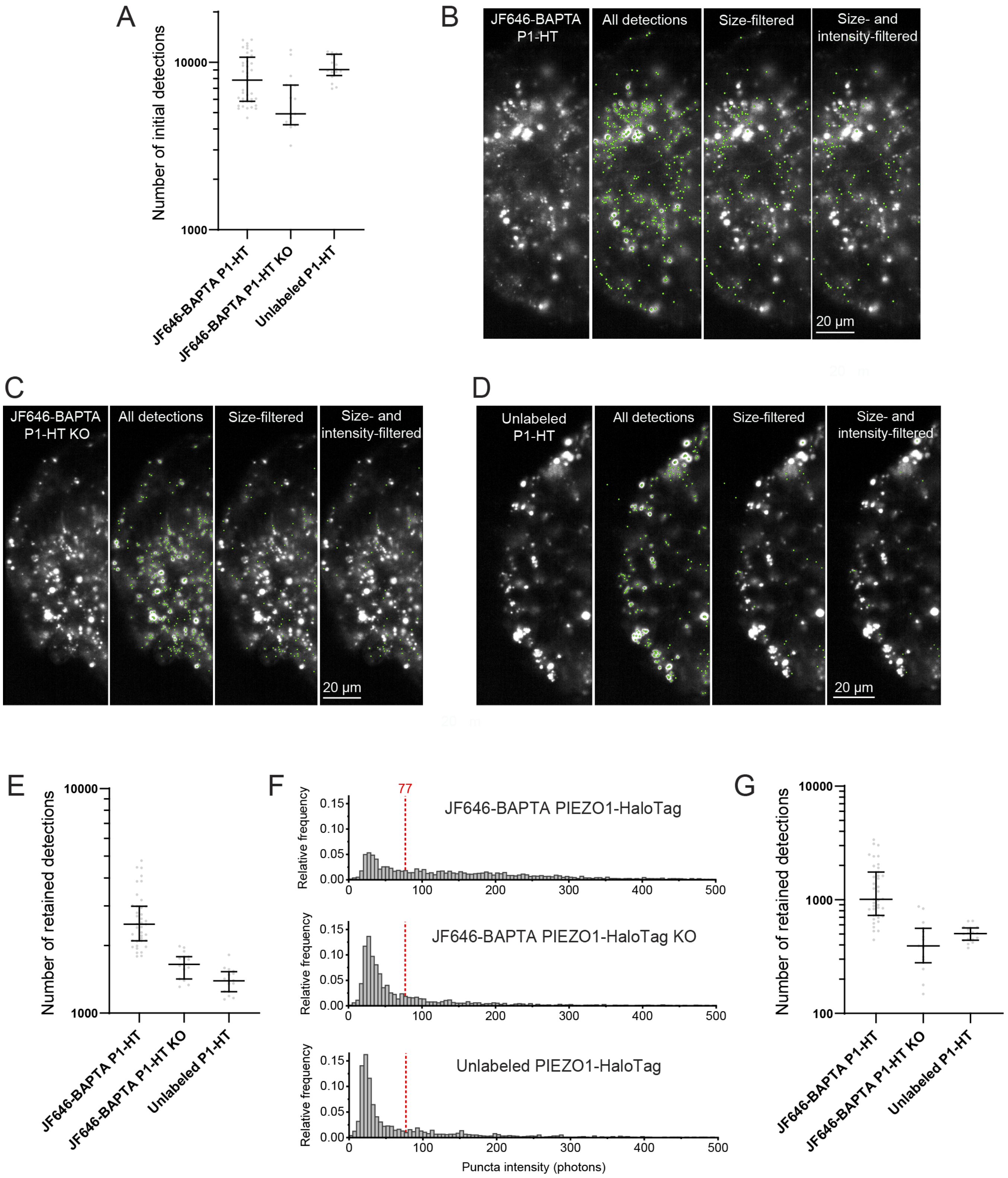
A
Number of initial detections
10000
1000
JF646-BAPTA P1-HT
JF646-BAPTA P1-HT KO
Unlabeled P1-HT
B
JF646-BAPTA P1-HT
All detections
Size-filtered
Size- and intensity-filtered
20 μm
C
JF646-BAPTA P1-HT KO
All detections
Size-filtered
Size- and intensity-filtered
20 μm
D
Unlabeled P1-HT
All detections
Size-filtered
Size- and intensity-filtered
20 μm
E
Number of retained detections
10000
1000
JF646-BAPTA P1-HT
JF646-BAPTA P1-HT KO
Unlabeled P1-HT
F
77
JF646-BAPTA PIEZO1-HaloTag
JF646-BAPTA PIEZO1-HaloTag KO
Unlabeled PIEZO1-HaloTag
Relative frequency
0.15
0.10
0.05
0.00
0
100
200
300
400
500
Puncta intensity (photons)
G
Number of retained detections
10000
1000
100
JF646-BAPTA P1-HT
JF646-BAPTA P1-HT KO
Unlabeled P1-HT

**Supplemental Figure 21. Size- and intensity-based filtering removes spurious localizations in JF646-BAPTA-labeled MNRs. A.** Number of puncta initially detected by ThunderSTORM in each sample type. Median values: JF646-BAPTA PIEZO1-HT: 7834 detections, JF646-BAPTA PIEZO1-HT KO: 4928 detections, Unlabeled PIEZO1-HT: 9046 detections. **B.** Representative images of puncta detections remaining after each step in the puncta filtering process in the first data frame of a JF646-BAPTA PIEZO1-HaloTag MNR. From left to right: PIEZO1-HaloTag data, initially detected puncta, remaining puncta after size-based filtering, and remaining puncta after intensity-based filtering. Green dots denote the detected puncta. **C.** As in B, but for a JF646-BAPTA PIEZO1-HaloTag Knockout MNR. **D.** As in B and C, but for an unlabeled PIEZO1-HaloTag MNR. **E.** Number of remaining puncta in each sample type after size-based filtering. Median values: JF635 PIEZO1-HT: 2491 detections, JF635 PIEZO1-HT KO: 1650 detections, Unlabeled PIEZO1-HT: 1393 detections. **F.** Histograms of integrated intensity values (the sum of intensity values for all pixels under the Gaussian curve fitted to each detected puncta) for all puncta detected in a representative sample of each condition after the size-based filtering step. Based on intensity profiles, we selected a threshold of 77 photons (shown in a dashed red line), which removed the large amounts of detections under that threshold, particularly in control samples, while preserving bona fide JF646 signal. **G.** Number of remaining puncta in each sample type after size- and intensity-based filtering. The two-step filtering process resulted in a substantial reduction of detections in JF646-BAPTA PIEZO1-HT KO MNRs and unlabeled PIEZO1-HaloTag MNRs, with a smaller reduction of puncta in JF646-BAPTA PIEZO1-HaloTag MNRs. Median values: JF635 PIEZO1-HT: 1009 detections, JF635 PIEZO1-HT KO: 393 detections, Unlabeled PIEZO1-HT: 504 detections. Plots in A, E, G: Each dot is one video, bars are the median with interquartile range. JF646-BAPTA PIEZO1-HaloTag: n=39 videos from 12 MNRs from 3 experiments; JF646-BAPTA PIEZO1-HaloTag KO: n=15 videos from 3 MNRs from 3 experiments; unlabeled PIEZO1-HaloTag: n=16 videos from 2 MNRs from 1 experiment.
